# RBPamp: Quantitative Modeling of Protein-RNA Interactions *in vitro* Predicts *in vivo* Binding

**DOI:** 10.1101/2022.11.08.515616

**Authors:** Marvin Jens, Michael McGurk, Ralf Bundschuh, Christopher B. Burge

## Abstract

RNA-binding proteins (RBPs) control the processing and function of cellular transcripts to effect post-transcriptional gene regulation. Sequence-specific binding of RBPs to millions of synthetic RNAs has been probed in vitro by RNA Bind-n-Seq (RBNS). Here we describe RBPamp, a bio-physically-based model of protein-RNA interactions and associated algorithm that inferred affinity spectra of 79 diverse human RBPs from RBNS data. RBPamp supports multiple motifs per RBP, models RBP concentration and binding site saturation, and accounts for the effects of RNA secondary structure. RBPamp affinities along transcripts are predictive of in vivo binding, as measured by eCLIP density. For many RBPs, average local eCLIP density increases monotonically with predicted affinity, and the shape of this relationship can suggest free protein concentrations and potential cooperativity. Together, these analyses demonstrate a powerful integrative approach for the quantitative dissection of RBP function.

## Introduction

RNA transcripts from eukaryotic genes undergo complex processing and regulation to control protein production and function. These co- and post-transcriptional processes, which are critical for organismal development (Becker et al., 2018; Grün et al., 2014; Lennox et al., 2018; Liu et al., 2016; Weil, 2015) and underlie many human diseases (Castello et al., 2013; Lukong et al., 2008), are coordinated by hundreds of RNA-binding proteins (RBPs) (Bartel, 2009; Gerstberger et al., 2014; Keene, 2007). Many RBPs function by recruiting or blocking other factors that ultimately affect the splicing, localization, stability, or translation of RNAs (Corley et al., 2020, 2020; Lunde et al., 2007), implying that occupancies are strongly related to activity. Despite recent progress (Alipanahi et al., 2015; Jolma et al., 2020; Ray et al., 2013; Van Nostrand et al., 2020), our ability to predict *in vivo* occupancy by RBPs across the transcriptome remains limited. Many current computational approaches treat *in vivo* binding as a classification problem (Li et al., 2010; Maticzka et al., 2014) or use dimensionless binding scores (Alipanahi et al., 2015) which are not directly analogous to physical binding affinities. On the other hand, biophysical approaches have been successfully employed to model transcription factor binding to DNA (Rastogi et al., 2018; Rube et al., 2022).

Assuming that RBP binding is similarly driven by the intrinsic affinity of individual proteins (Van Nostrand et al., 2020), one can use quantitative measurements of protein-RNA inteaction *in vitro* to address this issue. Available high-resolution methods include RNAcompete (Ray et al., 2009), RNA-Mitomi (Martin et al., 2012), RNA-MaP (Buenrostro et al., 2014) and RNA Bind-n-Seq (RBNS) (Lambert et al., 2014). While RNA-Mitomi and RNA-MaP directly measure binding to many sequences and can yield accurate affinities, they are experimentally challenging, difficult to scale to a large number of RBPs, and restrict the number of assayed sequences to 640 (Mitomi) or tens of thousands (RNA-MaP), which may not be sufficient to fully characterize binding motifs or the impact of RNA structure. RBNS is a comparatively straightforward *in vitro* assay similar to the DNA-binding method HT-SELEX (Zhao et al., 2009), in which tagged, recombinantly expressed RBP is mixed with an extremely diverse pool of synthetic random RNAs (exceeding the diversity of natural RNAs in the cell) at defined concentrations. After an equilibration period, RBP-bound RNA is isolated, followed by sequencing at depths of ∼10^7^ reads for both bound and input RNA, enabling comprehensive and reasonably quantitative assessment of binding across a wide spectrum of sequences. RNACompete is a related method, typically involving a designed pool of RNAs and assessment of binding by microarray, though a sequencing-based method has been introduced recently (Cook et al., 2017). To date, RNAcompete and RBNS have been applied to the largest numbers of RBPs.

Several features of RBPs complicate the analysis of data from RBNS and related methods, including three that are particularly challenging. First, RBPs frequently contain multiple RNA-binding domains (RBDs) and may interact with RNA in different modes that are not representable by a single position-specific frequency/weight matrix (PFM/PWM), which assumes a fixed “register” across motif positions (Berg and von Hippel, 1987). For example, RBPs often tolerate variable spacing between distinct subparts of the motif, or may tolerate a base that is “flipped out” relative to the bases contacted by the protein, both leading to recognition elements of different lengths (Afroz et al., 2015; Dominguez et al., 2018; Valley et al., 2012). Second, mass-action effects may complicate the relationship between affinity and site occupancy: depending on the relative concentrations used, the RBP may be titrated by large amounts of RNA, or binding sites of higher affinity may become saturated. Finally, as single-stranded RNA (ssRNA) is the preferred substrate for most RBPs (Lunde et al., 2007), the formation of RNA secondary and tertiary structures can substantially impair binding (Forties and Bundschuh, 2010; Jarmoskaite et al., 2019; Li et al., 2010; Wan et al., 2011).

Here, we employ a biophysically-motivated framework tailored to RNA-binding, and an associated algorithm to address these issues, which we call RBP affinity model with physical constraints (RBPamp). We were able to fit this model to RBNS data for 79 RBPs by computational optimization (Fig. 1a), yielding quantitative descriptions of RBP affinity. Integrative modeling of affinity and eCLIP data revealed aspects of RBP biology across the transcriptome.

**Fig. 1.**
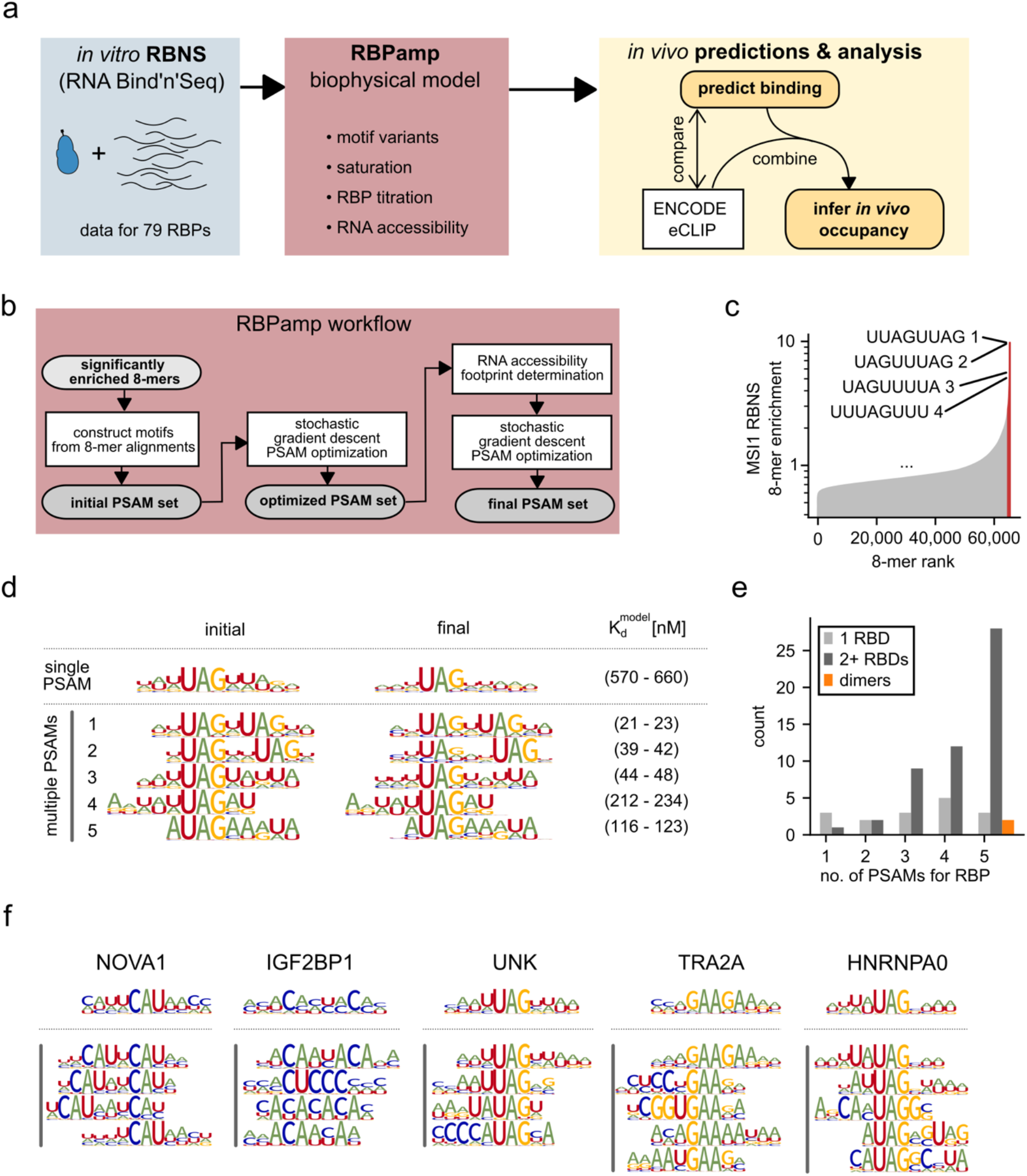
Multiple PSAMs capture details of RBP binding. **a)** Overview of this study: RBPamp (red) builds on *in vitro* RBNS data (blue). Biophysical models of RBP-binding allow comparison to and novel analyses (orange) of *in vivo* data (white boxes). **b)** Steps in the RBPamp algorithm. **c)** RNA 8-mers ranked by RBNS enrichment (y-axis) for the Musashi1 (MSI1) RBP. 8-mers with z-score > 4 are shown in red. 8-mers used to seed PSAMs are listed with PSAM number. **d)** Visual representations of initial PSAMs (left) and final PSAMs (middle) for MSI1, with 5%-95% confidence intervals of estimated dissociation constants for cognate sequences (*K*_*d*_^*model*^), for final PSAMs (right). Results of a RBPamp run restricted to a single PSAM (top) and results of a standard run yielding 5 PSAMs (bottom) are shown. **e)** Histogram of number of PSAMs assigned to RBPs grouped by whether they have single (light gray) or multiple (dark gray) KH, ZNF, or RRM domains, or probably dimerize (orange). RBPs with other or mixed domains were excluded. **f)** Initial PSAMs for selected RBPs. Single (top) vs. multiple (bottom, grouped by vertical bar) PSAMs as in (d). See also Fig. S1 and Table S1.

## Results

### Sets of PSAMs capture different modes of RBP binding

Short sequences of length *k* (“*k*-mers”) that are enriched in the RBNS bound pool versus the input pool are informative about the sequence preferences of an RBP (Lambert et al., 2014). The raw enrichment or “R” value of a *k*-mer is defined as the ratio of its frequency in the pulldown dataset to that in the control set. This reflects selection by the RBP, but is not directly proportional to the binding affinity for the *k*-mer because of the complexities discussed above (site accessibility, titration, saturation), spurious enrichment of overlapping *k*-mers (Lambert et al., 2014), and experiment-specific background. To quantitatively describe protein-RNA interactions, we modeled variable-length primary sequence motifs using position-specific affinity matrices (PSAMs) (Foat et al., 2006; Rube et al., 2022). Each matrix element in a PSAM represents the affinity of a motif containing the specified base relative to that of the matrix’s most preferred (cognate) sequence, with the cognate base at each position assigned an affinity of 1.0. These relative affinities are considered to reflect biophysical properties of the protein-RNA complex and are therefore independent of background nucleotide frequencies, differing from other weight-matrix schemes such as log-odds scores. Unlike Foat and colleagues, we model an RBP’s binding preferences with a mixture of multiple PSAMs to accommodate complex features of RBP binding, such as bipartite motifs. RBPamp merges similar significantly enriched *k*-mers (Fig. S1) to seed initial motifs, generating between one and five PSAMs for each RBP, with subsequent tuning of parameters (Fig. 1b).

Use of multiple PSAMs provides flexibility to capture alternate motifs or motif versions with variation in spacing between core elements, as discussed above. For example, the human RBP MSI1 shows strong enrichment for the 8-mers U**UAG**U**UAG** and **UAG**UU**UAG** (Fig. 1c) composed of two UAG core elements with variable spacing (Iwaoka et al., 2017) (Fig. 1d). The optimization procedure adjusts the parameters of each PSAM to more accurately reflect relative affinities of different sequence variants compared to the “ideal”, cognate sequence by optimizing fit between observed and predicted 6-mer enrichments. Each PSAM is also assigned a value playing the role of the absolute dissociation constant of the cognate (highest-affinity) sequence within the model (*K*_*d*_^*model*^), which is simultaneously optimized (Methods).

For MSI1, this procedure yielded a set of five PSAMs (Fig. 1d). To graphically represent a PSAM, we scale each column by its discrimination (Foat et al., 2006) – the degree to which non-cognate bases are penalized at this position – such that highly specific motif positions are larger, analogous to sequence logos (Schneider and Stephens, 1990). For example, an ideal UAGNNUAG PSAM 1 motif is assigned an affinity 5-to 10-fold higher than the ideal motifs for PSAMs 4 and 5, which feature a single UAG with different flanking sequence preferences.

The number of PSAMs assigned by RBPamp was greater for RBPs with multiple RRM, KH, or zinc finger RBDs compared to single-domain proteins (Fig. 1e, *P* < 0.0024, Mann-Whitney U test; see also Supplementary Table S1), suggesting that the algorithm detects increased binding complexity in data from multi-RBD proteins. Proteins with two or more RBDs were often assigned sets of PSAMs that involve similar pairs of sub-motifs with variable spacing. For example, RBPamp detects tandem repeats of CAY occurring 1-3 nucleotides apart for the triple KH-domain NOVA1 protein (Fig. 1f), consistent with previous studies of NOVA1 (Teplova et al., 2011). Similarly, different spacings of ACA core motifs are recognizable in IGF2BP1 PSAMs 1, 3, and 4, with PSAM 2 representing an alternate C-rich motif. We also observe such alternate motifs for UNK, TRA2A, HNRNPA0 and others. For example, HNRNPA0 PSAMs 1 and 2 favor UA or AU dinucleotides over UU or AA dinucleotides upstream of the central UAG motif, suggesting a dependency between the bases at these two positions. These examples illustrate the richer binding inferences enabled by use of multiple PSAMs.

### Comprehensive modeling of an RBNS experiment

To assess how well the PSAMs explain the observed binding, we employ a biophysically-motivated model of the RBNS experiment, using a framework based on the treatment of complex ligand mixtures (Cantor and Schimmel, 1980) that includes mass-action effects. In the first step, we predict RBP occupancy for a large number of sequences sampled from the random RNA pool as a function of the PSAMs and the total RBP concentration in each RBNS sample. We then describe the RBNS experiment as an RNA-protein mixture that accounts for: (1) binding site saturation (Fig. 2a); (2) titration of the RBP by substrate binding (Fig. 2b); and (3) RNA secondary structural accessibility of binding sites (Fig. 2c). Modeling binding site saturation was motivated by the observation that the pool of random RNA often contains binding sites with affinities spanning multiple orders of magnitude (Fig. 2d), so that high-affinity sites are expected to become saturated at high RBP concentrations, as seen for TRA2A (Fig. 2e). While SPR and RNA-MaP approaches use continuous flow to maintain a constant free protein concentration, RBNS uses a closed system with a fixed amount of protein, necessitating estimation of total bound RBP from the distribution of RNA site affinities across the input RNA pool (Fig. 2f).

**Fig. 2.**
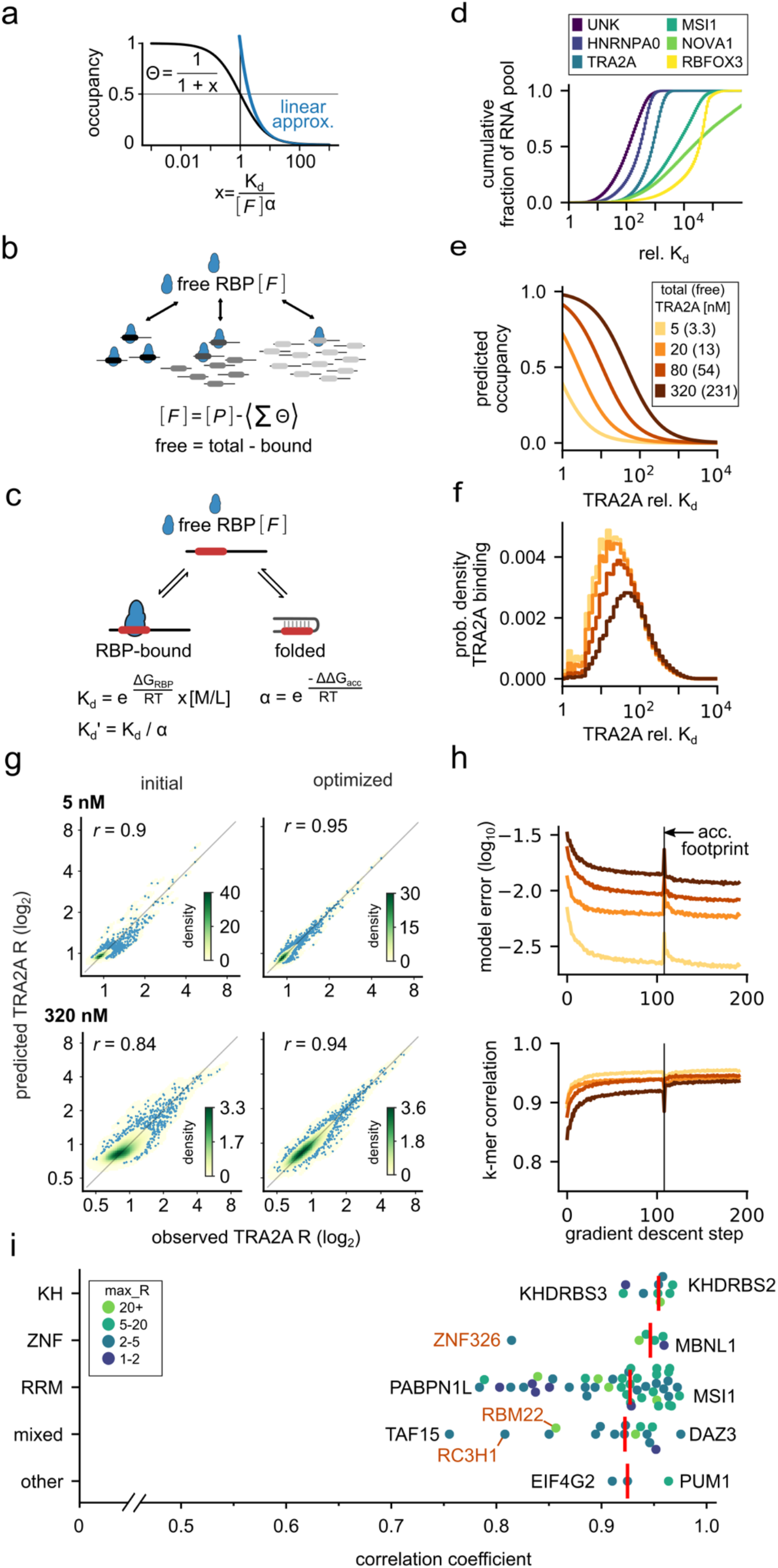
The RBPamp model provides an excellent fit to RBNS data. **a)** Probability of RBP binding (occupancy), Θ, as a function of *x* = K_d_/([F] α). When *x* >> 1, Θ ≈ ([F] α) / K_d_ (linear approximation, blue). When x → 0, Θ → 1 (saturation), use of the general formula is needed (black). **b)** Cartoon illustrating titration of total concentration [P] of an RBP (blue) by the RNA pool (black lines). Ovals indicate RBP motifs, with shade indicating relative affinity (darker = higher, lighter = lower). Titration reduces the concentration of free RBP, so that [F] < [P]. **c)** Cartoon illustration of possible states for a binding site (red) within an RNA molecule (black line) in presence of free RBP (blue): the RNA may be unbound (top), bound (left), or folded (right). The presence of RBP-inaccessible, folded states can be expressed by the apparent dissociation constant K’_d_ = K_d_ / α. **d)** Distribution of binding affinity in the random RNA pool for different RBPs, plotted as cumulative fraction of binding sites as a function of relative affinity, expressed as a multiple of the K_d_ of the highest-affinity motif. RBPs range from low specificity (e.g., UNK) to high specificity (e.g., RBFOX3). **e)** Predicted binding site occupancy plotted as a function of relative affinity (K_d_/K_d_^model^) for different total concentrations of TRA2A and estimated free TRA2A (in parentheses in the key). **f)** Predicted composition of RBP-bound RNA as a function of relative affinity (normalized histogram). **g)** Scatter plots of predicted (y-axis) versus observed (x-axis) 6-mer R-values (log_2_) for low and high TRA2A concentrations (top, bottom), before and after SGD optimization (left, right). Shown is a kernel density estimate (see color bar), beneath a scatter plot (blue dots) of 6-mers in regions of low density. Pearson correlations indicated by “r” are shown. **h)** Squared difference in log-R-values between experiment and prediction (top), and the corresponding log_2_ R-value correlation (bottom) during the course of SGD optimization performed in parallel on samples with four different RBP concentrations (colors as in e,f). Vertical line at step 122 indicates the transition to a model including RNA secondary structure. **i)** Swarm plot of the maximal Pearson correlation coefficients achieved after SGD optimization for all 79 proteins (x-axis), subdivided by RNA-binding domain type (y-axis). For each RBP the highest observed 6-mer R-value is color-coded. Within each domain category, the names of the RBPs with lowest and highest correlation are indicated. RBPs for which RNA structure is known to contribute to binding affinity in orange. Red vertical bars indicate the median correlation within each category (KH=0.95, ZNF=0.93, RRM=0.95, mixed=0.93, other=0.92, excluding structure-binding ZNF326, RBM22, RC3H1). See also Fig. S2 and Table S2.

### A single set of parameters describes binding across different RBP concentrations

The computational optimization employs an iterative, stochastic gradient descent (SGD) algorithm to find the model parameters most consistent with the experimental data. In addition to the PSAMs, one free parameter for each sample represents the extent of non-specific (background) contributions, modeled as admixture of input sequences. From the PSAMs and background parameters we derive the expected 6-mer R-values for an RBNS pulldown experiment as a function of the predicted RBP occupancies for a random sample of RNA sequences from the input library. The discrepancy between the experimentally observed and the predicted R-values serves as an objective function, which we aim to minimize via SGD (Methods). Minimizing this objective function amounts to adjusting the PSAM values to maximize agreement between model predictions and experimental RBNS data. Specifically, we minimize the total squared difference between predicted and observed log-R values of 6-mers (referred to as ‘model error’). With each iteration, we tune: the relative affinity entries for non-cognate bases in each PSAM, *K*_*d*_^*model*^ for each PSAM, and the background parameter for each sample.

Applying this procedure to four RBNS samples at TRA2A concentrations of 5, 20, 80, and 320 nM, the optimization decreased the total model error across concentrations by more than 2.7-fold (Fig. 2g,h). In parallel, the correlation coefficient between predicted and observed R values increased from 0.9 or below to 0.94 or above (Fig. 2g,h), demonstrating good agreement between the optimized PSAMs and experimental data. The model predicts a shift from binding mostly to a few, high-affinity sequences at 5 nM [TRA2A] to binding many more sub-optimal sequences at 320 nM [TRA2A] (Fig. 2g), matching the experimental data for TRA2A at concentrations spanning almost two orders of magnitude. Supporting the utility of our R-value-based PSAM optimization, we evaluated the PSAMs directly on the pulled down sequence set for TRA2A and found that observed and expected affinity distributions converged (Fig. S2a).

### RBPamp features together capture affinity landscapes of diverse RBPs

To more broadly survey RBPamp performance, we analyzed all 78 RBNS datasets previously published (Dominguez et al., 2018; Lambert et al., 2014) and new data generated for HNRNPA1. Overall, we observed strong agreement between the optimized model predictions and experimental data, with correlations ranging from 0.76 (TAF15) to 0.97 (MSI1) across diverse RBPs containing RRM, KH, Zinc-finger, or other types of RBDs (Fig. 2i, Supplementary Table S2). Among the RBPs with the lowest correlations were RC3H1, RBM22, and ZNF326, which are known to have preference for specific RNA secondary structures (Dominguez et al., 2018; Zhang et al., 2017). Such structural specificity is not representable using PSAMs, except in exceptional cases, e.g., a basepair could theoretically be represented by a set of four PSAMs each enforcing one of the four possible Watson-Crick pairs; we saw hints of this sort of thing in just a few cases (Fig. S2b). Excluding RC3H1, RBM22, and ZNF326, we observed a median correlation across 76 RBPs of *r =* 0.93 (Fig. S2c).

We sought to understand the extent to which each aspect of the RBPamp model contributes to performance. For this purpose, we systematically re-ran RBPamp with each feature disabled in turn, and compared results to the full model after the first round of PSAM optimization. First, we observed that allowing multiple PSAMs to describe the affinity landscape of an RBP reduced model error for virtually all RBPs (*P* < 2.2 × 10^−14^, binomial test *n*=68 out of *N*=74, Fig. 3a). As expected, the difference was most pronounced for RBPs that were assigned larger numbers of PSAMs (3, 4 or 5) (*P* < 0.01, *P* < 0.001, and *P* < 4 × 10^−10^, by 1-sample *t*-test, for correlation and similarly for model error (Fig. 3b, Fig. S3a, S3b).

**Figure 3.**
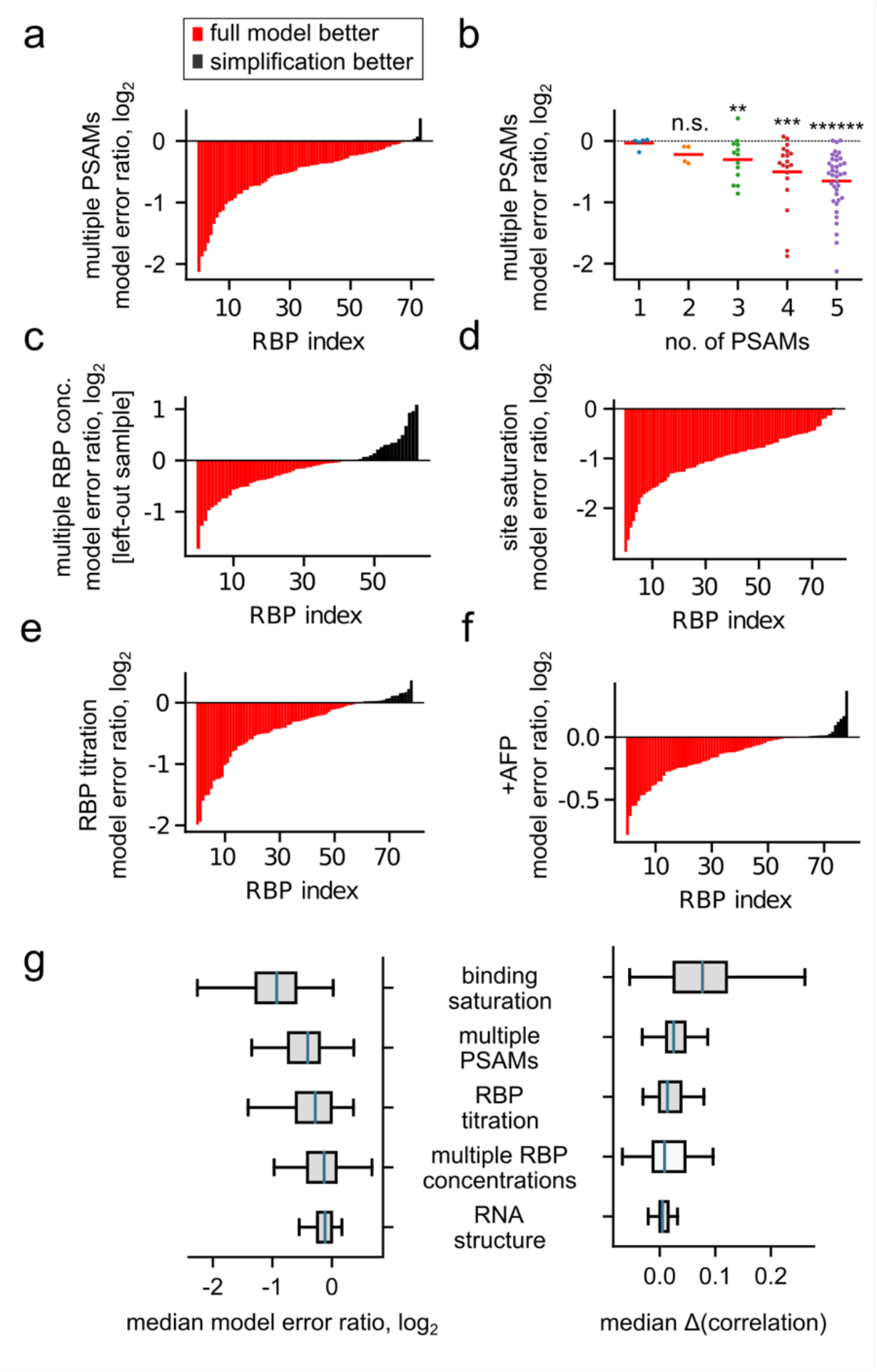
Distinct contributions of all biophysical features of the model. **a)** Bar plot of the log_2_ ratio of model error comparing the full model, allowing for up to 5 PSAMs per RBP, and a restricted model allowing only one PSAM across proteins ranked from lowest to highest difference. Values below 0 (red, *n*=68) indicate superiority of the full (multiple PSAM) model, while those above 0 (black, *n*=6) indicate superiority of the simplified (single PSAM) model. RBPs which were assigned only one PSAM regardless of restriction were excluded (*n*=5). Multiple PSAMs significantly outperform single PSAMs (*P* < 2.2 × 10^−14^, binomial test). **b)** Swarm plots of model error difference (as in a) for RBPs grouped by the number of assigned PSAMs in the full model. Differences are significant for 3, 4, and 5 PSAMs (*P* < 0.01, *P* < 0.001, and *P* < 4 × 10^−10^, by 1-sample *t*-test). **c)** Comparison of PSAMs derived from three RBP concentrations (top 3 based on maximal 6-mer R-value) to PSAMs from the best sample alone (rank 1), with colors as in a. The difference in model error for the held-out sample (rank 4) is shown (*P* < 0.0043, binomial test *n*=45 out of 66). **d)** Comparison (as in a) of a model with saturation (corresponding to black curve in Fig. 2a) to a simplified model with linear occupancy (blue curve in Fig. 2a). (*P* < 2.7 × 10^−22^, binomial test *n*=78 out of 79). **e)** Comparison (as in a) of a model with self-consistent determination of free RBP concentration to a simplified model with [*F*] = [*P*]. (*P* < 4.2 × 10^−6^, binomial test *n*=60 out of 79). **f)** Comparison (as in a) of optimized PSAMs before and after inclusion of an RNA accessibility footprint region in the model (*P* < 1.3 × 10^−6^, binomial test, *n*=61 out of 79). **g)** Box plots of log_2_ ratios of model error (left, reduction means better fit) and difference of correlation (right, increase means better fit), summarizing data from panels a-f above and from Figure S3a-f, respectively. See also Figure S3 and Table S2.

We also explored the value of running RBPamp on multiple samples obtained at different RBP concentrations versus using a single “best” sample. The single sample with highest R value has typically been used previously (Dominguez et al., 2018; Iwasaki et al., 2016; Lambert et al., 2014; McGeary et al., 2019; Miranda et al., 2017). Using samples from 3 concentrations reduced the model error when tested on a held-out fourth dataset, relative to PSAMs derived from only the single concentration with highest 6-mer R value (Fig. 3c, *P* < 0.0043, binomial test, *n* = 45 out of 66).

Next, we explored the impact of mass-action effects. We re-ran RBPamp assuming either that binding site occupancy is a linear function of free RBP concentration (commonly assumed when binding sites are in excess over RBP), or that free RBP equals total RBP (commonly assumed when RBP is in large excess over binding sites). We observed that the full model provides a substantially better fit than either of these simplifications (Fig. 3d,e; saturation *P* < 2.7 × 10^−22^, binomial test *n*=78 out of 79; RBP titration *P* < 4.2 × 10^−6^, binomial test *n*=60 out of 79; comparison of correlations in Fig. S3d, S3e), supporting the importance of considering these features.

Lastly, we compared RBPamp results based on primary sequence alone to those obtained when incorporating effects of secondary structure as described below (Fig. 2c,h). We found that improvements in model error were typically modest (Fig. 3f) but highly significant (*P* < 1.3 × 10^−6^, binomial test, *n* = 61 out of 79), and similarly for model error (Fig. S3f). In summary, each RBPamp feature improved modeling of the data, with the largest contributions from considering binding site saturation and allowing multiple PSAMs (Fig. 3g). All variant model results are listed in Supplementary Table S2.

### Analysis of accessibility can reveal the RNA footprint of an RBP

It is well-established that RNA structure competes with RNA binding for many RBPs (Kazan et al., 2010; Li et al., 2010), and that RBP-bound RNAs tend to have lower probabilities of basepairing within high-affinity *k*-mers (Dominguez et al., 2018; Taliaferro et al., 2016). We modeled the impact of RNA structure by considering that the relevant states of a binding site are: a) unfolded and unbound, b) folded and unbound, or c) bound along a “footprint” of the RBP, with no basepairs inside that region (Fig. 2c), following (Hackermüller et al., 2005). We define the “accessibility footprint” (AFP) as the minimal RNA segment that cannot be base-paired while the RBP is bound; we expect the AFP to correspond closely with the set of bases that are in direct contact with the RBP (Supplementary Notes).

To explore this idea, we considered the interaction of Rbfox family proteins with the 7-mer UGCAUGU, a known high-affinity motif bound by the Rbfox RRM (Auweter et al., 2006). We stratified occurrences of UGCAUGU in pulldown and control datasets by their RNA secondary structure accessibility predicted by a standard thermodynamics-based *in silico* folding algorithm (Bernhart et al., 2011). We observed that the R value of this 7-mer is strongly dependent on accessibility, increasing by two orders of magnitude from the lowest to the highest accessibility bins (Fig. 4a). The dependence of the R value on accessibility, representing the probability that all seven nucleotides are simultaneously unfolded, is much stronger than its dependence on the mean of individual base pairing probabilities across the motif (Dominguez et al., 2018; Lambert et al., 2014), motivating our use of accessibility here.

**Figure 4.**
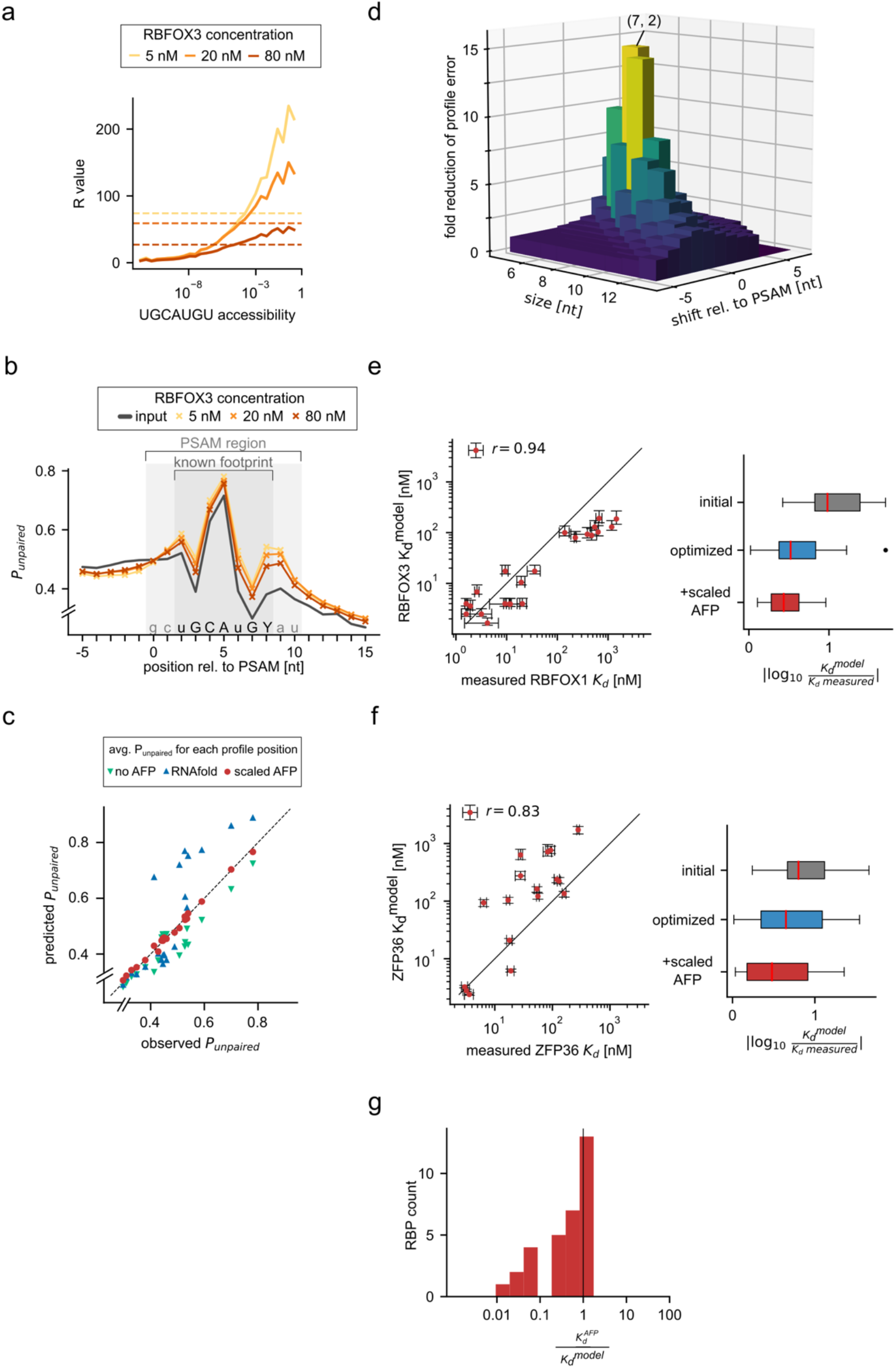
Impact of accessibility on RBP binding and inference of accessibility footprint. **a)** Measured 7-mer R value of known high affinity sequence UGCAUGU (y-axis), plotted as function of *in silico* predicted 7-mer accessibility (x-axis, log-scaled) for different RBFOX3 concentrations (colors), with corresponding dashed horizontal lines indicating overall UGCAUGU R value for each sample. **b)** Profile of mean per-base P_unpaired_ probabilities (y-axis) relative to RBFOX3 PSAM start (x-axis), averaged over all matches in RBNS reads, and weighted by affinity (Methods). Colors as in a, with addition of input control (black). PSAM region (light gray) and known footprint (dark gray) are shown. Preferred sequence of the PSAM for orientation (lower/upper case=low/high discrimination, Y=U or C). Positions with known direct contact to RBFOX3 in dark gray area (“known footprint”). **c)** Predicted mean P_unpaired_ (y-axis) versus mean P_unpaired_ (x-axis, as in b) for 5 nM RBFOX3 profile for different positions relative to the RBFOX3 PSAM. Predicted means were calculated by re-weighting sequences by simulated RBP-binding, under 3 scenarios: ignoring RNA folding (green); requiring accessibility of a 7 nt footprint by RNAplfold (blue); or using a scaled version of RNAplfold accessibility (red). **d)** Fold reduction of profile error (z-axis) is plotted for different footprint sizes (x-axis) and start positions relative to PSAM (y-axis) as a three-dimensional “lego-plot”. The peak occurs at footprint size=7, start=+2, corresponding to UGCAUGY. **e)** Left: scatter plot of RBPamp-estimated K_d_ for RBFOX3 (y-axis) after footprint-aware PSAM optimization (red) for 22 specific 7-mers plotted against SPR-measured K_d_ of RBFOX1 (x-axis) for the same sequences as measured previously by SPR from (Auweter *et al*., 2006; Stoltz, 2015) for the same sequences. Pearson *r*=0.94, P < 5e-11. Right: box plots of absolute deviations (log_10_ ratios) between predicted and observed K_d_ after PSAM initialization (gray), after SGD optimization (blue), and after second round of SGD with scaled footprint (red). Scaled footprint K_d_ predictions match the measured K_d_ significantly better than the initial draft PSAMs (P < 1.6e-6, Mann-Whitney U test). **f)** Same as e) but for ZFP36 using data from (Brewer *et al*., 2004) (MWU < 0.025). **g)** Histogram of ratios of K_d_^model^ values with scaled footprint over K_d_^model^ estimated without considering RNA secondary structure. Two RBPs (HNRNPCL1, SRSF4) had unreasonably low ratios (< .01), suggesting that K_d_^model^ estimates are unreliable, and were excluded. See also Figure S4 and Table S3.

We asked whether the position and size of an RBP’s AFP could be inferred from RBNS data, in the absence of structural information. For this purpose, we interrogated the probability of being unpaired (P_unpaired_) for individual nucleotide positions along binding sites by averaging over millions of sequences, each weighted by motif affinity (Methods). As expected, we observed an RBFOX3-concentration-dependent increase in P_unpaired_ in a region overlapping the RBFOX3 PSAM, with most pronounced effects at the lowest RBFOX3 concentration of 5 nM (Fig. 4b). The P_unpaired_ profile around the Rbfox RBNS motif in the input library was highly non-uniform, as were those of many other motifs tested, reflecting the intrinsic base-pairing propensities of the nucleotide sequences encoded in the PSAM, modulated by the composition of the random RBNS pool and flanking adapter sequences (Supplementary Notes). The elevation of P_unpaired_ in RBFOX3 pulldown samples relative to the input library implies a preference of RBFOX3 for accessibility at these positions.

Differential binding to primary sequence motif variants contributed modestly to differences between the input and pulldown library profiles, but could not explain the overall elevation observed in the pulldown (Fig. 4c, Fig. S4a). Multiplying each binding site’s affinity by the predicted 7-mer accessibility of the known footprint region (Fig. 2c) and re-weighting all sequences by the resulting structure-dependent binding probabilities substantially over-estimated the increase in P_unpaired_ seen across the footprint region (Fig. 4c, Fig. S4b). This observation suggests that less RNA structure forms under RBNS experimental conditions than predicted by standard thermodynamic folding algorithms, similar to recent observations by (Jarmoskaite et al., 2019) (Supplementary Notes). We therefore introduced an accessibility scaling parameter *a* that scales the predicted free energy differences between folded and unfolded binding sites which underlie the accessibilities (Methods). The value *a* = 0.16, obtained by computational optimization, yields model predictions that match the observed P_unpaired_ profiles substantially better than predictions based on default RNA folding parameters (i.e. *a* = 1.0), or those that ignore structure (*a* = 0) (Fig. 4c, Fig. S4c). Analogous to the model error defined above, we defined “profile error” as the mean-squared deviation between predicted and observed P_unpaired_ across positions of the AFP. This approach enables systematic comparison of different choices of footprint size and position relative to the PSAM. For RBFOX3 we found a distinct optimum with over 10-fold reduction in profile error relative to no footprint for an AFP of 7 nt, starting at position +2 relative to the PSAM (Fig. 4d). Thus, the footprint which minimizes profile error coincides precisely with the UGCAUGY motif at the center of the PSAM, in agreement with previous structural studies (Auweter et al., 2006). This example suggests that AFPs inferred by RBPamp (Supplementary Table S3) may provide clues to structural features of protein-RNA interactions.

Since secondary structure was ignored during the first round of PSAM optimization, the second round of optimization tended to lower the assigned K_d_^model^ values, e.g., reducing these values by about 5-fold and 1.5-fold for RBFOX3 and ZFP36, respectively, while leaving the PSAMs virtually unchanged (Fig. S4d,e). Comparing the resulting PSAM-based affinities of RBFOX3 to independent SPR measurements of the affinity of close paralog RBFOX1 to 22 specific 7-mers, spanning three orders of magnitude in affinity (Auweter et al., 2006; Stoltz, 2015) showed good agreement, especially for high-affinity sequences (Fig. 4e). Reasonable agreement was also observed for sub-optimal (secondary) motifs, which are bound under conditions of high Rbfox activity (Begg et al., 2020). RBNS data using a different RNA pool for the close paralog RBFOX2 also yielded similar results (Fig. S4f). The predicted affinity of ZFP36 for UUAUUUAUU accorded with measurements by fluorescence anisotropy (Brewer et al., 2004), with moderate agreement observed for lower-affinity sequences (Fig. 4f). Analogous comparisons to ITC data for CELF1 are shown in Fig. S4g. These examples show that, at least in some cases, the affinity parameters estimated by RBPamp can correspond well to affinities measured using classical biophysical approaches.

For 43 RBPs, our folding analysis identified an AFP for each PSAM that: (i) decreased the profile error, and (ii) had substantial overlap with the PSAM (Methods). Inclusion of AFPs generally led to higher estimates of affinity, with K_d_^model^ decreasing by ∼40% on average, reflecting the incorporation of competition between RNA folding and binding (Fig. 4g). Thus, the AFP formalism brings RNA secondary structure naturally into the RBPamp model, identifying motif positions likely to require single-strandedness for RBP binding.

### Averaged eCLIP binding often has a monotonic relationship to predicted affinity

It was of great interest to assess the relationship between RNA affinities estimated by RBPamp and *in vivo* binding. The largest *in vivo* binding datasets have been generated using the enhanced crosslinking/immunoprecipitation (eCLIP) method (Van Nostrand et al., 2016). eCLIP offers technical advantages over related techniques and has been applied in a large-scale and highly standardized way, facilitating comparisons across samples and proteins. Both RBPamp models and eCLIP data that passed minimal quality standards for both RBNS enrichment and eCLIP reproducibility were available for 20 RBPs in K562 and/or HepG2 cells, with eCLIP data available from both cell lines for 12 RBPs. For example, TRA2A crosslinks strongly to mRNAs for the BMS1 ribosome biogenesis factor in K562 cells and the pattern of eCLIP signal has clear correspondence to predicted relative affinity (i.e. affinity relative to the optimal motif, of affinity 1.0) (Fig. 5a). Similar correspondence between eCLIP enrichment was widely observed for other RBPs and mRNAs (examples are shown in Fig. S5a-f).

**Figure 5.**
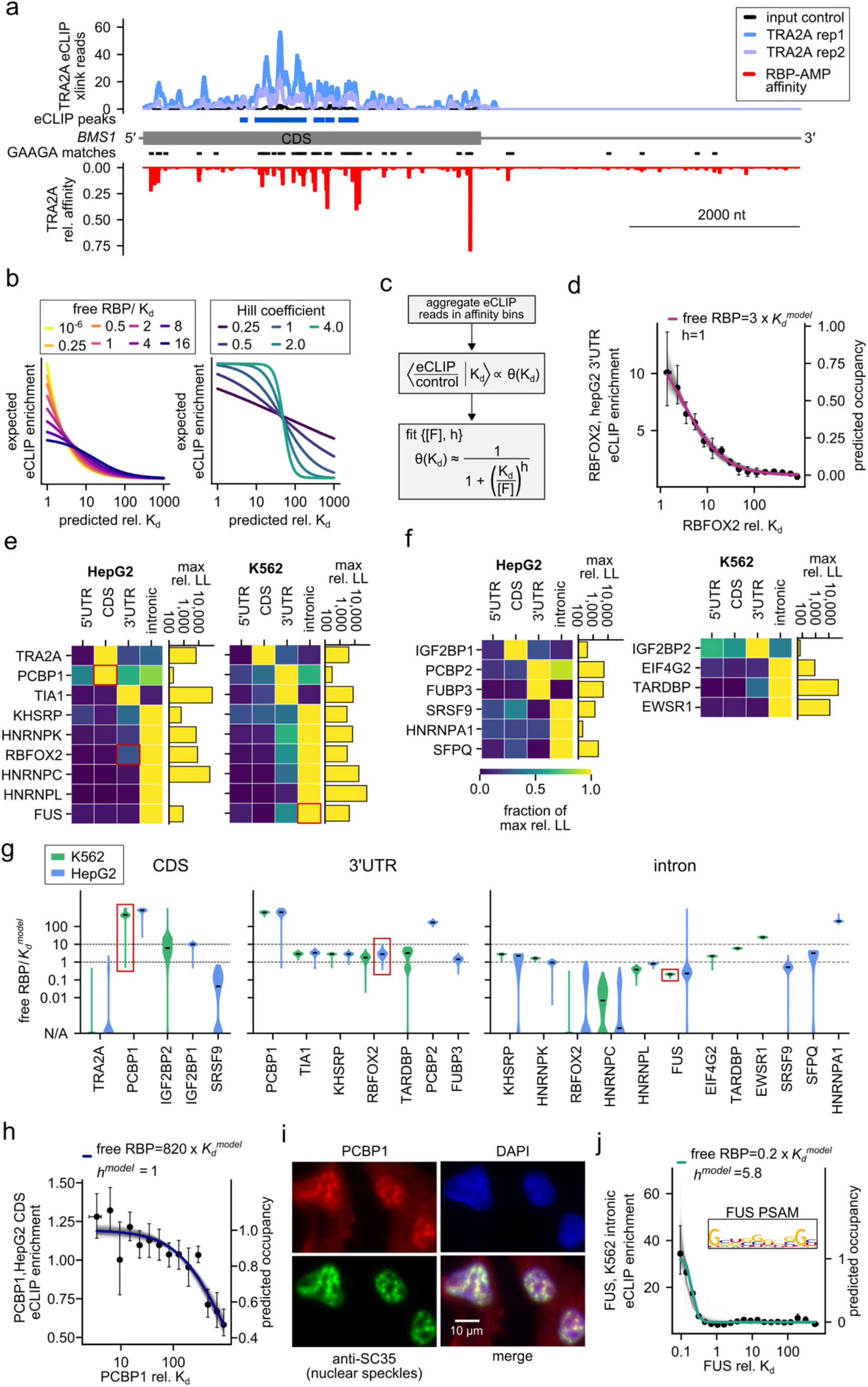
A quantitative model for *in vivo* RBP distribution interrogates eCLIP signal. **a)**Number of eCLIP-read 5’ ends (indicative of TRA2A crosslinking) (y-axis, top) for two eCLIP replicates (blue, purple), and relative affinity predicted by RBPamp (y-axis, below, red) are plotted along the mature mRNA transcript of *BMS1* (gray cartoon in middle indicates location of CDS region and UTRs). Blue boxes indicate eCLIP peaks, while black boxes indicate GAAGA 5mers, a core element of the TRA2A PSAM. **b)** Expected relationship between eCLIP enrichment and binding site affinity at different free RBP concentrations (left) from very low (yellow) to high (blue), and with different degrees of cooperative binding (right) from strong negative cooperativity (*h* = 0.25, purple) to strong positive cooperativity (*h* = 4, green). **c)** Schematic of eCLIP data processing: CLIP data are aggregated across the transcriptome, and binned by predicted local affinity, and the indicated expression for occupancy is used to fit average CLIP enrichment as a function of affinity, with free parameters [*F*] and *h*. For RBPs with multiple PSAMS, K_d_^model^ refers to the highest affinity PSAM. **d)** Result from (c) for RBFOX2 binding to 3’ UTR sequences in HepG2 cells (dots with error bars, left y-axis) relative to site affinities in each bin (x-axis). Error bars: 5^th^ - 95^th^ percentile range of bootstrapped eCLIP sites (left y-axis). Purple line: fit with *h*=1 and [F]/K_d_^model^ = 3, predicting *in vivo* occupancies (right y-axis). Gray lines in the background represent 1000 fits to the bootstrapped data. **e)** Heatmap of row-normalized relative LL of fits performed for K562 (left) and HepG2 (right) eCLIP data, for different transcript regions (x-axis) and RBPs (y-axis); values in each column are expressed as a fraction of the maximum rel. LL observed for the RBP, which is shown at top. The fits shown in panels (d, h, j) are highlighted by red rectangles. **f)** Same as e, but for RBPs for which eCLIP data is only available for either K562 (left) or HepG2 (right) cell lines. **g)** Violin plots of estimated free RBP concentration across bootstrap replicates [*F*]/K_d_^model^ (y-axis, log-scale). RBPs grouped by transcriptomic region for which the best fit was observed (see e,f). Additional, secondary fits (at least 10% of max rel. LL) for RBFOX2, PCBP1, SRSF9. Green: K562; Blue: HepG2 (blue). Red rectangle: fits shown in panels (d, h, j). Dashed lines: corresponding to occupancy 50%-90% for single, optimal site. **h)** Same as (d) but for CDS binding of PCBP1 in HepG2 cells. **i)** Fluorescence microscopy images of HepG2 cells. Red: PCBP1; Blue: DAPI; Green: anti-SC35(SRSF2), marker for nuclear speckles (white bar indicates scale of images). **j)** Same as (d) but for intronic binding of FUS in K562 cells. Insert: FUS PSAM. RBM22 was excluded because of its structural binding preference, TAF15 for extremely low eCLIP reproducibility below 1%, and PUM1 for lack of enrichment of the canonical Pumilio motif in eCLIP. See also Figure S5 and Table S4.

We next asked whether there was a quantitative relationship between predicted affinities and distribution of binding across the transcriptome, using RBPamp to predict binding site affinities along known transcript sequences (GENCODE) expressed in HepG2 or K562 cells. Under the simplest model, in which RBP molecules bind independently, both nucleus and cytoplasm are well-mixed, and RNA structure is ignored, transcriptomic binding sites of similar affinity located in the same compartment (nucleus or cytoplasm) will be similarly occupied by the RBP. Because eCLIP sequencing reads sample RBP-RNA complexes formed in the cell, the ratio of eCLIP signal to input control at a site (the “eCLIP enrichment”) is expected to be approximately proportional to the RBP occupancy of the site. Therefore, this model predicts that in each compartment the eCLIP enrichment will increase monotonically with predicted affinity. Ideally, a sigmoid relationship between eCLIP enrichment and affinity would be expected, with location depending on the ratio between free RBP concentration and K_d_ (Fig. 5b, left). Of course, for any individual binding site the extent of binding in vivo may be altered by competing RBPs or structures, and the measured binding may be distorted by technical factors such as low read coverage of the locus or variability in efficiency of crosslinking or PCR amplification. To reduce the impact of these factors on eCLIP enrichment, we grouped transcriptomic segments of 50 nt into bins of similar predicted affinity, with hundreds or thousands of distinct segments per bin, and used average values for each bin (Fig. 5c). Notably, we find that for many RBPs the mean eCLIP enrichment across bins is indeed a monotonic function of RBPamp-predicted affinity, with approximately the expected shape, as discussed below.

### RBPamp affinities best predict *in vivo* binding in introns and 3’ UTRs

To explore the relationship between *in vivo* binding and predicted affinity, we fit four free parameters to each pair of eCLIP replicates: the concentration of free RBP, *F*^*model*^, an exponent related to the slope, analogous to the Hill coefficient, *h*^*model*^, and one parameter per replicate describing the amount of eCLIP background in the experiment (Methods). We employed a bootstrapping scheme to derive confidence intervals for the per-bin enrichments, the mean estimated K_d_ of sites in each bin (on a relative scale, where the optimal motif is defined to have K_d_ = 1), and the four fitted parameters. Values of the *h*^*model*^ parameter that differed from 1.0 were accepted only if they provided a significantly better fit (by likelihood-ratio test) than a simpler model with three free parameters and *h*^*model*^ set to 1.0.

As an example, we obtained an excellent fit for RBFOX2 binding to 3’UTRs in HepG2 cells with *h*_*model*_ = 1.0, and *F*^*model*^ */ K*_*d*_^*model*^ *=* 3 (Fig. 5d). For RBFOX2, this represents 4.5 to 10 nM of free protein, or roughly 4,500 to 10,000 molecules of free RBFOX2 in the cytoplasm (Milo et al., 2010) (Bionumbers BNID 104519), where most interactions with 3’ UTRs presumably take place. Considering that most RBFOX2 protein is nuclear-localized, this estimate is compatible with estimates of about 300,000 total RBFOX protein molecules in mammalian cells (Schwanhäusser et al., 2011). The above estimate suggests that the *in vivo* occupancy at optimal RBFOX2 binding sites in 3’ UTRs approaches 75% in HepG2 cells, consistent with potentially high activity of such sites.

We performed similar fits across all RBPs which had available RBNS and eCLIP data, separately for 5’ UTR, coding, 3’ UTR and intronic regions. To assess goodness of fit, we compared the resulting log-likelihood (LL) to the LL of a null model with constant, mean eCLIP enrichment independent of predicted binding site affinity (Methods). By this measure of relative LL, we observed a good fit for 19 of 20 RBPs tested, in at least one transcript region/cell type, with high similarity observed between K562 and HepG2 cell data when available (Fig. 5e, f). For most RBP/cell type pairs (18/28) the best fits were observed to the intronic binding data, while the 5’ UTR never had the best fit, likely reflecting the large numbers of potential and actual binding sites in introns (>10^4^ bp of introns per gene on average) – yielding more accurate average enrichment values – compared to the relatively small sizes of 5’ UTRs (∼10^2^ bp per gene). Notably, the CDS region had fewer best fits than the 3’ UTR (6), and for the great majority of RBP/cell type pairs (22/28) a better fit was observed to 3’ UTR than CDS data, despite the smaller average sizes of 3’ UTRs. The most plausible explanation for this difference is that translating ribosomes frequently displace RBPs bound to CDS regions, so that RBP occupancies are disturbed from equilibrium in CDSs, but binding comes closer to equilibrium in 3’ UTRs. Scanning by the 40S ribosomal subunit could have a similar effect in 5’ UTRs. For three of the four cases where CDS was best fit, including TRA2A and PCBP1, predominant RNA binding is likely to occur in the nucleus, avoiding effects of ribosomal displacement. All fitting results are listed in Supplementary Table S4.

### Modeling eCLIP/RBPamp data suggests properties of RBPs in vivo

A trend observed in our modeling of in vivo conditions was that 5 out of 7 RBPs with good fits for 3’ UTR binding had 3’ UTR-derived estimates for *F*^*model*^*/K*_*d*_^*model*^ between 1 and 10, corresponding to ∼50% to 90% occupancy of an optimal binding site (Fig. 5g), while two RBPs had even higher predicted free concentrations (see below). High occupancy of target sites may be required for function in many cases; these observations suggest that cognate binding sites in 3’UTRs are often highly occupied but usually not saturated, enabling moderate changes in RBP activity to meaningfully change occupancy. On the other hand, for intronic binding, estimates of nuclear free protein concentration were typically somewhat lower, roughly evenly split between those where *F*^*model*^*/K*_*d*_^*model*^ was above or below 1. It is important to note that *F*^*model*^ is more difficult to estimate by this approach when *F*^*model*^ << *K*_*d*_^*model*^ (Fig. 5b, left).

On the other hand, we also observed *F*^*model*^*/K*_*d*_^*model*^ >> 1: the estimated free PCBP1 concentration in the nucleus was estimated to be several hundred-fold above *K*_*d*_^*modet*^ (Fig. 5g), and similarly for PCBP2, implying that many suboptimal as well as optimal binding motifs may be saturated with protein (Fig. 5h). Consistent with this inference, PCBP1 has sub-nanomolar K_d_ for optimal motifs and is present at micromolar concentrations in human cells (D. Dominguez, personal communication). Investigation of imaging data from the RBPimage database (Van Nostrand et al., 2020) further revealed that nuclear PCBP1 appears in foci, where it co-localizes with SRSF2 (SC35), a marker for nuclear speckles (Fig. 5i). Based on this marker, an average of 68% of total cellular PCBP1 intensity overlaps with nuclear speckles, which cover only about 11% to 13% of the nuclear area. By comparison, the speckle-overlapping fraction was 17% for FUS and 28% for RBFOX2 (Fig. S5g, h). The high occupancy of both optimal and suboptimal motifs observed for PCBP1 may therefore be explained in part by eCLIP signal deriving predominantly from PCBP1 protein in highly concentrated in nuclear speckles.

The slope-related parameter *h*^*model*^ might relate to whether the RBP binds cooperatively, but might also be influenced by a variety of technical factors, e.g., related to crosslinking efficiency. This parameter had a value of 1 for RBFOX2, FUBP3, TIA1, and PCBP1, which are not known to bind cooperatively, and for all RBPs but one the best fit occurred with *h*^*model*^ ≥ 1. Most of the remaining RBPs achieved significantly improved fits with mildly positive values, mostly 1 < *h*^*model*^ < 2, but higher values were observed for KHSRP, SRSF9, SFPQ, HNRNPA1, and FUS (Fig. S5i). FUS is known to exhibit highly cooperative binding in vitro (Wang et al., 2015). In our analysis, intronic binding of FUS could only be well fit with very strong cooperativity at *h*^*model*^ *> 5* (Fig. 5j). Indeed, our analysis suggests that multiple strong binding sites must be present within a region of RNA (yielding a relative K_d_ < 1) for efficient binding, consistent with the observed dependence of FUS binding on substrate length (Wang et al., 2015). Similarly, hnRNPA1 has been long known to display cooperative behavior (Cobianchi et al., 1988) with relevance for alternative splicing (Blanchette and Chabot, 1999), and HIV biology (Marchand et al., 2002). Likewise, SFPQ has been observed to polymerize (Lee et al., 2015).

Together, our analyses indicate that in vitro-derived PSAMs are highly predictive of in vivo binding along transcripts, particularly in noncoding regions, and can be combined with eCLIP data to yield testable predictions about in vivo occupancies and cooperativity.

## Discussion

Here we have sought to extract precise models of RBP binding from *in vitro* RBNS data, with several potential applications in mind. First, our data support that RBPamp PSAMs can be used to predict binding locations *in vivo* when CLIP data are not available for an RBP, cell type or transcript of interest. Comparisons to available CLIP data may prove useful in assessment of CLIP analysis algorithms, or in detection of binding patterns that are modulated by other factors that interact with the RBP or RNA target. We have also shown that RBPamp can be integrated with eCLIP data to make inferences about in vivo binding site occupancies and cooperativity of binding,

Key features of the RBPamp model such as the allowance for multiple motifs per RBP and the consideration of binding site saturation provided large improvements in modeling of RBP interactions, the latter benefiting from use of data from multiple RBP concentrations (Fig. 2b). Accounting for titration of free RBP by the RNA pool further reduced model error. The combined value of modeling these quantitative features is underscored by the ability of RBPamp to accurately describe RBNS experimental data, across varying RBP concentrations, with relatively few independent parameters: 34 per PSAM, plus one parameter describing the nonspecific background in each sample. Under the biophysical framework used, each PSAM entry should be related to the free energy penalty for substitution of a cognate base by a specific non-cognate base at that position. Thus, the number and interpretability of the model parameters contrast favorably with artificial neural network-based approaches. The inference of the high cooperativity of FUS from the combination of RBPamp and eCLIP data illustrates another potential application of the modeling framework. Together, the examples illustrate an approach to integrating *in vitro* and *in vivo* binding data for the study of RBP biology.

To account for effects of RNA structure, we associated an AFP with each PSAM, whose size and location are estimated from the data, identifying RNA regions within which base-pairing meaningfully competes with RBP binding. This analysis introduced a semi-empirical scaling parameter to bring the folding predictions into agreement with the experimental observations, an approach that is necessarily imprecise. This parameter was likely needed because of large differences in salt concentrations used in RBNS relative to those used in training of standard thermodynamic models (Supplementary discussion). Nonetheless, estimates for AFP size and position relative to primary sequence motifs appeared reasonable for several RBPs, illustrating another application.

In most applications, the free concentration of an RBP in the cell will not be known, and so neither will the ratio of free protein to K_d_, even if an absolute K_d_ has been determined. Thus, the relative ordering and scaling of affinities are of primary interest in typical applications. Our focus here was therefore on estimating accurate relative affinities of different sequences, and how meaningful the absolute values of other model parameters like *K*_*d*_^*model*^ and *F*^*model*^ are was not systematically investigated. In comparing RBPamp predictions to eCLIP data, we found that RBPamp’s relative ordering of RNA sequences by affinity accorded well with eCLIP enrichment, supporting the ability of our approach to predict patterns of in vivo binding.

An important limitation of our current approach is the consideration of each RBP independently of other RBPs, whereas binding may be impacted by interaction with other RBPs (Dassi, 2017; Huelga et al., 2012), as has been characterized extensively for DNA-binding transcription factors (Jolma et al., 2015). For some RBPs (PUM1, RBM22, HNRNPA1), predicted affinity was not well correlated with CLIP signal, or differed between cell types, suggesting that additional factors may modulate the distribution of these RBPs in vivo. For example, joint binding of RNA by multiple RBPs, sometimes altering specificity, has been observed (Kuwasako et al., 2014; Leeper et al., 2010; Weidmann et al., 2016; Lang et al., 2021). Such joint binding could direct the protein to RNAs containing motifs distinct from those bound by the protein by itself in vitro, or could distort the shape of the relationship between affinity and eCLIP density. RBPs can also be blocked from their preferred binding sites by competing RBPs, also potentially distorting the relationship between affinity and occupancy in vivo. Indeed, even unrelated RBPs may favor closely related sequence motifs in vitro (Dominguez et al., 2018), and CLIP data from a large swath of RBPs present in a cell type identified frequent binding to the same or overlapping sites, suggesting the widespread potential for binding competition (Plass et al., 2017; Van Nostrand et al., 2020). We envision that RBPamp can provide a baseline model for future studies of competitive or cooperative interactions between RBPs by predicting what in vivo binding should look like if each protein bound in isolation. Similarly, combining collections of RBPamp models of individual RBP affinities could serve as a baseline model of the collective activity of multiple RBPs present in a particular compartment to help detect and disentangle combinatorial aspects of post-transcriptional gene regulation.

## Acknowledgments

We thank Eric Lécuyer for permission to use imaging data from the RBP Image Database (http://rnabiology.ircm.qc.ca/RBPImage) and gratefully acknowledge Xiaofeng “Andy” Wang for providing the raw images and speckle quantifications. We thank Eric van Nostrand and Gene Yeo for insightful discussions and sharing raw eCLIP-enrichment tables for various transcriptomic regions. We thank Xintao Wei from the Graveley lab for support in processing the ENCODE RBP knock-down RNA-seq data. MJ acknowledges funding through EMBO long-term fellowship ALTF 1130-2015. This work was also funded by the National Human Genome Research Institute ENCODE Project, contract U54HG007005 to C.B.B., by the National Science Foundation, Grant number DMR-1719316 to R.B., and others. Source code and raw data: all data are available through the ENCODE portal. All source code is open-source and freely available at https://bitbucket.org/marjens/rbpamp

## Author contributions

MJ conceived and implemented RBPamp, supervised by CB and RB, with support from MG. MJ and CB wrote the paper.

## Methods

### Software Availability

Source code and raw data: all data are available through the ENCODE portal (www.encodeproject.org). All source code is open source and freely available at https://bitbucket.org/marjens/rbpamp.

### RBNS pre-processing

Raw RBNS data consisting of sequence reads from pulldown and input samples were pre-processed as previously described (Dominguez *et al*., 2018). For subsequent analyses flanking sequences shown below were appended to sequencing reads to capture the sequence of the full-length RNA molecules used in the experiment:

5’-GGGAGUUCUACAGUCCGACGAUC-*N*_*L*_-UGGAAUUCUCGGGUGUCAAGG-3’

With *N*_*L*_ indicating the random sequence insert, which was of length *L=*40 for RBFOX2, MSI1, MBNL1, PTBP3, SRSF2, and RBM47, and *L*=20 for all other RBPs. Sequencing reads containing unassigned bases (N) were discarded.

### Computation of *k*-mer enrichment

RBPamp first computes R values for *k* = 6, 7 and 8, for every sample of each RBP similarly as described in (Lambert *et al*., 2014), but considering up to *k*–1 nucleotides of flanking adapter sequence as well. These values are used at various steps below. We find that 7-mer R values provide a robust way to rank samples at different concentrations for the visibility of sequence enrichment due to RBP-binding. For PSAM construction, we utilize 8-mers which provide more detail but need to be restricted to a set of significantly enriched 8-mers in order to maintain statistical power. Lastly, for efficiency reasons, the RBPamp model optimization employs 6-mer R-values to monitor the convergence between predicted and observed sequence composition. We found that using 7-mers for this purpose added to the run time without improving the results.

### Sample selection and single versus multiple concentration runs

RBPamp first computes 7-mer R values for data from all available experiments (concentrations) for a given RBP. Then, the 7-mer with the maximal R value across all samples is selected as the best estimate of a sequence with direct affinity to the RBP and used as a diagnostic for sample quality. Samples with diagnostic R value below 1.2 (less than 20% enriched) are excluded. Remaining samples are then ranked by the R value of the diagnostic 7-mer. All runs utilized the top 4 available samples, with two exceptions: the single-sample runs utilize only the rank 1 sample; and the multiple concentration runs from Figure 3C utilize ranks 1, 2, and 3, saving the rank 4 sample for testing of the single- and multi-concentration models.

### Position-Specific Affinity Matrices

RBPamp describes primary sequence affinity of an RBP via a set of PSAMs, **M** = {A^1^,…,A^N^}. Each PSAM A^k^ has 4 rows (corresponding to A, C, G, U), and *n*_*k*_ columns (motif positions). The matrix elements represent the relative affinities of each nt at each motif position, relative to the cognate nt at the position, which is assigned an affinity of 1. Matrix elements are constrained to be in the range [10^−6^, 1], where 10^−6^ is a (somewhat arbitrary) minimum value to avoid divisions by zero. RBPamp further assigns an affinity A^k^_0_ to each PSAM, representing the absolute affinity of the cognate (optimal) sequence of nucleotides for that PSAM. Thus, the A_0_ value of a PSAM (expressed in units of nM^-1^) is equal to 1/K_d_^model^, where K_d_^model^ is the dissociation constant of the cognate sequence of the PSAM in the model.

Considering an RNA sequence *S* of length *n*_*k*_, and associated index vector *s*, where *s*_*i*_ is the index of nt *S*_*i*_ in the vector (A,C,G,U) (i.e. *s*_*i*_ = 1 if *S*_*i*_ = A, *s*_*i*_ = 2 if *S*_*i*_ = C, etc.), the PSAM A^k^ assigns an affinity to *S* given by

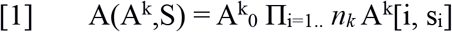

where A^k^[i, j] denotes the matrix element at column *i* and row *j*. For a set of N PSAMs **M**, all with *n*_*k*_ columns, the contributions are simply summed:

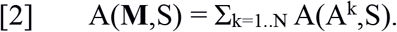

Similarly, when calculating the affinity to a sequence S’ of length L > *n*_*k*_, with associated index vector s’, the affinities at each position are summed:

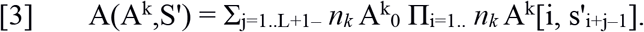

When considering a set of N PSAMS of the same or different lengths, the contributions of individual PSAMs are summed as in eq. [2] above.

### R value background correction

RBPamp generates initial PSAMs by iteratively selecting significantly enriched 8-mers. Specifically, 8-mer R values are averaged across the selected samples (see ‘Sample Selection’). Then, these mean 8-mer R values are corrected for non-specific contributions. For this, we employ a simplification of RBNS, pretending that binding happens directly to k-mers. Knowing that each RBNS oligo contains additional k-mers that were irrelevant for binding and that there is always an amount of RNA that non-specifically winds up in each pull-down sample (background), we split the total k-mer abundances in a pull-down sample into *X* specific and *Y* non-specific k-mer occurences. These numbers are bulk quantities of the sample. We then express the observed R-value *R* for a given k-mer as a function of *X, Y* and the *true* R-value *R’*, that would be observed in the absence of background (*i*.*e*. for *Y=0*):

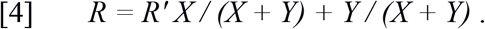

For any single non-specific k-mer *R’* = 0 and its observed enrichment *R*_*ns*_ is expected to take on the value of

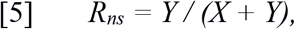

shared among all non-specific *k*-mers. Therefore *R*_*ns*_ allows estimation of the relationship between specific *(X)* and non-specific contributions *(Y)* to the pull-down:

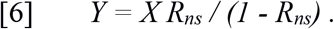

Inserting this expression into [4] and solving for *R’* yields:

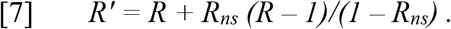

In practice, we take the lower 5^th^ percentile of the R-value distribution as a robust estimate for *R*_*ns*_ and use this to compute *R’* for each RBNS sample.

### Selecting significantly enriched 8-mers

Next, *R’* is converted to a z-score by subtracting the mean and dividing by the standard deviation. We discard 8-mers with *z* ≤ 4 and use the remaining, significantly enriched 8-mers, together with their *R’* value, for the initialization of PSAMs. In the absence of saturation, we expect *R’* to be proportional to direct binding probability and thus a linear function of affinity.

### Initial construction of PSAMs

An initial motif matrix, not yet normalized to PSAM form, is then initialized from the 8-mer with maximal R’ by setting the matrix elements corresponding to the bases in this 8-mer to the R’ value of that 8-mer, initializing other matrix elements to zero. Next, all remaining significantly enriched 8-mer sequences are aligned with this motif matrix, without gaps. The alignment score is computed by summing the matrix elements corresponding to the 8-mer in question and dividing by the maximal possible score (the sum of the column maxima). For example, the 8-mer used to initialize the matrix (which is no longer available) would be assigned a score of 1, while an 8-mer one mismatch away from this 8-mer would be assigned a score of 7R’/8R’ = 0.875, as would a “shifted” 8-mer whose first 7 nt matched the last 7 nt of the top 8-mer (or vice versa). The 8-mer with the highest alignment score is selected (with ties broken by highest R’ value), and merged with the motif matrix by adding the R’ value of the new 8-mer to the existing matrix element at each aligned position. Thus, merging an 8-mer with a motif matrix may result in expansion of the matrix to including additional column(s), and/or addition of variant bases at some positions, while strengthening the consensus at other positions. Because the matrix has been altered, all remaining significantly enriched 8-mers are re-aligned (as described above). This process is repeated until the best alignment score is < 0.75, where for reference an alignment shifted by two or with two mismatches (or one shift and one mismatch) in the first round would have an alignment score of 0.75. We observe that this procedure tends to group 8-mers that have a common core element, because the weight of such a core element tends to keep growing relative to the edges of the motif.

Once all 8-mers compatible with the current motif matrix are exhausted, we select from the remaining significant 8-mers the one with the highest R’ value and construct a new motif matrix. The process of aligning, merging, and addition of a new matrix is repeated until all significantly enriched 8-mers are exhausted or 5 matrices have been built. From this point, any remaining significant 8-mers are simply merged with the matrix to which they align best.

After all significantly enriched 8-mers have been assigned to a motif matrix, we discard matrices that have fewer than ten 8-mers assigned (unless there is only one matrix left) and reassign these 8-mers to the remaining matrices with the best alignment score. This step prevents carrying PSAMs that are built from a handful of sub-optimal sequences through the subsequent optimization; in practice, it never led to dropping a PSAM seeded by a k-mer with high *R’*. Each remaining motif matrix is then pruned: we first drop peripheral (left-most or right-most) columns where the sum of the column elements represents less than 1 percent of the total weight (sum of R’ values) accumulated over all columns. If the matrix is still wider than a maximum of 11 nt, we continue to remove the peripheral columns with the lowest total weight until the width of 11 is reached. We chose this maximal width as a reasonable compromise between desire for detail on the one hand and the need to have sufficient representation of motifs in random sequences of typical size of 20 nt on the other hand. The pruned matrix is then converted to an approximate PSAM by the following procedure: we compute a per-column pseudo-weight

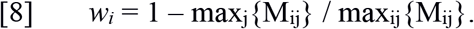

Analogous to pseudocounts used in construction of weight matrices, the *w*_*i*_ value is added to each entry in column *i*. The formula used ensures that this weight is zero for the most important columns (overlapping core elements), and has higher values approaching 1 for columns with lower maxima, serving to smooth peripheral columns with few contributing 8-mers. The resulting matrix is then normalized to values between 0 and 1 by dividing each column vector M_i_ by the column maximum max_j_{M_ij_}. Finally, we add a pseudocount of 0.001 to all elements and renormalize to enable tuning of every PSAM entry by gradient descent. An initial estimate for the relative affinity of each PSAM in a set of PSAMs is obtained by storing the highest 8-mer R’ value that was used in the construction of each PSAM. PSAMs are sorted by this R’ value und their relative affinity is set to the ratio between a PSAM’s highest 8-mer R’ value and the highest 8-mer R’ value observed overall.

This approach can be applied using other *k*-mer sizes, but we have found that the length 8 represents a reasonable tradeoff between the ability to capture details of the interaction while preserving statistical power to compute accurate R values.

### Graphical Representation of PSAMs

Following (Foat, Morozov and Bussemaker, 2006), we compute the *discrimination* for each PSAM column *i*:

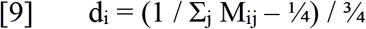

If each element in a column is one (no discrimination, or ‘N’), *d* is equal to zero. On the other hand, if only one element in a column is one and the other three are zero (only one base has affinity for the protein), then *d*=1. To visualize a PSAM, we multiply each column by its discrimination value, and arrange nucleotides on top of each other in order of, and scaled by, their PSAM values. This creates plots that are similar to the commonly used sequence logos, but without making assumptions about background nucleotide frequencies, which are needed to compute relative entropy-style information content/bit-score values. Visualization code uses svgpath2mpl code from Nezar Abdennur (https://github.com/nvictus/svgpath2mpl).

### Local RNA structure accessibility

We utilize RNAplfold 2.2.5 (Bernhart, Mückstein and Hofacker, 2011; Lorenz *et al*., 2011), slightly modified to read and fold one line from stdin at a time. RNAplfold is called with parameters -O -u 12 -W {L} -T 4 (or -T 22 depending on the experiment), which disables the sliding-window functionality (by setting window size to the sequence length), sets the folding temperature to 4 (or 22) Celsius, and reports unfolding energies (“open energies” in RNAfold terms) in kcal/mol for sub-sequences (here interpreted as RBP footprints) of length *f*=1 to *f*=12 nt. RBPamp collects this output from multiple instances of RNAplfold running in parallel and stores the values in binary format for quick random access. The secondary structure accessibility of a footprint region at position *x* within a given sequence, and of footprint size *f*, is easily derived from the pre-computed local unfolding energy ΔU^f^_x_ as

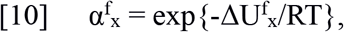

where R is the universal gas constant, and T is absolute temperature (277 Kelvin for typical RBNS conditions). Importantly, as long as binding of only a single RBP to an RNA molecule is considered, this accessibility completely determines the free energy “cost” of excluding base-pairing in the footprint region, eliminating the need for re-folding, which has computational complexity of O(N_reads_ L^3^).

### Partition function model

For each sequence from the RBNS pool the partition function is evaluated as follows:

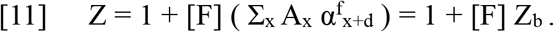

Where [F] is the free protein concentration, and Z_b_ sums over all possible binding locations for a single RBP molecule. We assume that instances of two or more RBPs binding to the same short, random RNA sequence at the same time represent exceedingly rare events in RBNS experiments and therefore exclude such configurations. Under this assumption, the product of [F] and A_x_ represents the Boltzmann-weight for RBP binding at position x ∈ [– n + 1, L – 1] (allowing overlap with flanking adapters). The RNA structure accessibility of a footprint region of size *f* that is shifted relative to the PSAM by *d* positions is factored in. If secondary structure is ignored, α=1. Of note, the computational complexity of evaluating this partition function of the RBP-RNA interaction, including the impact of secondary structure via exclusion of base-pairs from the footprint region, is just O(N_reads_ L).

The probability for binding to a given sequence is then determined by the sigmoid function

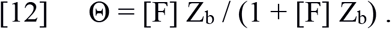

This recovers the well-known Langmuir isotherm binding equation

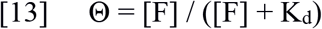

where K_d_ = 1 / Z_b_. In practice, the algorithm considers ∼10 million sequencing reads from the input control, which samples from the random pool used in RBNS, and sub-samples (with replacement) 1 million reads for each round of the stochastic gradient descent (described below). For a given random sub-sample of RBNS reads, RBPamp then predicts the sequence composition of the directly bound RNAs by computing weights for each 6-mer as a function of free protein concentration [F] as follows:

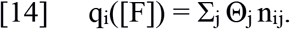

With *j* summing over the reads in the sub-sample and *n*_*ij*_ being the number of occurrences of kmer *i* in sequence *j*. To model an RBNS pulldown sample we further allow for a non-specific background contribution β, consisting of an admixture of sequences from the input pool with 6-mer frequencies *f*_*i*_:

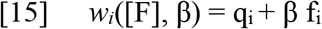

From these we obtain the expected 6-mer frequencies in the bound library, *m*_*i*_, through normalization, and the expected R values by comparing to the 6-mer frequencies in the input pool:

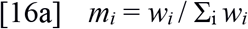

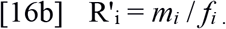

By adding the non-specific background at the level of 6-mer frequencies, we can find optimal values of β without re-computing the partition function.

### RBP titration effects

RBPamp explicitly accounts for RBP that is already bound by estimating that fraction from N_reads_ sequencing reads sampled from the random RNA pool:

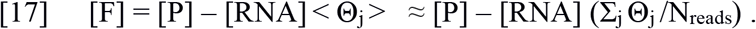

Where [P] is the total RBP concentration, and [RNA] is the total concentration of all RNA sequences present. In practice, we implement this by first binning reads by total affinity (Z_b_) and then computing the binding probability on a per-bin level. This allows quickly finding optimal values of [F] which minimize the discrepancy between [F] and [P] – [RNA](1/N_reads_ Σ_j_ Θ_j_) through binary search, because the problem is convex and one-dimensional.

### Stochastic Gradient Descent

The difference between observed, concentration-dependent log R-values and RBPamp predictions serves as an objective function to be minimized by modifying matrix elements in the set of PSAMs {M}:

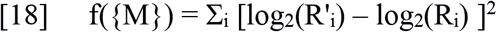

For this purpose, we analytically derive the gradient with respect to all matrix elements

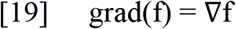

and compute its value for a given set of PSAMs at optimization step *t*. We then use the ADAM (ADAptive Moment estimation) algorithm (Kingma and Ba, 2017) to compute an update vector *u*_*t+1*_ to be added to the matrix elements of the PSAMs at each optimization step from two running variables *g*_*t*_, a decaying average over past gradient values to dampen oscillations, and *s*_*t*_, a decaying average over past, squared values of the gradient used to further amplify components of the gradient that showed lesser variation over past iterations:

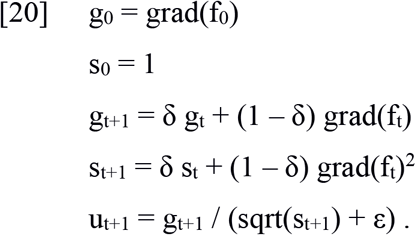

We use parameters δ=0.5 and ε=10^−5^. However, rather than enforcing a fixed learning rate we perform a line-search in the direction of u_t+1_ (Box *et al*., 1969).

### Simplifying approximation testing

We compared the performance of RBP-PSAM with a number of simplified variants. Since we were only interested in the relative performance of these variant models compared to the full model, we chose to ignore RNA structure and stop RBPamp after the first, sequence-only, gradient descent optimization. For the single PSAM model, we simply set the number of allowed PSAMs to be created in the initial phase of RBPamp to one. For the single concentration variant, we ranked all samples (see ‘Sample Selection’ above) and ran RBPamp only on the top-ranked sample (--ranks=1). The multiple-sample comparison was run on ranks 1, 2, and 3 (-- ranks=1,2,3), holding out the fourth-best sample for benchmarking. This was done by using the resulting, optimized PSAMs from single and multi-concentration runs to predict the expected 6-mer R-values for the second-best sample, keeping A_0_ fixed:

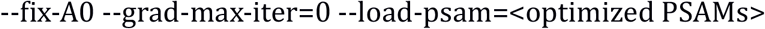

We tested the impact of RBP titration by replacing the self-consistent determination of [F] with [F] = [P] (--excess-rbp) and the impact of linear approximation of occupancy (--linear-occ), while keeping the initial construction of PSAMs and the gradient descent basically identical:

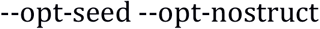

RNA secondary structure accessibility footprints were first identified using

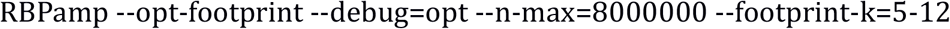

and then a second round of gradient descent with --opt-struct. Overview and comparison plots were generated from the combined output of all RBPamp runs with scripts in the RBPamp/rRBNS folder.

### RBP footprint estimation from secondary structure accessibility

We folded input and pulldown sequencing reads with RNAplfold from the Vienna RNA package as described above, storing accessibilities for sequence windows of varying width *k* and for positions along each RBNS molecule sequenced from the input and pull-down samples. Accessibilities for *k*=1 are identical to P_unpaired_, the predicted probability that a given single nucleotide is not base-paired. We then investigate a random subset of 8 million sequences from the input pool and first compute the Boltzmann weights *Z*_*xj*_ for binding at each position *x* in a given sequence *j* (see eq. [11], but without summation over *x* and setting α=1). Next, we use these per-position weights to compute weighted averages of *P*_*unpaired*_ from a range of positions *x – flank* to *x + l(PSAM) + flank*, covering the PSAM and flanking sequences. The average runs over all possible starting positions in all sequences but the Boltzmann weights ensure that high affinity motif instances contribute the most.

We repeated this procedure for pull-down libraries. The resulting profiles of average *P*_*unpaired*_ values across and around motif instances allow to monitor the selection of the RBP for accessible binding sites without knowing the exact size and position of the RBP footprint. We define the profile error as the sum of squared differences between two *P*_*unpaired*_ profiles.

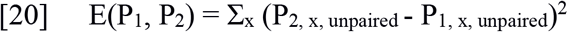

Next, we scan through a range of different footprint lengths *f* from 5 … 20nt in combination with a range of different footprint positions relative to the start of the PSAM (shifts) *s*. For each of these trial combinations, we now evaluate the full model on the random sequences from the input sample, this time with accessibilities α selected from the pre-computed RNAplfold up until we have per-oligo binding probabilities. We then additionally weigh each *P*_*unpaired*_ value that entered into the profile computation by the per-oligo binding probability. These re-weighted profiles *P’* are the predicted profiles we expect to observe at a given RBP concentration, should the RBP binding be modulated by local accessibility as defined through the tested choice of footprint size and relative position. Because the RNAplfold predictions seem to overestimate the impact of secondary structure on binding (see discussion in main text), we introduced a scaling parameter *0 ≤ a ≤ 1*

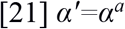

The exponentiation by *a* amounts to a multiplication of the predicted unfolding energy (see [10]). For *a=0*, secondary structure has no impact at all and for *a=1* RNAplfold predictions are not altered. Now, the identification of optimal parameters for a footprint region for a given PSAM consists in minimizing the profile error between observed and predicted P_unpaired_ profiles, summed over all samples *j*.

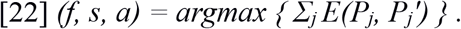

If the resulting footprint overlaps by at least 50% with the PSAM and serves to reduce the profile error over a naive estimate with a=0 by at least 10% (configurable via command line), RBPamp attaches these parameters to the PSAM. This is true for the majority of PSAMs.

### Qualitative comparison to eCLIP data

Raw eCLIP alignments (BAM format) and called peaks (BED format) were downloaded from ENCODE, using the online ENCODE sample selection tool. We wrote custom python scripts using the pysam package (https://github.com/pysam-developers/pysam) to extract and plot the 5’ end positions of eCLIP reads in various transcriptomic regions (part of RBPamp source code).

### Binning of eCLIP data, bootstrapping, and fit

RBPamp affinity models were evaluated on mature mRNA sequences of the transcriptome (GENCODE v28). Predicted affinity profiles were segmented into relevant regions by selecting the highest affinity peak plus 50nt up and down-stream, then setting affinity in this region to zero and repeating the procedure until the highest remaining affinity peak dropped below a cutoff value of 10^−3^ *K*_*d*_^*model*^ (1,000 times worse than a single, optimal binding site). Subsequently, eCLIP and input control reads were fetched from the BAM files (obtained from the ENCODE data portal) for each relevant region and assigned to bins according to the predicted total binding affinity of each region. Per-bin eCLIP enrichment for each bootstrap was computed by dividing the sum of CLIP reads (rep1, 2 separately) by the sum of input reads assigned to each bin.

Bootstrapping was performed by randomly sampling transcript regions, with replacement, from all transcript regions assigned to a bin. Fit between occupancy and eCLIP enrichment was performed using scipy.optimize.minimize on the squared difference of per-bin enrichments (normalized to sum up to one) and the predicted occupancy (normalized to sum up to one), using the L-BFGS-B method.

Fits were performed either treating *h*^*model*^ as a free parameter or fixing it to *h*^*model*^=1.0. A likelihood ratio test was performed to choose between these two fits, requiring P < 10^−5^ to accept *h*^*model*^ values different from one. In addition, the log-likelihood was computed for a negative control “fit” represented by the mean enrichment across bins, with no free parameters. Likelihood was assigned according to a normal distribution for each bin with mean and variance taken from the bootstrap samples.

