## Supplementary Notes, Figures and Tables for "RBPamp: Quantitative Modeling of Protein-RNA Interactions *in vitro* Predicts *in vivo* Binding"

### Index of Supplementary Information for Jens et al.

#### Summary

##### Supplementary Notes

##### Supplementary References

##### Supplementary Figures

- Figure S1 → relates to Figure 1 – details of PSAM initialization
- Figure S2 → relates to Figure 2 – SGD optimization, RC3H1 PSAMs
- Figure S3 → relates to Figure 3 – variant model performance
- Figure S4 → relates to Figure 4 – AFP calibration
- Figure S5 → relates to Figure 5 – eCLIP analysis

##### Supplementary Tables

- Table S1 → relates to Figure 1 – RBP domain and PSAM stats
- Table S2 → relates to Figure 2, 3 – error reductions and correlations for variant models
- Table S3 → relates to Figure 4 – accessibility footprints
- Table S4 → relates to Figure 5 – eCLIP fit results

##### Supplementary Download Items

- rbp_motifs.tgz – RBPamp PSAMs and motif visualizations

### Supplementary Notes

#### Secondary structure prediction and accessibility footprint

We fold RBNS sequencing reads, including the flanking adapter sequences, with RNAplfold (Bernhart, Mückstein and Hofacker, 2011; Lorenz *et al.*, 2011), disabling the windowing logic by setting window size to sequence length. Thus, the output reflects the predicted differences in free energy of two RNA structure ensembles: unconstrained, and with base-pairs restricted to positions outside of the particular stretch of the RNA sequence that is the footprint. This output is furthermore returned for any possible positioning of the footprint, and for sizes from 1 ($P_{unpaired}$) to the maximal value set by us (20).

However, the free energies of the underlying folding ensembles are predicted from nearest-neighborhood (NN) parameters measured at conditions differing from RBNS, and lacking detailed measurements on most loops with more than four nucleotides (Mathews *et al.*, 1999, 2004). Whereas in RBNS approximately physiological salt concentrations are used (150 mM $K^+$, 3mM $Mg^{2+}$) the Turner energy model parameters used by RNAfold have been obtained at very high salt concentrations (1M $Na^+$). The impact of this is difficult to predict and will most probably also depend on temperature. In a recent pre-print (Becker *et al.*, 2019) it was demonstrated that in silico folding predictions become much more accurate for RNA-MaP, an in vitro assay whose equilibrium of RNA and protein is not unlike RBNS, when missing loop energies are added to the energy parameters. In this work, the correction amounted to an offset in the predicted free energies, deriving from the corrected, more de-stabilizing, contribution of two specific loops. However, the authors were only able to apply this correction because they used a custom pool of RNA sequences, designed to fold predominantly into a desired shape and displaying only few loop sequences. In contrast, normal RBNS employs completely randomized sequences. We are therefore unable to determine the correct shift in folding energy for each sequence. Instead, we have to rely on a somewhat crude averaging over all sequences to which the RBP binds. Certain loops will occur more often than others in the subset of sequences that have affinity for a specific RBP. We strive to capture the average effect of the mismatch between RBNS and the Turner energy model by multiplicatively scaling the predicted difference in free energy, and treating the scale as a free parameter. For most RBPs, this approach assigns values to $a$ that are well below 0.5. We conclude that while in silico folding likely over-predicts the stability of secondary structure under RBNS conditions (see Figure S4B), the RBPamp model can still benefit substantially from inclusion of scaled down predictions ($0 < a < 1$), which can sometimes match the experimental data strikingly well (see Figure S4c).

Lastly, we would like to point out the even in the absence of any RBP-binding, interrogating primary sequence motif occurrences in the input library already selects a highly non-uniform profile of expected, per-nucleotide base-pairing probabilities ($P_{unpaired}$, see Figure S4a). We observed that the general shape of this profile does not change much when experimental RBP selection is added, rather the protein selection of more accessible instances of the motif raises the overall accessibility across the relevant bases. This elevation of $P_{unpaired}$ is usually the most pronounced for the lowest RBP concentration used in the assay. We caution that motif-finding algorithms which employ straight-forward extensions for the inclusion of secondary structure motifs might fall into a trap, where the strongest signal is actually induced by the primary sequence motif itself. This signal furthermore might differ between experimental conditions, because it depends on the design of the libraries. For instance, RBNS relies on the addition of stereotypical primer handles flanking the random region of each RNA molecule in the pool (Lambert *et al.*, 2014). In consequence, sub-sequences of the random stretch with complementarity to the adapters will display lower accessibility than non-complementary sequences, an effect observed in previous work (Dominguez *et al.*, 2018). We therefore caution that interpretation of the specific ups and downs of the $P_{unpaired}$ profiles as preference for specific configurations of secondary structure might be misguided. We feel that the success of using a conceptually much simpler AFP to reproduce the $P_{unpaired}$ profiles observed in the RBP pulldown data lends support to this notion and might indicate that for many RBPs the main RNA structural preference boils down to the requirement for single-strandedness in a subset of bases.

#### Relationship of AFP to primary sequence motifs

We have observed that the optimal accessibility footprint (AFP) often extends further than the corresponding PSAM or the positions inside the PSAM that display high discrimination for cognate versus non-cognate nucleotides. It is important to point out that the AFP does not necessarily model the stretch of RNA that is directly read out by the RBDs, but most likely approximates the stretch of RNA that would sterically hinder binding if it were forming any RNA base-pairs. If, for example, the RBP forms a binding pocket or channel, it is easy to imagine how a larger stretch of bound RNA would become restricted to more extended spatial conformations, even though the sequence specific RNA-protein contacts may only occur with a more centrally located subset of the nucleobases forming direct and contacts with the protein.

On the other hand, sometimes bases with high discrimination are positioned unexpectedly close to the AFP boundary or even slightly outside of the AFP proper. In these instances, the RBP might have preference for specific secondary structures forming around or close to a preferentially unpaired primary sequence motif. Such structural specificity is not well modeled by the AFP approach and would most likely manifest as exclusion of proximal motif positions from the AFP – which by definition is required to be unpaired for binding. However, we have also observed that the in silico folding results are only compatible with RNA binding in RBNS when they are scaled down (see above). The underlying inaccuracies of the thermodynamic NN-model almost certainly do not affect all primary sequences equally and may therefore translate into inaccurate AFPs which erroneously exclude positions which, for instance, would disproportionately suffer from over-prediction of their binding propensity.

### Supplementary Figures

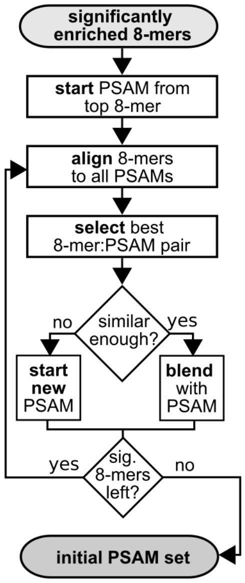

**Figure S1. Block diagram for the PSAM initialization algorithm**

Significantly enriched 8-mers have R value z-score ≥ 4. Starting a new proto-PSAM from an 8-mer equals addition of a new 8-column matrix with elements corresponding to the 8-mer nucleotides set to its corrected R value (see Methods) and zero for unused nucleotides. “Top 8-mer” refers to 8-mer with highest corrected R value. Alignment is performed for each proto-PSAM and 8-mer pair by sliding the 8-mer over the PSAM without gaps. The alignment score is divided by the maximal possible score a single sequence can be assigned. Blending an 8-mer with a proto-PSAM means adding its R-value to the matrix elements corresponding to the 8-mer nucleotides, when positioned over the PSAM columns for maximal alignment score. Sufficient similarity equals alignment score ≥ 0.75. Proto-PSAMs (containing sums of R values) are converted to initial PSAMs by column-wise division by column-maxima, after addition of a pseudo-count (Methods). The resulting PSAM values are in the interval [1e-6, 1].

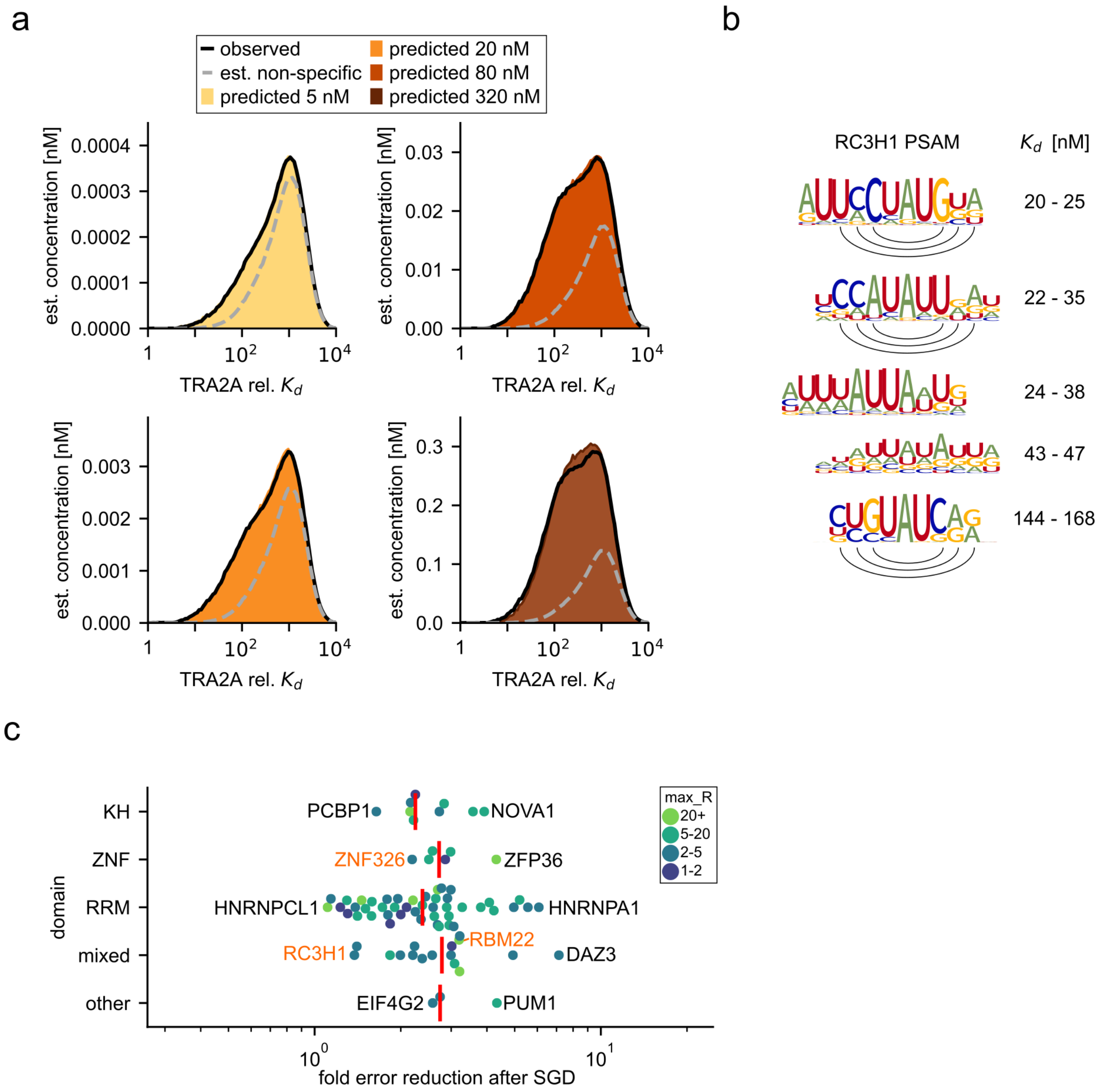

**Figure S2. Relates to main Figure 2. Convergence of affinity distributions and RC3H1 PSAMs**

**s)** Histograms of PSAM-derived affinities of $10^6$ pulldown read sequences for each experimental TRA2A concentration (black lines). Each is fitted as a mix of non-specific (composition of input RNA pool, dashed gray lines), and specific, RBP-bound RNA. Colored, filled histograms: resulting mix. Y-axis scaled by predicted, total TRA2A-bound RNA. **b)** RC3H1 (Roquin-1) binds UAU within hairpin loops (Dominguez *et al.*, 2018), but RBPamp does not model structural motifs. Black arcs: closing C:G, A:U, and G:C base-pairs, with potentially flanking base-pairs in PSAMs. **c)** Swarm plot of reduction (higher is better) in model error (axis on log scale) after SGD optimization for all 79 proteins (x-axis), subdivided by RBD (y-axis). Black/orange labels, red bars as in Figure 2i.

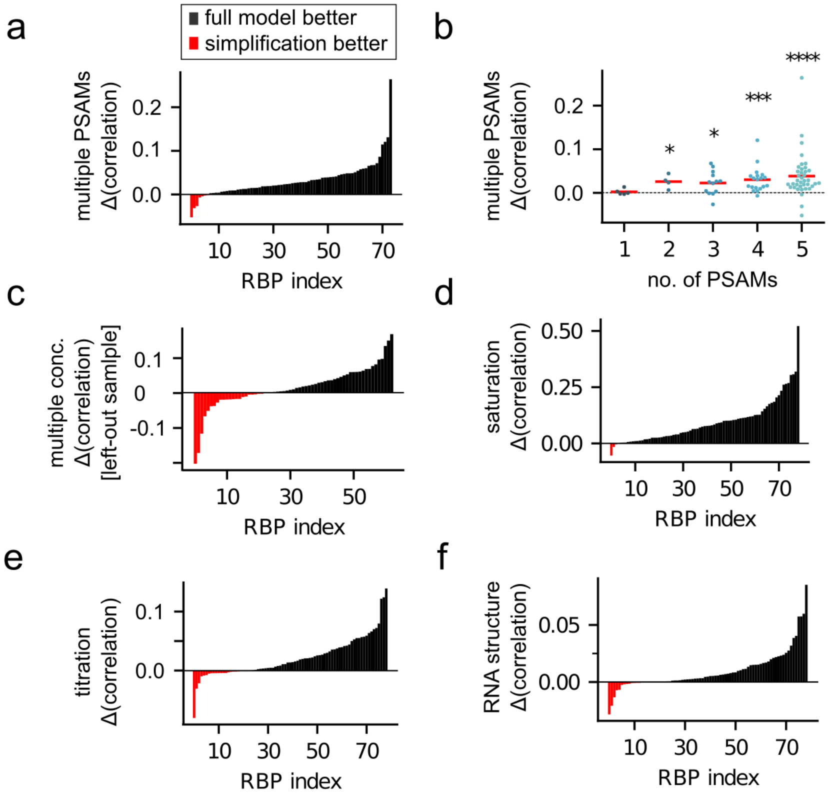

**Figure S3. Distinct contributions of all biophysical features of the model.**

**a)** Bar plot of the difference in final Pearson correlation coefficients of 6-mer R values, between the full model, allowing for up to 5 PSAMs per RBP, and a restricted model allowing only one PSAM across proteins ranked from lowest to highest difference. Values above 0 (black, $n$=67 out of $N$=74 RBPs) indicated superiority of the full (multiple PSAM) model, while those below 0 (red, $n$=6) indicate superiority of the simplified (single PSAM) model. Five RBPs which were assigned only one PSAM regardless of restriction were excluded. Multiple PSAMs significantly outperform single PSAMs ($P < 2.2 \times 10^{-13}$, binomial test). **b)** Swarm plots of correlation difference (as in a) for RBPs grouped by the number of assigned PSAMs in the full model. Differences are significant for 2, 3, 4 and 5 PSAMs (1 sample $t$-test $P < 0.048$, $P < 0.0012$, $P < 2.8 \times 10^{-4}$ and $P < 4.1 \times 10^{-5}$, respectively. **c)** Comparison of PSAMs derived from three RBP concentrations (top 3 based on maximal 6-mer R-value) to PSAMs from the best sample alone (rank 1), with colors as in a. The difference in correlation coefficient for the held-out sample (rank 4) is not significant ($P < 0.065$). **d)** Comparison (as in a) of a model with saturation (corresponding to black curve in Fig. 2a) to a simplified model with linear occupancy (blue curve in Fig. 2a) ($P < 5.3 \times 10^{-18}$, binomial test $n$=75, $N$=79). **e)** Comparison (as in a) of a model with self-consistent determination of $F^{model}$ to a simplified model with $F^{model}=[P]$ ($P < 1.1 \times 10^{-4}$, binomial test $n$=57, $N$=79). **f)** Comparison, as in a, of optimized PSAMs before and after inclusion of an AFP in the model ($P < 9.5 \times 10^{-8}$, binomial test $n$=63, $N$=79).

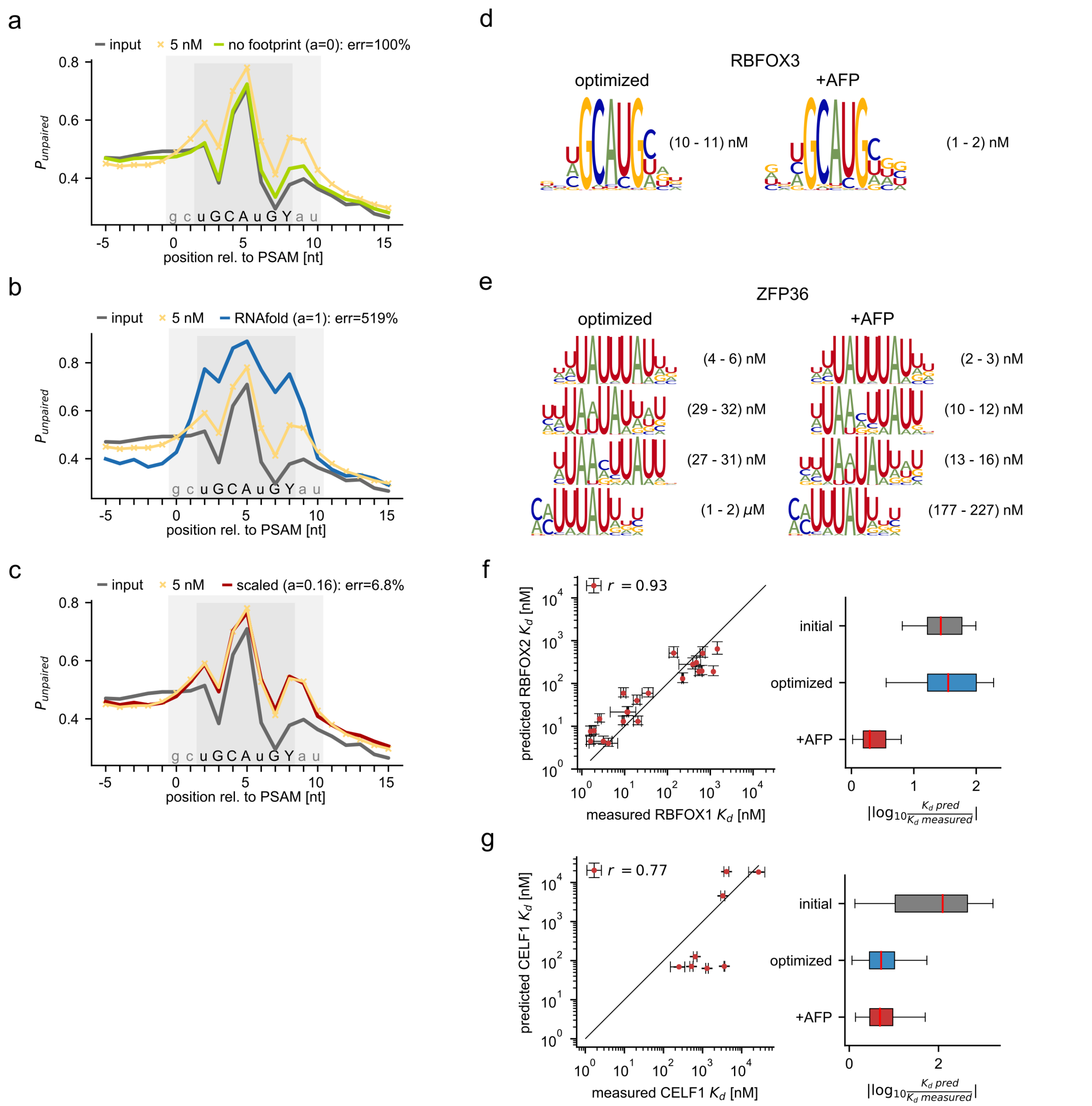

**Figure S4. AFP impact on $P_{unpaired}$, PSAMs and $K_d$ estimates. a-c)** Yellow: average, affinity-weighted $P_{unpaired}$ across PSAM matches (light gray) in RBNS pull-down data from 5 nM total RBFOX3. Black: same for random RBNS input. Dark gray: footprint region (compare main Figure 4B). Expected 5 nM $P_{unpaired}$ profile predicted from input (black) by weighting sampled random sequences with probability of binding and ignoring secondary structure ($a$=0, green, **a**), using unscaled accessibility ($a$=1, blue, **b**), scaling accessibility by $\alpha^a$ (a=0.16, red, (c)). Each y value from (a-c) is plotted against the corresponding yellow line in main Figure 4c. **d,e)** PSAMs after first (left) and second, AFP aware, SGD (right) for RBFOX3 **(d)** and ZFP36 **(e)**. Consideration of structure mainly mostly lowers the estimated $K_d^{model}$, as expected. **f)** Same as main Figure 4e) but for RBFOX2. Note that RBFOX2 RBNS used random 40mers while and RBFOX3 RBNS used random 20mers, yet $K_d^{model}$ values resemble SPR data from (Auweter *et al.*, 2006; Stoltz, 2015) in both cases. **g)** Similar to Figure 4f but for CELF1 SPR data from (Mori *et al.*, 2008) and one ITC data point from (Teplova *et al.*, 2010).

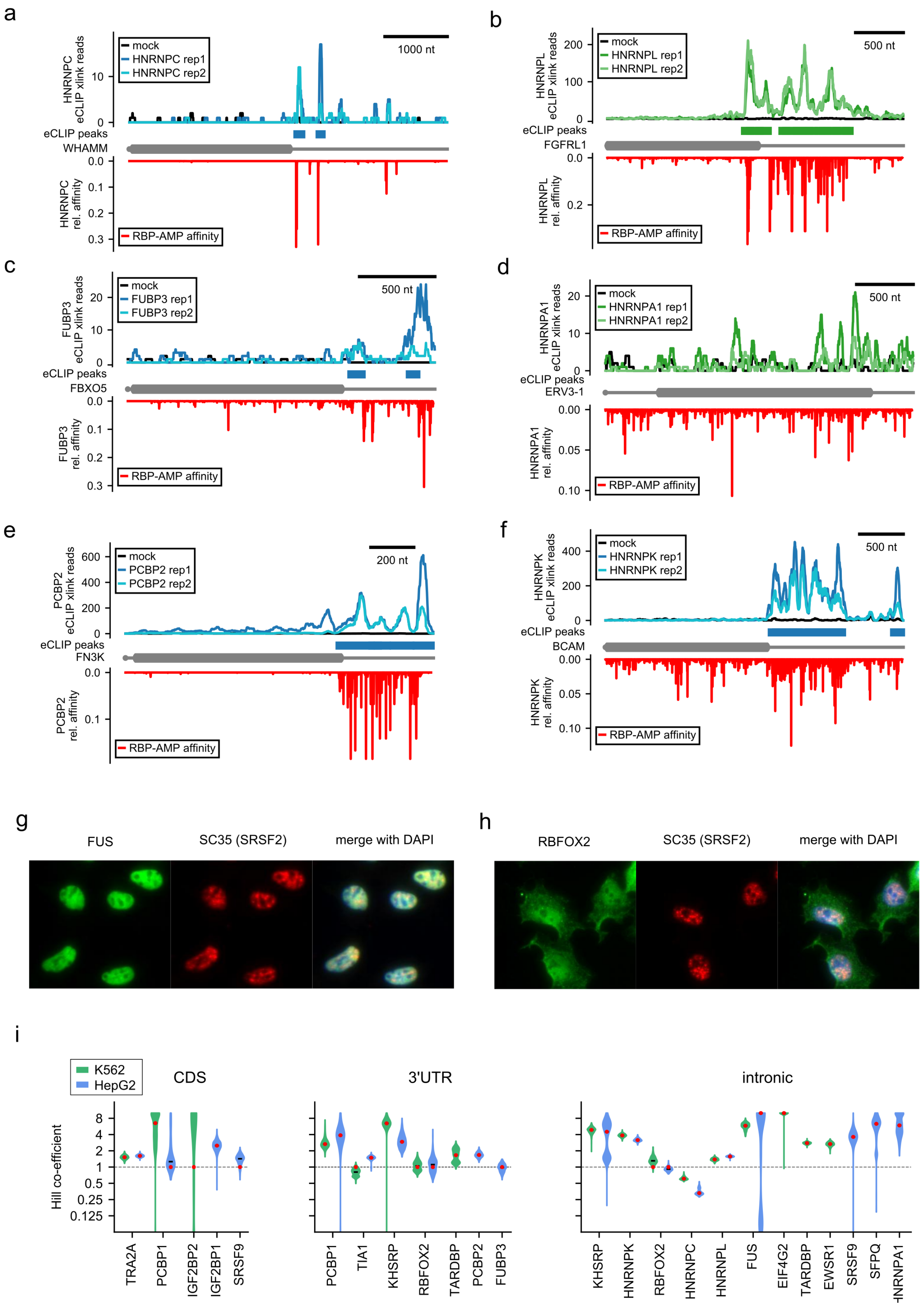

**Figure S5. eCLIP analysis, imaging and cooperativity.**

**a-f)** top, (light) blue/green: profiles of eCLIP crosslink signal (5' end position) for two biological replicates of eCLIP in HepG2 (blue) or K562 (green) cells. Middle, gray: cartoon of the mature mRNA (horizontal line) from 5' to 3' and coding sequence as thick, rounded box for WHAMM (A), FGFRL1 (B), FBXO5 (C), ERV3-1 (D), FN3K (E), BCAM (F). Bottom, red: PSAM-predicted binding affinity for the eCLIP'ed RBP HNRNPC (A), HNRNPL (B), FUBP3 (C), HNRNPA1 (D), PCBP2 (E), and HNRNPK (F). Examples were selected for strong eCLIP enrichment, regardless of agreement with RBPamp. Transcripts with obvious isoform mismatches (short/long 3' UTRs) or very noisy eCLIP signal were not considered. **g,h)** Fluorescence microscopy images, analogous to Figure 5i, showing the lack of strong co-localization with nuclear speckle marker SC35 (red) for FUS (**g**) and RBFOX2 (**h**) in green. Merge with DAPI (right) demonstrates nuclear localization of FUS and nuclear enrichment of RBFOX. **(i)** Violin plots of fitted $h^{model}$ values analogous to those for $F^{model}$ in Figure 5g. Fits were performed 1,000 times over bootstrapped re-samples of transcriptome regions with eCLIP signal from HepG2 (blue) or K562 (green) cells. Black bars: median. Red dots: selected $h^{model}$ values after comparing a model with $h^{model}$ as a free parameter to a model with $h^{model}$ with fixed value of 1. The free parameter model was selected only likelihood ratio test P-value < $10^{-5}$. Distributions are shown if eCLIP data in the indicated transcript region (CDS, 3′UTR, intronic) had either the maximal relative likelihood or no less than 10% of that maximal value (rel. LL, compare to Fig. 5e).

| **Supplementary Table S1.** RBP domain and PSAM numbers | | |
|---|---|---|
| RBP | name of RNA-binding protein | |
| Domains | Annotated RNA-binding domains | |
| n_psams | Number of PSAMs assigned by RBPamp | |
| **rbp** | **domains** | **n_psams** |
| BOLL | RRM,other | 5 |
| CELF1 | RRM,RRM,RRM | 5 |
| CNOT4 | RRM,ZNF,ZNF | 3 |
| CPEB1 | RRM,RRM | 3 |
| DAZ3 | RRM,other | 5 |
| DAZAP1 | RRM,RRM | 4 |
| EIF4G2 | other,other | 4 |
| ELAVL4 | RRM,RRM,RRM | 5 |
| ESRP1 | RRM,RRM,RRM | 4 |
| EWSR1 | other,RRM,other | 4 |
| FUBP1 | KH,KH,KH,Kh | 5 |
| FUBP3 | KH,KH,KH,KH | 5 |
| FUS | RRM,other | 1 |
| A1CF | RRM,RRM,RRM | 5 |
| HNRNPA0 | RRM,RRM | 5 |
| HNRNPA1 | RRM,RRM | 5 |
| HNRNPA2B1 | RRM,RRM | 3 |
| HNRNPC | RRM | 2 |
| HNRNPCL1 | RRM | 4 |
| HNRNPD | RRM,RRM | 5 |
| HNRNPDL | RRM,RRM | 5 |
| HNRNPF | RRM,RRM,RRM | 4 |
| HNRNPH2 | RRM,RRM,RRM | 5 |
| HNRNPK | KH,KH,KH | 5 |
| HNRNPL | RRM,RRM,RRM,RRM | 5 |

| | | |
|---|---|---|
| IGF2BP1 | RRM,RRM,KH,KH,KH,KH | 4 |
| IGF2BP2 | RRM,RRM,KH,KH,KH,KH | 5 |
| ILF2 | other | 5 |
| KHDRBS2 | KH | 5 |
| KHDRBS3 | KH | 5 |
| KHSRP | KH,KH,KH,KH | 5 |
| MBNL1 | ZNF,ZNF,ZNF,ZNF | 5 |
| MSI1 | RRM,RRM | 5 |
| NOVA1 | KH,KH,KH | 4 |
| NUPL2 | ZNF | 4 |
| PABPN1L | RRM | 4 |
| PCBP1 | KH,KH,KH | 5 |
| PCBP2 | KH,KH,KH | 4 |
| PCBP4 | KH,KH,KH | 5 |
| PRR3 | ZNF | 4 |
| PTBP3 | RRM,RRM,RRM,RRM | 5 |
| PUF60 | RRM,RRM,RRM | 5 |
| PUM1 | other | 5 |
| RALY | RRM | 3 |
| RBFOX2 | RRM | 1 |
| RBFOX3 | RRM | 1 |
| RBM15B | RRM,RRM,RRM | 5 |
| RBM22 | ZNF,RRM | 3 |
| RBM23 | RRM,RRM | 3 |
| RBM24 | RRM | 3 |
| RBM25 | RRM | 4 |
| RBM4 | RRM,RRM,ZNF | 5 |
| RBM41 | RRM | 5 |
| RBM45 | RRM,RRM,RRM | 5 |
| RBM47 | RRM,RRM,RRM | 4 |
| RBM4B | RRM,RRM,ZNF | 4 |
| RBM6 | RRM,RRM,ZNF,ZNF | 5 |

| | | |
|---|---|---|
| RBMS2 | RRM,RRM | 4 |
| RBMS3 | RRM,RRM | 3 |
| RC3H1 | other,ZNF | 5 |
| SF1 | KH,ZNF | 4 |
| SFPQ | RRM,RRM | 5 |
| SNRPA | RRM,RRM | 3 |
| SRSF10 | RRM | 2 |
| SRSF11 | RRM | 1 |
| SRSF2 | RRM | 3 |
| SRSF4 | RRM,RRM | 1 |
| SRSF5 | RRM,RRM | 2 |
| SRSF8 | RRM | 5 |
| SRSF9 | RRM,RRM | 2 |
| TAF15 | RRM,other | 3 |
| TARDBP | RRM,RRM | 3 |
| TIA1 | RRM,RRM,RRM | 5 |
| TRA2A | RRM | 5 |
| TRNAU1AP | RRM,RRM | 5 |
| UNK | ZNF,ZNF,ZNF,ZNF,ZNF | 4 |
| ZCRB1 | RRM,ZNF | 3 |
| ZFP36 | ZNF,ZNF | 4 |
| ZNF326 | ZNF,ZNF | 5 |

**Supplementary Table S2.** Error reductions and correlations for variant models

| | |
|---|---|
| **RBP** | name of RNA-binding protein |
| **rbp_conc** | Experimental RBP concentrations of the RBNS samples used in this run (nM) |
| **err_initial** | model error right after PSAM initialization. Comma-separated list, corresponding to rbp_conc |
| **corr_initial** | 6-mer R value correlation values between RBNS experimental data and RBPamp model predictions. |
| **err_nostruct** | model error after first round of SGD optimization (ignoring RNA secondary structure) |
| **corr_nostruct** | 6-mer correlation after first round of SGD optimization (ignoring RNA secondary structure) |
| **err_final** | model error after second round of SGD optimization (considerring RNA secondary structure) |
| **corr_final** | 6-mer correlation after second round of SGD optimization (considerring RNA secondary structure) |

| **RBP** | **rbp_conc** | **err_initial** | **corr_initial** | **err_nostruct** | **corr_nostruct** | **err_final** | **corr_final** |
|---|---|---|---|---|---|---|---|
| BOLL | 5.0,20.0,80.0,320.0 | 0.106,0.106,0.091,0.081 | 0.831,0.828,0.753,0.854 | 0.034,0.031,0.038,0.031 | 0.874,0.923,0.910,0.925 | 0.030,0.027,0.034,0.028 | 0.889,0.932,0.922,0.932 |
| CELF1 | 16.0,64.0,130.0,500.0 | 0.030,0.024,0.007,0.024 | 0.884,0.889,0.883,0.866 | 0.003,0.008,0.008,0.012 | 0.946,0.963,0.960,0.921 | 0.003,0.008,0.008,0.012 | 0.946,0.963,0.961,0.921 |
| CNOT4 | 5.0,20.0,80.0,320.0 | 0.012,0.018,0.017,0.015 | 0.667,0.598,0.468,0.508 | 0.003,0.004,0.003,0.003 | 0.932,0.925,0.950,0.926 | 0.003,0.004,0.003,0.003 | 0.922,0.921,0.943,0.928 |
| CPEB1 | 20.0,80.0,160.0,320.0 | 0.078,0.013,0.018,0.012 | 0.636,0.607,0.480,0.545 | 0.012,0.018,0.029,0.012 | 0.572,0.454,0.778,0.510 | 0.013,0.018,0.028,0.012 | 0.562,0.444,0.789,0.501 |
| DAZ3 | 5.0,20.0,80.0,320.0 | 0.042,0.055,0.064,0.040 | 0.844,0.825,0.806,0.789 | 0.010,0.008,0.008,0.012 | 0.962,0.967,0.965,0.942 | 0.007,0.006,0.006,0.009 | 0.971,0.975,0.974,0.956 |
| DAZAP1 | 5.0,20.0,80.0,320.0 | 0.029,0.023,0.014,0.006 | 0.791,0.822,0.862,0.856 | 0.004,0.003,0.004,0.004 | 0.965,0.970,0.959,0.910 | 0.004,0.003,0.004,0.004 | 0.969,0.972,0.959,0.915 |
| EIF4G2 | 20.0,80.0,320.0,1300.0 | 0.011,0.005,0.004,0.004 | 0.605,0.720,0.593,0.403 | 0.002,0.001,0.005,0.002 | 0.816,0.905,0.836,0.913 | 0.001,0.001,0.005,0.002 | 0.834,0.910,0.854,0.893 |
| ELAVL4 | 5.0,20.0,80.0,320.0 | 0.056,0.044,0.032,0. | 0.928,0.906,0.929,0.9 | 0.017,0.020,0.014,0.014 | 0.928,0.948,0.951,0.932 | 0.016,0.018,0.01 | 0.927,0.950,0.953,0.93 |

| | | | | | | | |
|---|---|---|---|---|---|---|---|
| | | 029 | 03 | | | 3,0.013 | 4 |
| ESRP1 | 5.0,80.0,320.0,1300.0 | 0.025,0.041,0.019,0.023 | 0.814,0.607,0.761,0.697 | 0.010,0.010,0.009,0.011 | 0.902,0.870,0.886,0.910 | 0.010,0.010,0.009,0.011 | 0.901,0.871,0.885,0.911 |
| EWSR1 | 5.0,20.0,80.0,320.0 | 0.005,0.006,0.006,0.024 | 0.694,0.552,0.736,0.418 | 0.004,0.003,0.005,0.008 | 0.725,0.806,0.787,0.847 | 0.004,0.003,0.005,0.008 | 0.722,0.804,0.785,0.850 |
| FUBP1 | 5.0,20.0,80.0,320.0 | 0.007,0.009,0.007,0.010 | 0.693,0.722,0.544,0.730 | 0.003,0.004,0.002,0.003 | 0.898,0.763,0.942,0.919 | 0.002,0.004,0.002,0.003 | 0.903,0.767,0.952,0.928 |
| FUBP3 | 5.0,20.0,80.0,320.0 | 0.037,0.033,0.081,0.019 | 0.930,0.917,0.835,0.919 | 0.024,0.008,0.022,0.017 | 0.936,0.959,0.950,0.942 | 0.020,0.007,0.020,0.014 | 0.950,0.965,0.955,0.951 |
| FUS | 20.0,80.0,320.0,1300.0 | 0.021,0.046,0.029,0.021 | 0.456,0.435,0.260,0.324 | 0.006,0.010,0.007,0.015 | 0.844,0.894,0.839,0.642 | 0.006,0.010,0.007,0.015 | 0.843,0.894,0.838,0.640 |
| A1CF | 5.0,20.0,80.0,320.0 | 0.084,0.055,0.035,0.036 | 0.748,0.819,0.840,0.851 | 0.041,0.027,0.019,0.018 | 0.854,0.893,0.903,0.921 | 0.029,0.018,0.012,0.012 | 0.893,0.924,0.935,0.952 |
| HNRNPA0 | 20.0,80.0,320.0,1300.0 | 0.020,0.022,0.010,0.020 | 0.913,0.901,0.895,0.711 | 0.005,0.008,0.009,0.010 | 0.946,0.962,0.955,0.849 | 0.004,0.007,0.008,0.009 | 0.951,0.967,0.961,0.867 |
| HNRNPA1 | 160 | 0.022 | 0.74 | 0.004 | 0.961 | 0.004 | 0.964 |
| HNRNPA2B1 | 5.0,20.0,80.0,320.0 | 0.024,0.023,0.010,0.025 | 0.299,0.416,0.240,0.421 | 0.005,0.012,0.011,0.011 | 0.730,0.714,0.779,0.784 | 0.003,0.008,0.007,0.008 | 0.829,0.830,0.869,0.856 |
| HNRNPC | 20.0,80.0,320.0,1300.0 | 0.017,0.012,0.001,0.003 | 0.887,0.828,0.816,0.076 | 0.003,0.001,0.009,0.009 | 0.080,0.828,0.906,0.845 | 0.003,0.001,0.009,0.009 | 0.083,0.827,0.907,0.846 |
| HNRNPCL1 | 160.0,320.0,1300.0 | 0.092,0.029,0.019 | 0.769,0.839,0.735 | 0.088,0.018,0.023 | 0.775,0.735,0.837 | 0.065,0.021,0.040 | 0.772,0.742,0.839 |
| HNRNPD | 20.0,80.0,320.0,1300.0 | 0.005,0.006,0.012,0.009 | 0.865,0.860,0.837,0.815 | 0.002,0.003,0.005,0.008 | 0.953,0.929,0.892,0.894 | 0.002,0.003,0.005,0.008 | 0.952,0.928,0.892,0.893 |
| HNRNPDL | 20.0,80.0,320.0,1300.0 | 0.013,0.014,0.008,0.007 | 0.832,0.772,0.726,0.462 | 0.006,0.006,0.008,0.009 | 0.571,0.797,0.885,0.841 | 0.006,0.006,0.008,0.009 | 0.568,0.796,0.886,0.839 |
| HNRNPF | 20.0,80.0,320.0,1300.0 | 0.023,0.018,0.029,0.030 | 0.872,0.821,0.716,0.645 | 0.015,0.017,0.011,0.010 | 0.809,0.828,0.879,0.919 | 0.015,0.016,0.011,0.011 | 0.810,0.828,0.879,0.919 |
| HNRNPH2 | 5.0,20.0,8 | 0.023,0.02 | 0.822,0.80 | 0.012,0.010,0 | 0.912,0.905,0. | 0.009,0. | 0.927,0.926 |

| | | | | | | | |
|---|---|---|---|---|---|---|---|
| | 0.0,320.0 | 1,0.033,0.037 | 0,0.769,0.764 | .016,0.018 | 884,0.873 | 008,0.012,0.014 | ,0.912,0.902 |
| HNRNPK | 5.0,20.0,80.0,320.0 | 0.024,0.104,0.027,0.020 | 0.668,0.525,0.784,0.820 | 0.013,0.046,0.011,0.022 | 0.829,0.801,0.916,0.806 | 0.016,0.030,0.008,0.011 | 0.802,0.864,0.940,0.904 |
| HNRNPL | 20.0,80.0,320.0,1300.0 | 0.035,0.045,0.033,0.023 | 0.886,0.872,0.884,0.844 | 0.012,0.011,0.009,0.006 | 0.962,0.966,0.965,0.953 | 0.010,0.010,0.008,0.005 | 0.962,0.963,0.965,0.957 |
| IGF2BP1 | 20.0,80.0,320.0,1300.0 | 0.037,0.030,0.033,0.016 | 0.748,0.694,0.778,0.575 | 0.005,0.010,0.017,0.015 | 0.894,0.901,0.884,0.887 | 0.005,0.010,0.018,0.016 | 0.893,0.899,0.882,0.884 |
| IGF2BP2 | 20.0,80.0,320.0,1300.0 | 0.028,0.024,0.018,0.015 | 0.753,0.658,0.721,0.716 | 0.008,0.009,0.012,0.010 | 0.864,0.833,0.842,0.913 | 0.008,0.009,0.012,0.010 | 0.863,0.834,0.843,0.913 |
| ILF2 | 20.0,80.0,320.0,1300.0 | 0.016,0.012,0.007,0.006 | 0.724,0.677,0.644,0.356 | 0.004,0.003,0.006,0.005 | 0.555,0.865,0.903,0.884 | 0.004,0.002,0.005,0.003 | 0.562,0.883,0.925,0.914 |
| KHDRBS2 | 20.0,320.0 | 0.050,0.036 | 0.929,0.924 | 0.016,0.013 | 0.958,0.959 | 0.015,0.009 | 0.960,0.968 |
| KHDRBS3 | 5.0,80.0,320.0,1300.0 | 0.034,0.031,0.022,0.005 | 0.827,0.827,0.718,-0.621 | 0.005,0.014,0.012,0.013 | -0.616,0.912,0.914,0.830 | 0.005,0.013,0.011,0.012 | -0.609,0.915,0.921,0.837 |
| KHSRP | 5.0,20.0,80.0,320.0 | 0.009,0.011,0.008,0.005 | 0.677,0.750,0.487,0.609 | 0.003,0.005,0.002,0.004 | 0.891,0.742,0.856,0.906 | 0.004,0.005,0.002,0.003 | 0.881,0.709,0.844,0.923 |
| MBNL1 | 365.0,1090.0,3280.0,9800.0 | 0.001,0.042,0.032,0.039 | 0.822,0.900,0.865,0.806 | 0.001,0.014,0.013,0.016 | 0.855,0.957,0.937,0.912 | 0.001,0.014,0.011,0.014 | 0.833,0.960,0.943,0.920 |
| MSI1 | 8.0,16.0,64.0,256.0 | 0.003,0.008,0.012,0.023 | 0.920,0.939,0.928,0.910 | 0.001,0.003,0.004,0.009 | 0.960,0.974,0.970,0.962 | 0.001,0.003,0.004,0.008 | 0.960,0.974,0.970,0.962 |
| NOVA1 | 5.0,20.0,80.0,320.0 | 0.126,0.123,0.106,0.074 | 0.791,0.767,0.784,0.740 | 0.045,0.039,0.030,0.019 | 0.927,0.925,0.942,0.939 | 0.038,0.032,0.024,0.016 | 0.935,0.939,0.954,0.949 |
| NUPL2 | 20.0,80.0,320.0,1300.0 | 0.050,0.033,0.024,0.026 | 0.813,0.860,0.894,0.879 | 0.020,0.015,0.012,0.012 | 0.916,0.933,0.945,0.940 | 0.018,0.013,0.010,0.010 | 0.924,0.940,0.951,0.944 |
| PABPN1L | 20.0,80.0,320.0,1300.0 | 0.005,0.005,0.004,0.007 | 0.747,0.670,0.754,0.334 | 0.004,0.005,0.003,0.007 | 0.695,0.766,0.769,0.373 | 0.004,0.004,0.003,0.006 | 0.704,0.777,0.784,0.399 |
| PCBP1 | 20.0,80.0,320.0,130 | 0.048,0.044,0.041,0. | 0.402,0.767,0.911,0.9 | 0.031,0.021,0 | 0.646,0.879,0. | 0.031,0.022,0.02 | 0.650,0.877,0.939,0.95 |

| | | | | | | | |
|---|---|---|---|---|---|---|---|
| | 0.0 | 042 | 38 | .027,0.027 | 940,0.954 | 7,0.026 | 4 |
| PCBP2 | 20.0,80.0,320.0,1300.0 | 0.068,0.062,0.198,0.071 | 0.264,0.628,0.902,0.893 | 0.064,0.048,0.092,0.037 | 0.361,0.722,0.941,0.937 | 0.062,0.045,0.052,0.025 | 0.382,0.741,0.956,0.955 |
| PCBP4 | 160.0,320.0,1300.0 | 0.030,0.025,0.017 | 0.220,0.797,0.702 | 0.024,0.009,0.005 | 0.449,0.936,0.931 | 0.023,0.006,0.004 | 0.497,0.958,0.943 |
| PRR3 | 5.0,20.0,80.0,320.0 | 0.231,0.155,0.091,0.064 | 0.864,0.864,0.861,0.836 | 0.080,0.050,0.031,0.019 | 0.919,0.936,0.935,0.942 | 0.078,0.052,0.031,0.020 | 0.920,0.936,0.937,0.942 |
| PTBP3 | 20.0,80.0,320.0,1300.0 | 0.029,0.026,0.005,0.015 | 0.853,0.843,0.834,0.863 | 0.002,0.007,0.009,0.007 | 0.914,0.960,0.939,0.933 | 0.002,0.007,0.009,0.007 | 0.915,0.959,0.938,0.934 |
| PUF60 | 20.0,80.0,320.0,1300.0 | 0.030,0.038,0.030,0.002 | 0.622,0.585,0.777,0.602 | 0.002,0.021,0.026,0.020 | 0.703,0.753,0.742,0.862 | 0.002,0.020,0.024,0.017 | 0.714,0.776,0.766,0.885 |
| PUM1 | 5.0,20.0,80.0,320.0 | 0.047,0.038,0.024,0.017 | 0.823,0.824,0.861,0.880 | 0.011,0.008,0.007,0.007 | 0.955,0.958,0.958,0.943 | 0.010,0.007,0.006,0.007 | 0.960,0.963,0.964,0.951 |
| RALY | 5.0,20.0,80.0,1300.0 | 0.026,0.028,0.025,0.026 | 0.791,0.854,0.890,0.534 | 0.024,0.019,0.018,0.017 | 0.598,0.831,0.886,0.911 | 0.021,0.016,0.014,0.015 | 0.642,0.860,0.903,0.928 |
| RBFOX2 | 121.0,365.0,1100.0,3300.0 | 0.020,0.008,0.033,0.030 | 0.922,0.898,0.744,0.693 | 0.008,0.013,0.024,0.023 | 0.896,0.935,0.799,0.749 | 0.006,0.012,0.024,0.024 | 0.906,0.938,0.792,0.737 |
| RBFOX3 | 5.0,20.0,80.0,320.0 | 0.245,0.188,0.113,0.131 | 0.828,0.840,0.797,0.659 | 0.123,0.096,0.073,0.101 | 0.852,0.871,0.845,0.738 | 0.092,0.068,0.057,0.090 | 0.878,0.897,0.872,0.768 |
| RBM15B | 20.0,80.0,320.0,1300.0 | 0.041,0.020,0.024,0.021 | 0.587,0.614,0.806,0.906 | 0.026,0.015,0.017,0.011 | 0.714,0.699,0.848,0.924 | 0.026,0.015,0.018,0.011 | 0.713,0.698,0.847,0.924 |
| RBM22 | 20.0,80.0,320.0,1300.0 | 0.084,0.009,0.005,0.015 | 0.779,0.708,0.315,-0.095 | 0.004,0.015,0.002,0.012 | 0.407,0.078,0.841,0.864 | 0.004,0.015,0.002,0.013 | 0.406,0.077,0.840,0.857 |
| RBM23 | 5.0,20.0,80.0,320.0 | 0.013,0.011,0.011,0.010 | 0.451,0.393,0.292,0.328 | 0.006,0.005,0.006,0.005 | 0.701,0.716,0.782,0.747 | 0.006,0.005,0.006,0.005 | 0.732,0.748,0.803,0.775 |
| RBM24 | 5.0,20.0,80.0,320.0 | 0.025,0.081,0.080,0.051 | 0.641,0.649,0.612,0.671 | 0.006,0.017,0.014,0.011 | 0.931,0.947,0.947,0.941 | 0.008,0.023,0.017,0.014 | 0.915,0.932,0.934,0.928 |
| RBM25 | 20.0,80.0,320.0,130 | 0.013,0.011,0.011,0. | 0.622,0.649,0.505,0.5 | 0.006,0.005,0.005,0.003 | 0.750,0.871,0.870,0.841 | 0.005,0.005,0.00 | 0.792,0.885,0.881,0.86 |

| | | | | | | | |
|---|---|---|---|---|---|---|---|
| | 0.0 | 006 | 99 | | | 4,0.003 | 2 |
| RBM4 | 5.0,20.0,80.0,320.0 | 0.034,0.038,0.033,0.029 | 0.587,0.628,0.688,0.689 | 0.014,0.014,0.014,0.016 | 0.846,0.874,0.883,0.826 | 0.009,0.009,0.010,0.013 | 0.907,0.922,0.914,0.866 |
| RBM41 | 5.0,20.0,80.0,320.0 | 0.124,0.127,0.124,0.097 | 0.718,0.744,0.704,0.694 | 0.037,0.041,0.031,0.026 | 0.934,0.939,0.941,0.937 | 0.034,0.035,0.026,0.021 | 0.943,0.949,0.953,0.949 |
| RBM45 | 5.0,20.0,80.0,320.0 | 0.074,0.087,0.067,0.119 | 0.764,0.802,0.705,0.780 | 0.038,0.043,0.036,0.060 | 0.873,0.902,0.844,0.893 | 0.032,0.036,0.030,0.047 | 0.892,0.920,0.872,0.920 |
| RBM47 | 20.0,60.0,240.0,960.0 | 0.010,0.007,0.018,0.011 | 0.881,0.890,0.862,0.895 | 0.004,0.004,0.010,0.007 | 0.938,0.940,0.916,0.934 | 0.004,0.003,0.010,0.007 | 0.940,0.942,0.922,0.937 |
| RBM4B | 5.0,80.0,320.0,1300.0 | 0.078,0.031,0.041,0.033 | 0.801,0.841,0.852,0.823 | 0.016,0.008,0.014,0.021 | 0.949,0.949,0.939,0.879 | 0.016,0.008,0.014,0.021 | 0.948,0.948,0.938,0.877 |
| RBM6 | 80.0,160.0,320.0 | 0.059,0.026,0.033 | 0.758,0.818,-0.253 | 0.014,0.033,0.009 | 0.937,-0.258,0.939 | 0.012,0.033,0.008 | 0.946,-0.241,0.942 |
| RBMS2 | 20.0,160.0,320.0 | 0.012,0.086,0.030 | 0.890,0.359,0.888 | 0.006,0.084,0.013 | 0.937,0.389,0.945 | 0.005,0.083,0.011 | 0.946,0.401,0.949 |
| RBMS3 | 20.0,80.0,320.0,1300.0 | 0.024,0.016,0.030,0.032 | 0.863,0.849,0.890,0.881 | 0.010,0.009,0.015,0.014 | 0.921,0.897,0.935,0.936 | 0.007,0.006,0.010,0.008 | 0.950,0.932,0.961,0.964 |
| RC3H1 | 80.0,160.0,1300.0 | 0.018,0.060,0.011 | 0.727,0.378,0.678 | 0.024,0.028,0.016 | 0.622,0.768,0.485 | 0.026,0.024,0.015 | 0.588,0.808,0.513 |
| SF1 | 80.0,320.0 | 0.024,0.035 | 0.905,0.905 | 0.015,0.020 | 0.936,0.937 | 0.014,0.018 | 0.934,0.937 |
| SFPQ | 5.0,20.0,320.0,1300.0 | 0.060,0.061,0.056,0.061 | 0.497,0.436,0.520,0.475 | 0.010,0.013,0.009,0.014 | 0.932,0.917,0.933,0.907 | 0.010,0.011,0.009,0.012 | 0.933,0.925,0.934,0.921 |
| SNRPA | 5.0,20.0,80.0,320.0 | 0.056,0.055,0.048,0.054 | 0.796,0.798,0.806,0.799 | 0.016,0.014,0.012,0.014 | 0.944,0.948,0.946,0.950 | 0.012,0.010,0.009,0.010 | 0.959,0.964,0.965,0.963 |
| SRSF10 | 80.0,320.0 | 0.004,0.001,0.002 | 0.000,0.634,0.561 | 0.001,0.003,0.634 | 0.780,0.000,0.000 | 0.001,0.002 | 0.705,0.837 |
| SRSF11 | 80.0,320.0,1300.0 | 0.014,0.018,0.012,0.005 | 0.576,0.501,0.455,0.241 | 0.005,0.007,0.006,0.813 | 0.839,0.851,0.000,0.000 | 0.005,0.007,0.006 | 0.812,0.838,0.850 |
| SRSF2 | 5.0,20.0,8 | 0.018,0.01 | 0.488,0.58 | 0.005,0.011,0 | 0.806,0.697,0. | 0.004,0. | 0.826,0.705 |

| | | | | | | | |
|---|---|---|---|---|---|---|---|
| | 0.0,320.0 | 3,0.009,0.017 | 9,0.566,0.545 | .008,0.011 | 778,0.744 | 011,0.007,0.010 | ,0.809,0.777 |
| SRSF4 | 20.0,80.0,320.0,1300.0 | 0.035,0.038,0.022,0.017 | 0.380,0.372,0.382,0.379 | 0.011,0.015,0.015,0.007 | 0.754,0.814,0.796,0.812 | 0.011,0.016,0.014,0.007 | 0.761,0.791,0.799,0.814 |
| SRSF5 | 20.0,80.0,320.0,1300.0 | 0.018,0.018,0.020,0.025 | 0.501,0.534,0.631,0.647 | 0.007,0.009,0.009,0.011 | 0.833,0.797,0.846,0.853 | 0.007,0.009,0.009,0.011 | 0.834,0.799,0.847,0.854 |
| SRSF8 | 20.0,80.0,320.0,1300.0 | 0.027,0.023,0.032,0.024 | 0.470,0.624,0.659,0.549 | 0.015,0.012,0.011,0.016 | 0.733,0.793,0.834,0.832 | 0.012,0.009,0.008,0.011 | 0.807,0.857,0.884,0.894 |
| SRSF9 | 20.0,80.0,320.0,1300.0 | 0.012,0.042,0.006,0.005 | 0.777,0.611,0.747,0.679 | 0.003,0.003,0.011,0.010 | 0.792,0.872,0.816,0.928 | 0.003,0.003,0.008,0.009 | 0.834,0.903,0.865,0.926 |
| TAF15 | 20.0,80.0,320.0,1300.0 | 0.014,0.016,0.015,0.016 | 0.425,0.474,0.387,0.251 | 0.006,0.009,0.012,0.014 | 0.784,0.766,0.597,0.423 | 0.007,0.010,0.012,0.015 | 0.755,0.733,0.560,0.384 |
| TARDBP | 320.0,1300.0 | 0.021,0.004 | 0.785,0.729 | 0.003,0.012 | 0.808,0.871 | 0.002,0.007 | 0.881,0.928 |
| TIA1 | 5.0,20.0,320.0,1300.0 | 0.003,0.003,0.003,0.003 | 0.816,0.489,0.538,0.590 | 0.001,0.002,0.002,0.002 | 0.822,0.593,0.687,0.833 | 0.002,0.003,0.002,0.002 | 0.802,0.495,0.693,0.834 |
| TRA2A | 5.0,20.0,80.0,320.0 | 0.007,0.013,0.024,0.033 | 0.893,0.900,0.877,0.839 | 0.002,0.006,0.010,0.014 | 0.951,0.939,0.938,0.920 | 0.002,0.006,0.008,0.012 | 0.954,0.941,0.945,0.935 |
| TRNAU1AP | 5.0,20.0 | 0.053,0.061 | 0.799,0.835 | 0.020,0.024 | 0.900,0.921 | 0.018,0.021 | 0.909,0.928 |
| UNK | 20.0,80.0,320.0,1300.0 | 0.036,0.036,0.020,0.025 | 0.898,0.896,0.913,0.880 | 0.014,0.018,0.010,0.012 | 0.945,0.935,0.953,0.942 | 0.012,0.015,0.009,0.010 | 0.951,0.942,0.959,0.949 |
| ZCRB1 | 5.0,20.0,80.0,320.0 | 0.008,0.006,0.006,0.005 | 0.732,0.791,0.737,0.713 | 0.004,0.003,0.003,0.002 | 0.883,0.907,0.889,0.878 | 0.003,0.003,0.002,0.002 | 0.904,0.923,0.904,0.909 |
| ZFP36 | 20.0,80.0,320.0,1300.0 | 0.058,0.078,0.070,0.049 | 0.884,0.886,0.873,0.872 | 0.018,0.019,0.015,0.013 | 0.917,0.925,0.930,0.931 | 0.017,0.017,0.013,0.012 | 0.920,0.931,0.934,0.936 |
| ZNF326 | 5.0,20.0 | 0.015,0.019 | 0.513,0.413 | 0.007,0.008 | 0.835,0.825 | 0.007,0.008 | 0.814,0.811 |

**Supplementary Table S3.** AFP calibration results.

| | |
|---|---|
| **PSAM_id** | RBP name, dot, index of associated PSAM |
| **PSAM_motif** | Cognate sequence of PSAM. Upper-case indicates positions with high nucleotide discrimination |
| **PSAM_GC** | G+C fraction of motif (weight assigned by the PSAM) |
| **AFP_width** | Optimal size of accessibility footprint |
| **AFP_start** | Optimal start position of accessibility footprint, relative to the PSAM start |
| **AFP_scale** | Optimal scaling parameter *a* of accessibility footprint |
| **AFP_PSAM_overlap** | Fraction of AFP that overlaps with the PSAM |
| **rel_profile_err** | Relative reduction in $P_{unpaired}$ profile error by using the scaled AFP, compared to ignoring structure |
| **QC_pass** | AFP reduces profile error by at least 10% and has more than 50% overlap with PSAM? (T or F) |

| **PSAM_id** | **PSAM_motif** | **PSAM_GC** | **AFP_width** | **AFP_start** | **AFP_scale** | **AFP_PSAM_overlap** | **rel_profile_err** | **QC_pass** |
|---|---|---|---|---|---|---|---|---|
| BOLL.0 | cuuUuuuuugc | 0.43 | 11 | -3 | 0.0836 | 0.73 | 0.16 | True |
| BOLL.1 | guuugugUUau | 0.40 | 10 | -1 | 0.0798 | 0.90 | 0.11 | True |
| BOLL.2 | cuuuuaauUUu | 0.30 | 12 | -1 | 0.0615 | 0.92 | 0.13 | True |
| BOLL.3 | uugUUauuUgu | 0.30 | 14 | -4 | 0.1220 | 0.71 | 0.05 | True |
| BOLL.4 | ccuuuuUUauu | 0.36 | 10 | 0 | 0.0629 | 1.00 | 0.10 | True |
| CELF1.0 | uugUauGUuuu | 0.35 | 14 | -3 | 0.0998 | 0.79 | 0.26 | True |
| CELF1.1 | uaaugUuuuuu | 0.29 | 10 | -2 | 0.0875 | 0.80 | 0.55 | True |
| CELF1.2 | cuGucuugUcc | 0.48 | 10 | -1 | 0.1064 | 0.90 | 0.26 | True |
| CELF1.3 | uugUccgugug | 0.47 | 13 | -2 | 0.1639 | 0.85 | 0.16 | True |
| CELF1.4 | guaaaugUaua | 0.35 | 20 | -3 | 0.0482 | 0.70 | 0.80 | True |
| CNOT4.0 | caaacAcauuc | 0.41 | 5 | -5 | 0.0249 | 0.00 | 0.79 | False |
| CNOT4.1 | ggccggacagc | 0.61 | 16 | -5 | 0.0896 | 0.69 | 0.07 | True |
| CNOT4.2 | ggggcucccgc | 0.59 | 13 | -3 | 0.1135 | 0.77 | 0.06 | True |
| CPEB1.0 | auuuUUUuaaa | 0.25 | 16 | -5 | 0.0855 | 0.69 | 0.20 | True |
| CPEB1.1 | guuuauUUUaa | 0.37 | 5 | 8 | 0.2226 | 0.60 | 0.12 | True |
| CPEB1.2 | uUUUAAaauaa | 0.32 | 5 | -4 | 0.0416 | 0.20 | 0.75 | False |

| | | | | | | | | |
|---|---|---|---|---|---|---|---|---|
| DAZ3.0 | gaGUUauguuu | 0.39 | 11 | 0 | 0.1487 | 1.00 | 0.12 | True |
| DAZ3.1 | ccaagUUaucg | 0.44 | 5 | 3 | 0.1323 | 1.00 | 0.09 | True |
| DAZ3.2 | gcgUUuaaauc | 0.40 | 5 | 0 | 0.1542 | 1.00 | 0.35 | True |
| DAZ3.3 | cuuuuuuuauu | 0.34 | 17 | -5 | 0.0559 | 0.71 | 0.38 | True |
| DAZ3.4 | auccacguuuc | 0.53 | 8 | 4 | 0.1328 | 0.88 | 0.10 | True |
| DAZAP1.0 | aUauauauAg | 0.33 | 5 | 9 | 0.0367 | 0.20 | 0.64 | False |
| DAZAP1.1 | uUagcauagg | 0.44 | 15 | -5 | 0.0470 | 0.67 | 0.16 | True |
| DAZAP1.2 | cauaaguagg | 0.46 | 5 | 9 | 0.0350 | 0.20 | 0.67 | False |
| DAZAP1.3 | aauauaguuu | 0.35 | 20 | -2 | 0.0700 | 0.60 | 0.45 | True |
| EIF4G2.0 | ugcguugcaag | 0.49 | 5 | 2 | 0.1650 | 1.00 | 0.16 | True |
| EIF4G2.1 | gccgcggccgg | 0.58 | 12 | -3 | 0.1201 | 0.75 | 0.21 | True |
| EIF4G2.2 | cgugcgagggg | 0.59 | 10 | 0 | 0.1094 | 1.00 | 0.36 | True |
| EIF4G2.3 | cuggccgucgg | 0.56 | 12 | -5 | 0.1743 | 0.58 | 0.26 | True |
| ELAVL4.0 | uuaUuauUUu | 0.16 | 5 | -5 | 0.0680 | 0.00 | 0.34 | False |
| ELAVL4.1 | uuuUaagUUu | 0.20 | 5 | 6 | 0.0396 | 0.80 | 0.82 | True |
| ELAVL4.2 | cauUuuuuUu | 0.24 | 6 | -5 | 0.0603 | 0.17 | 0.14 | False |
| ELAVL4.3 | cuuuUuUaau | 0.26 | 20 | -1 | 0.1212 | 0.55 | 0.51 | True |
| ELAVL4.4 | auauuUuuuu | 0.18 | 14 | -4 | 0.0876 | 0.71 | 0.23 | True |
| ESRP1.0 | cgaucgucggg | 0.55 | 6 | 5 | 0.0887 | 1.00 | 0.15 | True |
| ESRP1.1 | ugggUgauggc | 0.52 | 13 | -3 | 0.1712 | 0.77 | 0.16 | True |
| ESRP1.2 | ggggaagggga | 0.58 | 10 | -3 | 0.0000 | 0.70 | 1.00 | False |
| ESRP1.3 | auugggggggg | 0.52 | 13 | 0 | 0.1727 | 0.85 | 0.07 | True |
| EWSR1.0 | uacccGagaau | 0.50 | 5 | 6 | 0.0659 | 1.00 | 0.62 | True |
| EWSR1.1 | cgaCacccguc | 0.66 | 5 | 8 | 0.0643 | 0.60 | 0.64 | True |
| EWSR1.2 | ggggugcccgg | 0.54 | 15 | -5 | 0.1091 | 0.67 | 0.36 | True |
| EWSR1.3 | uuggcgcccgg | 0.63 | 9 | 2 | 0.1151 | 1.00 | 0.52 | True |
| FUBP1.0 | uuuuauguauu | 0.19 | 5 | -5 | 0.0920 | 0.00 | 0.59 | False |
| FUBP1.1 | guauguauuuu | 0.31 | 7 | -5 | 0.0692 | 0.29 | 0.35 | False |
| FUBP1.2 | uaaUguuuuuu | 0.38 | 16 | -4 | 0.0364 | 0.75 | 0.63 | True |
| FUBP1.3 | cuuuauugugg | 0.51 | 16 | -5 | 0.1853 | 0.69 | 0.15 | True |
| FUBP1.4 | cuuuugugugg | 0.50 | 15 | -4 | 0.0959 | 0.73 | 0.11 | True |
| FUBP3.0 | uaauauauaau | 0.24 | 6 | 10 | 0.1325 | 0.17 | 0.24 | False |

| | | | | | | | | |
|---|---|---|---|---|---|---|---|---|
| FUBP3.1 | auuuaUguaau | 0.23 | 18 | -5 | 0.1164 | 0.72 | 0.37 | True |
| FUBP3.2 | auaaauuaaau | 0.11 | 5 | 11 | 0.1198 | 0.00 | 0.44 | False |
| FUBP3.3 | aaauaauauaa | 0.17 | 5 | 11 | 0.1073 | 0.00 | 0.37 | False |
| FUBP3.4 | uuuauuauaaa | 0.22 | 19 | -5 | 0.1274 | 0.74 | 0.36 | True |
| FUS.0 | gcacggaggc | 0.55 | 12 | -3 | 0.2093 | 0.75 | 0.18 | True |
| A1CF.0 | auuAAUuaggg | 0.41 | 11 | -1 | 0.1024 | 0.91 | 0.28 | True |
| A1CF.1 | aaaUuuaauag | 0.30 | 12 | -1 | 0.0797 | 0.92 | 0.19 | True |
| A1CF.2 | cuaauauuAau | 0.28 | 13 | -5 | 0.0966 | 0.62 | 0.27 | True |
| A1CF.3 | aAUcAuaauaa | 0.33 | 5 | -1 | 0.1797 | 0.80 | 0.07 | True |
| A1CF.4 | gucauuAAUcg | 0.39 | 10 | -1 | 0.1390 | 0.90 | 0.30 | True |
| HNRNPA0.0 | auuuaUAguaa | 0.31 | 15 | -2 | 0.0746 | 0.87 | 0.47 | True |
| HNRNPA0.1 | cagcacuaggc | 0.58 | 5 | 4 | 0.1456 | 1.00 | 0.74 | True |
| HNRNPA0.2 | aauUAGguaaa | 0.34 | 5 | 1 | 0.0741 | 1.00 | 0.53 | True |
| HNRNPA0.3 | cauaggccuag | 0.49 | 5 | -5 | 0.0560 | 0.00 | 0.64 | False |
| HNRNPA0.4 | gauAGaguagc | 0.48 | 15 | -2 | 0.0473 | 0.87 | 0.55 | True |
| HNRNPA1.0 | auuAgguagug | 0.41 | 12 | -1 | 0.0738 | 0.92 | 0.52 | True |
| HNRNPA1.1 | auuuAGcagaa | 0.38 | 13 | -4 | 0.0466 | 0.69 | 0.63 | True |
| HNRNPA1.2 | gcauuagggua | 0.50 | 10 | -1 | 0.0897 | 0.90 | 0.30 | True |
| HNRNPA1.3 | cacaggguagc | 0.57 | 7 | 0 | 0.0948 | 1.00 | 0.21 | True |
| HNRNPA1.4 | AugauCGucgg | 0.57 | 7 | 0 | 0.1450 | 1.00 | 0.26 | True |
| HNRNPA2B1.0 | gggggggua | 0.61 | 5 | 2 | 0.0919 | 1.00 | 0.58 | True |
| HNRNPA2B1.1 | GAUCGucggc | 0.58 | 5 | -2 | 0.0471 | 0.60 | 0.65 | True |
| HNRNPA2B1.2 | agggggcagc | 0.52 | 11 | 4 | 0.1383 | 0.55 | 0.09 | True |
| HNRNPC.0 | UUUuaauUUUa | 0.22 | 6 | 6 | 0.1731 | 0.83 | 0.69 | True |

| | | | | | | | | |
|---|---|---|---|---|---|---|---|---|
| HNRNPC.1 | ccauuUUUUuu | 0.29 | 8 | 5 | 0.0673 | 0.75 | 0.30 | True |
| HNRNPCL 1.0 | uaUuUUuuuuu | 0.31 | 12 | -1 | 0.2673 | 0.92 | 0.11 | True |
| HNRNPCL 1.1 | gUUUUaauUUU | 0.23 | 5 | 7 | 0.2978 | 0.80 | 0.50 | True |
| HNRNPCL 1.2 | uaaUUUUUaaa | 0.38 | 5 | 0 | 0.0939 | 1.00 | 0.25 | True |
| HNRNPCL 1.3 | auauaUUUUag | 0.28 | 5 | -5 | 0.1251 | 0.00 | 0.37 | False |
| HNRNPD.0 | cauuuuAauau | 0.29 | 19 | -5 | 0.1627 | 0.74 | 0.24 | True |
| HNRNPD.1 | auaauauAuaa | 0.20 | 5 | -5 | 0.0774 | 0.00 | 0.54 | False |
| HNRNPD.2 | cuuuuauuuaa | 0.31 | 20 | -5 | 0.1636 | 0.75 | 0.22 | True |
| HNRNPD.3 | uuauuauuAua | 0.28 | 20 | -5 | 0.0969 | 0.75 | 0.34 | True |
| HNRNPD.4 | uauAauuaauu | 0.24 | 20 | -5 | 0.0957 | 0.75 | 0.38 | True |
| HNRNPDL. 0 | auuuauauuug | 0.18 | 20 | 1 | 0.1860 | 0.50 | 0.15 | False |
| HNRNPDL. 1 | aauaauauAaa | 0.24 | 20 | -5 | 0.1023 | 0.75 | 0.50 | True |
| HNRNPDL. 2 | auauAguAauu | 0.20 | 5 | 10 | 0.1112 | 0.20 | 0.30 | False |
| HNRNPDL. 3 | guaaguuuaaa | 0.28 | 20 | -5 | 0.0994 | 0.75 | 0.35 | True |
| HNRNPDL. 4 | aauUAauuaaa | 0.25 | 5 | 11 | 0.1011 | 0.00 | 0.39 | False |
| HNRNPF.0 | ggggaaggggc | 0.55 | 8 | 2 | 0.0813 | 1.00 | 0.28 | True |
| HNRNPF.1 | gaagagggugg | 0.49 | 5 | 5 | 0.2147 | 1.00 | 0.17 | True |
| HNRNPF.2 | aggggggguaa | 0.53 | 15 | -3 | 0.1740 | 0.80 | 0.14 | True |
| HNRNPF.3 | aaagggguaaa | 0.36 | 5 | 3 | 0.1702 | 1.00 | 0.39 | True |
| HNRNPH2. 0 | uagGgggGggc | 0.64 | 8 | 2 | 0.1861 | 1.00 | 0.09 | True |
| HNRNPH2. 1 | agaGGgaggua | 0.54 | 6 | 2 | 0.1311 | 1.00 | 0.53 | True |
| HNRNPH2. 2 | Gauggugggua | 0.56 | 5 | 2 | 0.1000 | 1.00 | 0.41 | True |
| HNRNPH2. 3 | aaagGGggaaa | 0.48 | 5 | 2 | 0.1343 | 1.00 | 0.63 | True |
| HNRNPH2. | aGgggagggcc | 0.59 | 9 | 1 | 0.1388 | 1.00 | 0.23 | True |

| | | | | | | | | |
|---|---|---|---|---|---|---|---|---|
| 4 | | | | | | | | |
| HNRNPK.0 | ccagcccaacc | 0.52 | 6 | -3 | 0.0791 | 0.50 | 0.32 | False |
| HNRNPK.1 | cugccccaaca | 0.50 | 5 | -3 | 0.0778 | 0.40 | 0.57 | False |
| HNRNPK.2 | ggccccaucca | 0.55 | 20 | -2 | 0.0884 | 0.65 | 0.33 | True |
| HNRNPK.3 | gccaccacccc | 0.55 | 11 | -5 | 0.0739 | 0.55 | 0.39 | True |
| HNRNPK.4 | cccaugcccca | 0.56 | 16 | -4 | 0.1613 | 0.75 | 0.18 | True |
| HNRNPL.0 | uauAcAUaaca | 0.28 | 18 | -5 | 0.1715 | 0.72 | 0.10 | True |
| HNRNPL.1 | uauacaCaaca | 0.35 | 19 | -5 | 0.1552 | 0.74 | 0.08 | True |
| HNRNPL.2 | caacAuacauc | 0.42 | 18 | -5 | 0.1438 | 0.72 | 0.07 | True |
| HNRNPL.3 | ucAuucacacc | 0.44 | 17 | -5 | 0.1576 | 0.71 | 0.11 | True |
| HNRNPL.4 | cccaccacaua | 0.48 | 20 | -5 | 0.1708 | 0.75 | 0.02 | True |
| IGF2BP1.0 | uacaauacAua | 0.34 | 20 | -5 | 0.1649 | 0.75 | 0.50 | True |
| IGF2BP1.1 | ccacucccccc | 0.50 | 20 | -5 | 0.2072 | 0.75 | 0.18 | True |
| IGF2BP1.2 | caaacaacaaa | 0.37 | 19 | -2 | 0.0968 | 0.68 | 0.38 | True |
| IGF2BP1.3 | ucAcacacAca | 0.46 | 20 | -5 | 0.1075 | 0.75 | 0.68 | True |
| IGF2BP2.0 | aauacAuuauc | 0.26 | 9 | 1 | 0.2074 | 1.00 | 0.69 | True |
| IGF2BP2.1 | cacauucauaa | 0.28 | 9 | -3 | 0.1274 | 0.67 | 0.59 | True |
| IGF2BP2.2 | caaCaacaacc | 0.40 | 5 | -5 | 0.0638 | 0.00 | 0.84 | False |
| IGF2BP2.3 | cgcaaaacaag | 0.50 | 17 | -5 | 0.1455 | 0.71 | 0.43 | True |
| IGF2BP2.4 | aacAcacAaua | 0.36 | 17 | -5 | 0.0962 | 0.71 | 0.18 | True |
| ILF2.0 | gggguaggggu | 0.57 | 5 | 0 | 0.1048 | 1.00 | 0.49 | True |
| ILF2.1 | aaugggguaaa | 0.33 | 5 | 3 | 0.0878 | 1.00 | 0.47 | True |
| ILF2.2 | ggugugggaaa | 0.46 | 5 | 2 | 0.1336 | 1.00 | 0.07 | True |
| ILF2.3 | cGAUCGucggg | 0.61 | 8 | 1 | 0.1107 | 1.00 | 0.24 | True |
| ILF2.4 | cuGggggggac | 0.57 | 6 | 2 | 0.1345 | 1.00 | 0.06 | True |
| KHDRBS2.0 | uuaaauUAAaa | 0.25 | 17 | -5 | 0.1657 | 0.71 | 0.05 | True |
| KHDRBS2.1 | auaUAAuacuu | 0.22 | 20 | -5 | 0.2083 | 0.75 | 0.09 | True |
| KHDRBS2.2 | cuaaUAAaaau | 0.34 | 18 | -3 | 0.1688 | 0.78 | 0.17 | True |
| KHDRBS2.3 | uaaauuaUAAa | 0.29 | 17 | -3 | 0.1506 | 0.82 | 0.10 | True |

| | | | | | | | | |
|---|---|---|---|---|---|---|---|---|
| KHDRBS2.4 | gaauAauUAAa | 0.28 | 17 | -3 | 0.2019 | 0.82 | 0.10 | True |
| KHDRBS3.0 | auUAUUUAuua | 0.06 | 6 | -3 | 0.1321 | 0.50 | 0.35 | False |
| KHDRBS3.1 | cuaUAAuauaA | 0.14 | 14 | 0 | 0.1822 | 0.79 | 0.05 | True |
| KHDRBS3.2 | cuaUaAaaaau | 0.30 | 20 | 1 | 0.1955 | 0.50 | 0.17 | False |
| KHDRBS3.3 | caaaaUAAaaa | 0.34 | 18 | -3 | 0.1033 | 0.78 | 0.31 | True |
| KHDRBS3.4 | aacgcauaauc | 0.51 | 18 | -5 | 0.0953 | 0.72 | 0.23 | True |
| KHSRP.0 | uuguaAuuuau | 0.17 | 5 | -1 | 0.1265 | 0.80 | 0.27 | True |
| KHSRP.1 | uaGuauauuau | 0.21 | 5 | -5 | 0.0858 | 0.00 | 0.51 | False |
| KHSRP.2 | uuaaauGuaua | 0.31 | 5 | -5 | 0.0591 | 0.00 | 0.48 | False |
| KHSRP.3 | uguauuauuaa | 0.28 | 7 | -5 | 0.0801 | 0.29 | 0.10 | False |
| KHSRP.4 | gggugcuuauu | 0.50 | 17 | -5 | 0.1743 | 0.71 | 0.11 | True |
| MBNL1.0 | cGCuucGCuuc | 0.48 | 17 | -4 | 0.3304 | 0.76 | 0.27 | True |
| MBNL1.1 | uuuGCuuuugC | 0.30 | 19 | -5 | 0.2904 | 0.74 | 0.52 | True |
| MBNL1.2 | aacgCGCuuaa | 0.39 | 20 | -5 | 0.2058 | 0.75 | 0.73 | True |
| MBNL1.3 | cacgCuGCuua | 0.46 | 8 | -4 | 0.0464 | 0.50 | 0.54 | False |
| MBNL1.4 | ccGCuuGCuug | 0.48 | 19 | -5 | 0.2602 | 0.74 | 0.57 | True |
| MSI1.0 | auuAguuAgua | 0.34 | 16 | -2 | 0.1223 | 0.81 | 0.31 | True |
| MSI1.1 | cuuagauUAGu | 0.40 | 20 | -2 | 0.1004 | 0.65 | 0.33 | True |
| MSI1.2 | uuuUAguauua | 0.25 | 19 | -4 | 0.1025 | 0.79 | 0.35 | True |
| MSI1.3 | uaUAgaaauaa | 0.34 | 8 | -4 | 0.0628 | 0.50 | 0.60 | False |
| MSI1.4 | aauauUAGauu | 0.28 | 6 | 7 | 0.0745 | 0.67 | 0.41 | True |
| NOVA1.0 | uCauuuCAuac | 0.41 | 12 | -2 | 0.1761 | 0.83 | 0.07 | True |
| NOVA1.1 | cucauuCAuua | 0.33 | 15 | -3 | 0.1767 | 0.80 | 0.06 | True |
| NOVA1.2 | acCAucaccgc | 0.56 | 15 | -5 | 0.1628 | 0.67 | 0.09 | True |
| NOVA1.3 | ucauCAUaaaa | 0.30 | 7 | -3 | 0.1792 | 0.57 | 0.15 | True |
| NUPL2.0 | acaaacaccca | 0.51 | 19 | -4 | 0.1282 | 0.79 | 0.29 | True |
| NUPL2.1 | gcaaaguaAag | 0.44 | 13 | 0 | 0.1290 | 0.85 | 0.32 | True |
| NUPL2.2 | accaAaaaaaa | 0.43 | 8 | 3 | 0.2256 | 1.00 | 0.46 | True |

| | | | | | | | | |
|---|---|---|---|---|---|---|---|---|
| NUPL2.3 | caAauaaaaag | 0.37 | 16 | -2 | 0.1244 | 0.81 | 0.24 | True |
| PABPN1L.0 | auaAAAaAaa | 0.27 | 19 | -5 | 0.1394 | 0.74 | 0.08 | True |
| PABPN1L.1 | gAAgAauAau | 0.28 | 14 | -3 | 0.3114 | 0.79 | 0.10 | True |
| PABPN1L.2 | aAAAuAuaaA | 0.19 | 18 | -5 | 0.2195 | 0.72 | 0.42 | True |
| PABPN1L.3 | aaauUAAaau | 0.15 | 5 | 7 | 0.1541 | 0.60 | 0.41 | True |
| PCBP1.0 | acCCcgGucgg | 0.71 | 20 | -1 | 0.0605 | 0.60 | 0.30 | True |
| PCBP1.1 | cccaagCCcag | 0.64 | 8 | 2 | 0.2301 | 1.00 | 0.35 | True |
| PCBP1.2 | gcccgcccccc | 0.68 | 8 | 1 | 0.1519 | 1.00 | 0.39 | True |
| PCBP1.3 | cccuuucCccu | 0.54 | 6 | 1 | 0.1627 | 1.00 | 0.13 | True |
| PCBP1.4 | cgcCcacccgc | 0.61 | 9 | 0 | 0.1288 | 1.00 | 0.49 | True |
| PCBP2.0 | acCCccgucgg | 0.72 | 6 | 0 | 0.4227 | 1.00 | 0.28 | True |
| PCBP2.1 | uaCCccccccg | 0.58 | 12 | -1 | 0.3373 | 0.92 | 0.04 | True |
| PCBP2.2 | cccacaacCCu | 0.50 | 14 | -2 | 0.3117 | 0.86 | 0.05 | True |
| PCBP2.3 | cuacCCCcacg | 0.56 | 7 | 1 | 0.3888 | 1.00 | 0.05 | True |
| PCBP4.0 | caauCCccccg | 0.53 | 7 | 2 | 0.4108 | 1.00 | 0.10 | True |
| PCBP4.1 | uCCuaauccug | 0.45 | 13 | -3 | 0.3629 | 0.77 | 0.32 | True |
| PCBP4.2 | uccuuUucCcc | 0.52 | 12 | -2 | 0.4241 | 0.83 | 0.02 | True |
| PCBP4.3 | gucccuuCCca | 0.50 | 10 | 1 | 0.4213 | 1.00 | 0.04 | True |
| PCBP4.4 | aucccggucgg | 0.59 | 7 | -1 | 0.3251 | 0.86 | 0.17 | True |
| PRR3.0 | aUuACCGgacu | 0.44 | 5 | 11 | 0.0470 | 0.00 | 0.40 | False |
| PRR3.1 | aUGAGCGgacc | 0.56 | 5 | -3 | 0.2606 | 0.40 | 0.22 | False |
| PRR3.2 | auAUaAgcggg | 0.46 | 5 | -3 | 0.1444 | 0.40 | 0.19 | False |
| PRR3.3 | aaauAUaaguc | 0.35 | 8 | -3 | 0.0752 | 0.63 | 0.47 | True |
| PTBP3.0 | uuuCUuucucu | 0.34 | 20 | -5 | 0.1645 | 0.75 | 0.19 | True |
| PTBP3.1 | cuCUuuucUuu | 0.35 | 19 | -5 | 0.1721 | 0.74 | 0.17 | True |
| PTBP3.2 | cuuuacuauau | 0.34 | 20 | -5 | 0.1750 | 0.75 | 0.17 | True |
| PTBP3.3 | cuucuaucuuu | 0.38 | 20 | -5 | 0.1614 | 0.75 | 0.27 | True |
| PTBP3.4 | ccuuCuuuuca | 0.36 | 14 | -2 | 0.1862 | 0.86 | 0.21 | True |
| PUF60.0 | guguuuuUuUu | 0.39 | 14 | -2 | 0.3023 | 0.86 | 0.17 | True |

| | | | | | | | | |
|---|---|---|---|---|---|---|---|---|
| PUF60.1 | ucuuuucuccc | 0.39 | 12 | -2 | 0.2211 | 0.83 | 0.32 | True |
| PUF60.2 | uuUcuuaUuug | 0.37 | 15 | -5 | 0.2489 | 0.67 | 0.25 | True |
| PUF60.3 | guucucuuuuc | 0.60 | 12 | -2 | 0.0767 | 0.83 | 1.15 | False |
| PUF60.4 | cuUcUuUcucu | 0.42 | 15 | -3 | 0.1343 | 0.80 | 0.82 | True |
| PUM1.0 | GGGagugUaag | 0.52 | 5 | 3 | 0.0850 | 1.00 | 0.71 | True |
| PUM1.1 | caUguauAuag | 0.37 | 12 | 0 | 0.0908 | 0.92 | 0.17 | True |
| PUM1.2 | guguaauauag | 0.29 | 5 | -1 | 0.0730 | 0.80 | 0.55 | True |
| PUM1.3 | cauaUgUaauu | 0.28 | 6 | 7 | 0.0977 | 0.67 | 0.20 | True |
| PUM1.4 | gguaaaUAaug | 0.36 | 20 | -2 | 0.0489 | 0.65 | 0.41 | True |
| RALY.0 | GGuuuaauUUU | 0.37 | 5 | 9 | 0.0458 | 0.40 | 0.47 | False |
| RALY.1 | uuuuuuuUacu | 0.36 | 11 | -4 | 0.0987 | 0.64 | 0.09 | True |
| RALY.2 | cUUuuucuugg | 0.36 | 12 | -3 | 0.1327 | 0.75 | 0.06 | True |
| RBFOX2.0 | uuuGCAugcau | 0.44 | 8 | 2 | 0.2706 | 1.00 | 0.12 | True |
| RBFOX3.0 | gcuGCAuGcau | 0.50 | 7 | 2 | 0.1565 | 1.00 | 0.07 | True |
| RBM15B.0 | auuaauuuuac | 0.25 | 5 | -4 | 0.2366 | 0.20 | 0.61 | False |
| RBM15B.1 | uuuuuuaauuu | 0.20 | 5 | -3 | 0.4384 | 0.40 | 0.29 | False |
| RBM15B.2 | uuuuUauuuau | 0.21 | 7 | -5 | 0.3303 | 0.29 | 0.27 | False |
| RBM15B.3 | aauuuuuuuuc | 0.31 | 15 | -3 | 0.2733 | 0.80 | 0.06 | True |
| RBM15B.4 | ucCuuuuuuac | 0.37 | 13 | -5 | 0.2721 | 0.62 | 0.16 | True |
| RBM22.0 | auUUaCCGgac | 0.48 | 5 | -3 | 0.2357 | 0.40 | 0.08 | False |
| RBM22.1 | auauUUACCaa | 0.41 | 5 | -5 | 0.0000 | 0.00 | 1.00 | False |
| RBM22.2 | aACCGGacuaa | 0.53 | 16 | -4 | 0.0540 | 0.75 | 0.86 | True |
| RBM23.0 | agccccccaa | 0.49 | 18 | -5 | 0.0407 | 0.72 | 0.41 | True |
| RBM23.1 | acgcccuGga | 0.56 | 5 | 3 | 0.0781 | 1.00 | 0.35 | True |
| RBM23.2 | GAUCGucgac | 0.58 | 8 | -3 | 0.0621 | 0.63 | 0.49 | True |
| RBM24.0 | uugcuuUuauc | 0.40 | 12 | 1 | 1.0000 | 0.83 | 0.65 | True |
| RBM24.1 | guugugugcag | 0.47 | 7 | 2 | 1.0000 | 1.00 | 0.86 | True |
| RBM24.2 | cggacugcgug | 0.56 | 5 | -5 | 0.0084 | 0.00 | 0.98 | False |
| RBM25.0 | gAUcgcgggca | 0.61 | 7 | 0 | 0.0587 | 1.00 | 0.36 | True |
| RBM25.1 | cGAUCGucggg | 0.58 | 11 | 0 | 0.0525 | 1.00 | 0.28 | True |
| RBM25.2 | caggggcaagc | 0.51 | 14 | -2 | 0.0546 | 0.86 | 0.51 | True |
| RBM25.3 | acugggggggc | 0.60 | 11 | 1 | 0.1145 | 0.91 | 0.02 | True |

| | | | | | | | | |
|---|---|---|---|---|---|---|---|---|
| RBM4.0 | uugcgugcgca | 0.51 | 5 | 1 | 0.0931 | 1.00 | 0.67 | True |
| RBM4.1 | augcuacguau | 0.46 | 20 | -1 | 0.0851 | 0.60 | 0.30 | True |
| RBM4.2 | uuugCguaaau | 0.37 | 18 | 1 | 0.0719 | 0.56 | 0.44 | True |
| RBM4.3 | gcgcggcgcac | 0.57 | 17 | -3 | 0.0996 | 0.82 | 0.14 | True |
| RBM4.4 | cuugcgcguaa | 0.48 | 20 | -4 | 0.0614 | 0.75 | 0.23 | True |
| RBM41.0 | ucuuccggacc | 0.61 | 9 | -4 | 0.3399 | 0.56 | 0.12 | True |
| RBM41.1 | ccuUaCuUuac | 0.39 | 7 | 2 | 0.1840 | 1.00 | 0.14 | True |
| RBM41.2 | cggcccuaCau | 0.62 | 7 | 5 | 0.2212 | 0.86 | 0.07 | True |
| RBM41.3 | agccauuaCac | 0.53 | 8 | 4 | 0.1914 | 0.88 | 0.18 | True |
| RBM41.4 | caCuuuuaCuc | 0.37 | 12 | -1 | 0.1107 | 0.92 | 0.11 | True |
| RBM45.0 | cgACuuaCgca | 0.46 | 18 | -5 | 0.2769 | 0.72 | 0.22 | True |
| RBM45.1 | uaCacgaCgcc | 0.54 | 17 | -5 | 0.3028 | 0.71 | 0.10 | True |
| RBM45.2 | cacgACGcaaa | 0.52 | 10 | -5 | 0.3373 | 0.50 | 0.28 | False |
| RBM45.3 | GaacgaCgaca | 0.53 | 20 | 0 | 0.2876 | 0.55 | 0.13 | True |
| RBM45.4 | uaCaACauaac | 0.36 | 18 | -5 | 0.4070 | 0.72 | 0.13 | True |
| RBM47.0 | aauaaucaaaa | 0.32 | 10 | 0 | 0.4909 | 1.00 | 0.97 | False |
| RBM47.1 | aauuaaAUuua | 0.19 | 11 | 1 | 1.0000 | 0.91 | 0.86 | True |
| RBM47.2 | cuAaUuUaaac | 0.26 | 11 | 2 | 1.0000 | 0.82 | 0.86 | True |
| RBM47.3 | aAUuauuaAcc | 0.32 | 11 | 2 | 1.0000 | 0.82 | 0.93 | False |
| RBM4B.0 | uCGgugucggc | 0.60 | 6 | 0 | 0.1471 | 1.00 | 0.24 | True |
| RBM4B.1 | accgCGguaaa | 0.49 | 5 | 1 | 0.2652 | 1.00 | 0.26 | True |
| RBM4B.2 | auCgggacggg | 0.58 | 8 | 1 | 0.1453 | 1.00 | 0.29 | True |
| RBM4B.3 | accgcGcgggg | 0.61 | 6 | 2 | 0.2580 | 1.00 | 0.13 | True |
| RBM6.0 | Gcgccuccgga | 0.71 | 6 | 4 | 0.0931 | 1.00 | 0.49 | True |
| RBM6.1 | uccacuccgug | 0.53 | 6 | 2 | 0.0910 | 1.00 | 0.56 | True |
| RBM6.2 | acauaccggCu | 0.49 | 20 | 0 | 0.1095 | 0.55 | 0.45 | True |
| RBM6.3 | agccuauccac | 0.51 | 15 | -5 | 0.0836 | 0.67 | 0.23 | True |
| RBM6.4 | cacguccccuu | 0.51 | 8 | -5 | 0.0642 | 0.38 | 0.58 | False |
| RBMS2.0 | cauuaUAacaa | 0.37 | 8 | 1 | 0.2531 | 1.00 | 0.13 | True |
| RBMS2.1 | gcaUAaucAag | 0.39 | 9 | -1 | 0.3130 | 0.89 | 0.16 | True |
| RBMS2.2 | gcuauaUauag | 0.42 | 10 | 1 | 0.3356 | 1.00 | 0.04 | True |
| RBMS2.3 | cauuauUAauc | 0.37 | 13 | -1 | 0.2230 | 0.92 | 0.10 | True |

| | | | | | | | | |
|---|---|---|---|---|---|---|---|---|
| RBMS3.0 | ucauauUAaua | 0.31 | 11 | -1 | 0.2332 | 0.91 | 0.07 | True |
| RBMS3.1 | guuuAuuUAau | 0.34 | 12 | -1 | 0.2859 | 0.92 | 0.08 | True |
| RBMS3.2 | caauaUAuucg | 0.38 | 9 | 1 | 0.3882 | 1.00 | 0.07 | True |
| RC3H1.0 | aUucCuaUGua | 0.37 | 5 | 4 | 0.3501 | 1.00 | 0.36 | True |
| RC3H1.1 | uccauAuugau | 0.34 | 5 | -5 | 0.0139 | 0.00 | 1.06 | False |
| RC3H1.2 | auuuauUaaug | 0.19 | 7 | 9 | 0.0533 | 0.29 | 0.37 | False |
| RC3H1.3 | auauuauauua | 0.36 | 13 | 3 | 0.0517 | 0.62 | 0.27 | True |
| RC3H1.4 | gcuguAUcagg | 0.50 | 5 | -5 | 0.0000 | 0.00 | 0.93 | False |
| SF1.0 | cUAAcuuaAcg | 0.30 | 14 | -2 | 0.2634 | 0.86 | 0.05 | True |
| SF1.1 | caauUaAccac | 0.37 | 15 | -5 | 0.2026 | 0.67 | 0.10 | True |
| SF1.2 | gaacuuUAAug | 0.33 | 17 | -5 | 0.2373 | 0.71 | 0.06 | True |
| SF1.3 | acuUAAUacuu | 0.25 | 17 | -4 | 0.2431 | 0.76 | 0.07 | True |
| SFPQ.0 | uauAgugugua | 0.38 | 12 | -5 | 0.2905 | 0.58 | 0.24 | True |
| SFPQ.1 | uacggggccgg | 0.65 | 9 | -5 | 0.2412 | 0.44 | 0.73 | False |
| SFPQ.2 | auuuguaaggu | 0.35 | 14 | -5 | 0.2960 | 0.64 | 0.30 | True |
| SFPQ.3 | cguagugguGc | 0.51 | 11 | -5 | 0.3100 | 0.55 | 0.12 | True |
| SFPQ.4 | guagcccgggu | 0.62 | 14 | -4 | 0.2447 | 0.71 | 0.03 | True |
| SNRPA.0 | gacauugcacg | 0.49 | 8 | 3 | 0.0917 | 1.00 | 0.18 | True |
| SNRPA.1 | caugcACaacc | 0.48 | 8 | 0 | 0.1019 | 1.00 | 0.08 | True |
| SNRPA.2 | ucacacgcgcc | 0.57 | 14 | -5 | 0.0644 | 0.64 | 0.46 | True |
| SRSF10.0 | aGcgugcaguu | 0.53 | 5 | 9 | 0.0488 | 0.40 | 5.54 | False |
| SRSF10.1 | gaagCagcaua | 0.40 | 5 | 11 | 0.0305 | 0.00 | 2.77 | False |
| SRSF11.0 | aagcagagag | 0.50 | 16 | -5 | 0.2069 | 0.69 | 0.84 | True |
| SRSF2.0 | cgcgcaguaaa | 0.43 | 10 | -3 | 0.0874 | 0.70 | 0.22 | True |
| SRSF2.1 | agcagcaGaaa | 0.50 | 20 | -5 | 0.0664 | 0.75 | 0.46 | True |
| SRSF2.2 | guaucCagcaa | 0.47 | 19 | -5 | 0.1074 | 0.74 | 0.20 | True |
| SRSF4.0 | gauccagggg | 0.54 | 14 | 0 | 0.1908 | 0.71 | 1.72 | False |
| SRSF5.0 | gaGcggggccA | 0.58 | 5 | 3 | 0.0815 | 1.00 | 0.32 | True |
| SRSF5.1 | ccgccgccccu | 0.57 | 13 | -2 | 0.1007 | 0.85 | 0.37 | True |
| SRSF8.0 | aGccuugacag | 0.53 | 5 | 3 | 0.0524 | 1.00 | 0.63 | True |
| SRSF8.1 | gcacccgagaa | 0.58 | 9 | 1 | 0.0990 | 1.00 | 0.49 | True |
| SRSF8.2 | ugcacccuggg | 0.60 | 8 | 2 | 0.0996 | 1.00 | 0.26 | True |

| | | | | | | | | |
|---|---|---|---|---|---|---|---|---|
| SRSF8.3 | gggacaccugc | 0.61 | 11 | -2 | 0.0966 | 0.82 | 0.40 | True |
| SRSF8.4 | uaugcagcagc | 0.46 | 20 | 1 | 0.1163 | 0.50 | 0.36 | False |
| SRSF9.0 | aaagGaagaaa | 0.44 | 5 | 1 | 0.0762 | 1.00 | 1.31 | False |
| SRSF9.1 | ccaggagacgc | 0.61 | 15 | -3 | 0.1419 | 0.80 | 0.07 | True |
| TAF15.0 | GAUCGucggg | 0.54 | 8 | -4 | 0.7630 | 0.50 | 0.90 | False |
| TAF15.1 | ggggggguaa | 0.54 | 5 | -5 | 1.0000 | 0.00 | 0.85 | False |
| TAF15.2 | gAucgcggaa | 0.61 | 9 | -4 | 1.0000 | 0.56 | 0.55 | True |
| TARDBP.0 | auaUGaGuaua | 0.31 | 5 | -5 | 0.4130 | 0.00 | 0.72 | False |
| TARDBP.1 | uauGuauGagu | 0.39 | 13 | -3 | 0.2245 | 0.77 | 0.14 | True |
| TARDBP.2 | gguaaaGUaUg | 0.39 | 5 | -5 | 0.4028 | 0.00 | 0.64 | False |
| TIA1.0 | uuuaaaaaaaa | 0.25 | 5 | -3 | 0.0030 | 0.40 | 0.16 | False |
| TIA1.1 | aauuuuuuuua | 0.28 | 20 | -5 | 0.2298 | 0.75 | 0.03 | True |
| TIA1.2 | uaaAAUuuuaa | 0.09 | 5 | 11 | 0.0016 | 0.00 | 0.10 | False |
| TIA1.3 | cuUuuaauuuu | 0.17 | 20 | -2 | 0.3240 | 0.65 | 0.28 | True |
| TIA1.4 | uuUAUuuaaaa | 0.07 | 5 | 11 | 0.2901 | 0.00 | 0.28 | False |
| TRA2A.0 | ucgguGAAgac | 0.50 | 8 | 7 | 0.1551 | 0.50 | 0.16 | False |
| TRA2A.1 | aaaGAAgaaaa | 0.39 | 16 | -2 | 0.0384 | 0.81 | 0.92 | False |
| TRA2A.2 | aaaGaaaauaa | 0.31 | 5 | -5 | 0.0163 | 0.00 | 1.03 | False |
| TRA2A.3 | acucccgaaga | 0.50 | 11 | 1 | 0.1105 | 0.91 | 0.25 | True |
| TRA2A.4 | aaaaauGaAga | 0.39 | 6 | 6 | 0.1111 | 0.83 | 0.38 | True |
| TRNAU1AP.0 | gCauuuuuUuu | 0.37 | 13 | 0 | 0.1171 | 0.85 | 0.19 | True |
| TRNAU1AP.1 | auuuuaauUuu | 0.20 | 17 | -4 | 0.1008 | 0.76 | 0.12 | True |
| TRNAU1AP.2 | aaUuauuauua | 0.16 | 18 | -5 | 0.0903 | 0.72 | 0.22 | True |
| TRNAU1AP.3 | uuuuacauuuu | 0.19 | 15 | -3 | 0.1248 | 0.80 | 0.11 | True |
| TRNAU1AP.4 | uaaaauUuaca | 0.28 | 5 | -5 | 0.0208 | 0.00 | 0.92 | False |
| UNK.0 | ccccauAgcag | 0.52 | 8 | 7 | 0.1337 | 0.50 | 0.20 | False |
| UNK.1 | aauUAguuaaa | 0.34 | 20 | -5 | 0.0783 | 0.75 | 0.25 | True |
| UNK.2 | ccaauUAgagg | 0.44 | 5 | 7 | 0.1021 | 0.80 | 0.12 | True |
| UNK.3 | caaauauAguc | 0.35 | 5 | 8 | 0.1312 | 0.60 | 0.19 | True |

| | | | | | | | | |
|---|---|---|---|---|---|---|---|---|
| ZCRB1.0 | aauaauuuuaa | 0.16 | 5 | 5 | 0.0864 | 1.00 | 0.31 | True |
| ZCRB1.1 | cgaauUAAuuu | 0.39 | 14 | -5 | 0.0468 | 0.64 | 0.19 | True |
| ZCRB1.2 | gccgcauuaau | 0.52 | 16 | -4 | 0.0845 | 0.75 | 0.36 | True |
| ZFP36.0 | uuUAUUUAuuu | 0.21 | 5 | -4 | 0.0688 | 0.20 | 0.25 | False |
| ZFP36.1 | uUAAcuuAuuu | 0.24 | 14 | -3 | 0.0965 | 0.79 | 0.20 | True |
| ZFP36.2 | uuUaaUAuuau | 0.17 | 5 | -5 | 0.1247 | 0.00 | 0.22 | False |

| **Supplementary Table S4.** eCLIP fit results | | | | | | | | | | | | |
|---|---|---|---|---|---|---|---|---|---|---|---|---|
| **RBP** | name of RBP | | | | | | | | | | | |
| **cell** | cell line from which eCLIP data was generated | | | | | | | | | | | |
| **chain_type:** | parts of the transcript which were analyzed. SPLICED=longest mature mRNA isoform in GENCODE28. INTRONS=all introns of longest isoform | | | | | | | | | | | |
| **rel_LL:** | difference between LL_fit and LL_const. Reflects how much better the model fits the data, compared to a constant, average value | | | | | | | | | | | |
| **LL_const** | likelihood that clip enrichment is a constant, independent of predicted affinity | | | | | | | | | | | |
| **LL_fit** | likelihood that clip enrichment fits the RBPamp affinity-derived prediction with optimized values for free RBP, [optionally hill coefficient], and replicate-specific clip background | | | | | | | | | | | |
| **P_const** | likelihood ratio derived probability that data are better explained by a constant | | | | | | | | | | | |
| **P_hill_one** | likelihood ratio-derived probability that data are better explained by a model with $h^{model}$ fixed at 1. Only if P($h^{model}$ = 1) < 1e-5 is the model with free $h^{model}$ selected. | | | | | | | | | | | |
| **hill_lo** | $5^{th}$ percentile of $h^{model}$ values over 1,000 bootstraps | | | | | | | | | | | |
| **hill_med** | median of $h^{model}$ values over 1,000 bootstraps | | | | | | | | | | | |
| **hill_hi** | $95^{th}$ percentile of $h^{model}$ values over 1,000 bootstraps | | | | | | | | | | | |
| **rbp_free_lo** | 5th percentile of $h^{model}$ values over 1,000 bootstraps | | | | | | | | | | | |
| **rbp_free_med** | median of $F^{model}$ over 1,000 bootstraps | | | | | | | | | | | |
| **rbp_free_hi** | 95th percentile of $F^{model}$ over 1,000 bootstraps | | | | | | | | | | | |
| **RBP** | **cell** | **chain_type** | **rel_LL** | **LL_const** | **LL_fit** | **P_const** | **P_h=1** | **$h^{model}$_lo** | **$h^{model}$_med** | **$h^{model}$_hi** | **$F^{model}$_lo** | **$F^{model}$_med** | **$F^{model}$_hi** |
| TRA2A | hepG2 | CDS | 6,622.68 | -6,571.97 | 50.7055 | 0.0000 | 0.0000 | 1.48 | 1.61 | 1.76 | 0.00 | 0.00 | 0.45 |
| TRA2A | k562 | CDS | 3,990.19 | -3,902.44 | 87.7432 | 0.0000 | 0.0000 | 1.32 | 1.52 | 1.75 | 0.00 | 0.00 | 0.00 |
| TRA2A | hepG2 | SPLICED | 3,225.68 | -3,128.58 | 97.1081 | 0.0000 | 0.0000 | 1.74 | 2.30 | 3.99 | 0.00 | 0.42 | 3.01 |
| TRA2A | hep | INTRO | 2,121. | - | 6.33 | 0.00 | 0.0 | 1.96 | 2.31 | 3.4 | 0.0 | 0.46 | 2.73 |

| | | | | | | | | | | | | | |
|---|---|---|---|---|---|---|---|---|---|---|---|---|---|
| | G2 | NS | 22 | 2,114.89 | 873 | 00 | 000 | | | 5 | 0 | | |
| TRA2A | k562 | SPLICED | 1,858.10 | -1,756.63 | 101.465 | 0.0000 | 0.0000 | 1.57 | 2.09 | 2.79 | 0.00 | 0.00 | 2.48 |
| TRA2A | hepG2 | UTR3 | 1,457.71 | -1,396.37 | 61.347 | 0.0000 | 0.0000 | 1.64 | 1.95 | 2.36 | 0.00 | 0.00 | 0.43 |
| TRA2A | k562 | UTR3 | 1,066.74 | -969.56 | 97.1805 | 0.0000 | 0.0000 | 1.59 | 1.84 | 2.27 | 0.00 | 0.00 | 3.94 |
| TRA2A | k562 | INTRONS | 349.45 | -259.54 | 89.9166 | 0.0000 | 0.0000 | 1.52 | 7.48 | 10.00 | 0.00 | 5.08 | 6.80 |
| TRA2A | hepG2 | UTR5 | 138.70 | -76.64 | 62.0665 | 0.0000 | 0.0074 | 1.00 | 1.00 | 1.00 | 0.00 | 0.00 | 2.47 |
| TRA2A | k562 | UTR5 | 39.69 | 29.39 | 69.0852 | 0.0000 | 1.0000 | 1.00 | 1.00 | 1.00 | 0.00 | 0.87 | 294.15 |
| TIA1 | hepG2 | UTR3 | 81,287.60 | -81,146.16 | 141.439 | 0.0000 | 0.0000 | 1.30 | 1.49 | 1.64 | 1.95 | 3.26 | 4.53 |
| TIA1 | hepG2 | SPLICED | 34,274.46 | -34,138.28 | 136.173 | 0.0000 | 1.0000 | 1.00 | 1.00 | 1.00 | 0.00 | 0.05 | 0.29 |
| TIA1 | k562 | UTR3 | 7,827.73 | -7,691.21 | 136.517 | 0.0000 | 1.0000 | 1.00 | 1.00 | 1.00 | 2.03 | 2.87 | 3.80 |
| TIA1 | hepG2 | INTRONS | 7,501.10 | -7,618.78 | -117.677 | 0.0000 | 0.0000 | 2.09 | 2.49 | 2.95 | 10.34 | 11.65 | 12.92 |
| TIA1 | k562 | SPLICED | 5,379.40 | -5,240.80 | 138.607 | 0.0000 | 1.0000 | 1.00 | 1.00 | 1.00 | 0.76 | 1.31 | 2.43 |
| TIA1 | hepG2 | CDS | 4,894.62 | -4,783.63 | 110.984 | 0.0000 | 0.0777 | 1.00 | 1.00 | 1.00 | 0.56 | 2.15 | 4.28 |
| TIA1 | hepG2 | UTR5 | 2,755.07 | -2,663.12 | 91.9476 | 0.0000 | 1.0000 | 1.00 | 1.00 | 1.00 | 1.55 | 2.62 | 4.04 |
| TIA1 | k562 | CDS | 1,544.21 | -1,425.86 | 118.349 | 0.0000 | 0.0693 | 1.00 | 1.00 | 1.00 | 2.61 | 5.14 | 8.75 |
| TIA1 | k562 | INTRONS | 478.14 | -343.80 | 134.341 | 0.0000 | 0.0000 | 0.53 | 3.24 | 4.48 | 0.49 | 7.10 | 8.90 |

| | | | | | | | | | | | | |
|---|---|---|---|---|---|---|---|---|---|---|---|---|
| TIA1 | k562 | UTR5 | 358.66 | -272.21 | 86.4474 | 0.0000 | 0.0014 | 1.00 | 1.00 | 1.00 | 0.00 | 2.68 | 25.48 |
| TARDBP | k562 | INTRONS | 51,052.87 | -51,086.57 | -33.7027 | 0.0000 | 0.0000 | 2.55 | 2.78 | 3.01 | 5.35 | 5.89 | 6.36 |
| TARDBP | k562 | UTR3 | 17,619.53 | -17,548.42 | 71.1065 | 0.0000 | 0.0000 | 1.12 | 1.66 | 2.42 | 0.44 | 3.17 | 5.59 |
| TARDBP | k562 | SPLICED | 6,355.75 | -6,243.49 | 112.257 | 0.0000 | 0.0000 | 1.92 | 2.77 | 5.19 | 4.19 | 7.17 | 9.49 |
| TARDBP | k562 | CDS | 458.84 | -395.82 | 63.0217 | 0.0000 | 1.0000 | 1.00 | 1.00 | 1.00 | 24.65 | 41.46 | 76.85 |
| TARDBP | k562 | UTR5 | 456.99 | -396.59 | 60.3972 | 0.0000 | 0.0000 | 2.21 | 3.23 | 4.18 | 0.00 | 0.43 | 0.45 |
| SRSF9 | hepG2 | INTRONS | 1,306.90 | -2,129.80 | -822.904 | 0.0000 | 0.0000 | 2.51 | 3.62 | 4.65 | 0.22 | 0.51 | 0.73 |
| SRSF9 | hepG2 | CDS | 559.27 | -517.73 | 41.5313 | 0.0000 | 1.0000 | 1.00 | 1.00 | 1.00 | 0.00 | 0.04 | 0.32 |
| SRSF9 | hepG2 | SPLICED | 534.16 | -480.72 | 53.4437 | 0.0000 | 0.0000 | 1.00 | 1.00 | 1.00 | 0.00 | 0.00 | 0.00 |
| SRSF9 | hepG2 | UTR5 | 261.57 | -184.58 | 76.998 | 0.0000 | 0.0867 | 1.00 | 1.00 | 1.00 | 0.00 | 0.00 | 0.15 |
| SRSF9 | hepG2 | UTR3 | 0.00 | 21.39 | 21.3883 | 1.0000 | 0.4432 | 1.00 | 1.00 | 1.00 | 0.00 | 0.50 | 18.04 |
| SFPQ | hepG2 | INTRONS | 2,211.81 | -3,661.05 | -1449.25 | 0.0000 | 0.0000 | 1.66 | 6.32 | 10.00 | 0.00 | 3.17 | 3.78 |
| SFPQ | hepG2 | CDS | 494.18 | -542.17 | -47.9962 | 0.0000 | 0.0000 | 2.70 | 3.88 | 5.29 | 1.17 | 1.97 | 2.36 |
| SFPQ | hepG2 | UTR5 | 120.34 | -49.49 | 70.8462 | 0.0000 | 1.0000 | 1.00 | 1.00 | 1.00 | 0.00 | 0.00 | 0.46 |
| SFPQ | hepG2 | SPLICED | 75.51 | -127.37 | -51.8608 | 0.0000 | 1.0000 | 1.00 | 1.00 | 1.00 | 0.00 | 0.00 | 21.53 |
| SFPQ | hepG2 | UTR3 | 0.00 | -478.34 | -478.343 | 1.0000 | 1.0000 | 1.00 | 1.00 | 1.00 | 10.58 | 11.06 | 11.38 |
| RBFOX2 | hep | INTRO | 8,282. | -8,148. | 133. | 0.00 | 0.0 | 1.00 | 1.00 | 1.0 | 0.0 | 0.00 | 0.43 |

| | | | | | | | | | | | | | |
|---|---|---|---|---|---|---|---|---|---|---|---|---|---|
| | G2 | NS | 10 | 97 | 129 | 00 | 171 | | | 0 | 0 | | |
| RBFOX2 | k562 | INTRONS | 3,115.44 | -3,146.14 | -30.704 | 0.0000 | 0.0014 | 1.00 | 1.00 | 1.00 | 0.00 | 0.00 | 0.00 |
| RBFOX2 | hepG2 | UTR3 | 2,180.24 | -2,064.86 | 115.377 | 0.0000 | 1.0000 | 1.00 | 1.00 | 1.00 | 1.35 | 2.78 | 5.00 |
| RBFOX2 | k562 | UTR3 | 1,539.37 | -1,402.65 | 136.722 | 0.0000 | 1.0000 | 1.00 | 1.00 | 1.00 | 0.66 | 1.79 | 3.24 |
| RBFOX2 | hepG2 | SPLICED | 949.80 | -861.60 | 88.1984 | 0.0000 | 1.0000 | 1.00 | 1.00 | 1.00 | 0.00 | 0.00 | 7.46 |
| RBFOX2 | k562 | SPLICED | 343.03 | -235.69 | 107.341 | 0.0000 | 1.0000 | 1.00 | 1.00 | 1.00 | 0.00 | 0.00 | 295.68 |
| RBFOX2 | hepG2 | CDS | 192.61 | -80.52 | 112.094 | 0.0000 | 0.2320 | 1.00 | 1.00 | 1.00 | 0.00 | 3.18 | 22.00 |
| RBFOX2 | hepG2 | UTR5 | 109.95 | -47.88 | 62.0691 | 0.0000 | 1.0000 | 1.00 | 1.00 | 1.00 | 0.35 | 6.69 | 19.48 |
| RBFOX2 | k562 | CDS | 108.49 | 27.61 | 136.109 | 0.0000 | 1.0000 | 1.00 | 1.00 | 1.00 | 0.00 | 2.92 | 9.64 |
| RBFOX2 | k562 | UTR5 | 55.49 | 10.59 | 66.0842 | 0.0000 | 0.1639 | 1.00 | 1.00 | 1.00 | 0.00 | 0.00 | 0.89 |
| PCBP2 | hepG2 | UTR3 | 5,329.33 | -5,255.22 | 74.1142 | 0.0000 | 0.0000 | 1.41 | 1.68 | 2.02 | 125.49 | 168.56 | 208.69 |
| PCBP2 | hepG2 | INTRONS | 4,828.04 | -4,746.97 | 81.0636 | 0.0000 | 0.0003 | 1.00 | 1.00 | 1.00 | 104.19 | 129.44 | 155.45 |
| PCBP2 | hepG2 | CDS | 1,064.33 | -984.97 | 79.3665 | 0.0000 | 0.0089 | 1.00 | 1.00 | 1.00 | 94.51 | 126.00 | 160.43 |
| PCBP2 | hepG2 | SPLICED | 1,032.26 | -966.61 | 65.647 | 0.0000 | 0.0000 | 0.92 | 1.65 | 2.43 | 36.99 | 129.65 | 205.23 |
| PCBP2 | hepG2 | UTR5 | 767.98 | -721.66 | 46.3214 | 0.0000 | 1.0000 | 1.00 | 1.00 | 1.00 | 19.24 | 47.13 | 92.82 |
| PCBP1 | k562 | UTR3 | 300.51 | -220.01 | 80.5035 | 0.0000 | 0.0000 | 2.11 | 2.66 | 4.08 | 480.86 | 614.35 | 712.74 |
| PCBP1 | hepG2 | CDS | 198.03 | -95.72 | 102.305 | 0.0000 | 0.3982 | 1.00 | 1.00 | 1.00 | 539.10 | 805.09 | 1000.00 |
| PCBP1 | k562 | INTRONS | 173.33 | -70.89 | 102.438 | 0.0000 | 0.0000 | 0.15 | 2.50 | 3.81 | 0.50 | 700.27 | 798.90 |

| | | | | | | | | | | | | |
|---|---|---|---|---|---|---|---|---|---|---|---|---|
| PCBP1 | hepG2 | INTRONS | 153.70 | -103.49 | 50.2109 | 0.0000 | 0.0000 | 0.25 | 2.80 | 5.73 | 0.00 | 0.29 | 629.62 |
| PCBP1 | hepG2 | UTR3 | 105.54 | -23.94 | 81.601 | 0.0000 | 0.0000 | 1.53 | 3.88 | 7.16 | 422.16 | 633.81 | 796.80 |
| PCBP1 | k562 | SPLICED | 76.03 | 20.61 | 96.6376 | 0.0000 | 0.0000 | 2.40 | 8.94 | 10.00 | 344.88 | 470.55 | 710.15 |
| PCBP1 | k562 | CDS | 74.95 | 39.32 | 114.273 | 0.0000 | 0.0000 | 1.83 | 6.50 | 10.00 | 351.18 | 445.79 | 972.89 |
| PCBP1 | hepG2 | UTR5 | 57.68 | -32.28 | 25.4001 | 0.0000 | 0.3191 | 1.00 | 1.00 | 1.00 | 0.00 | 170.44 | 884.44 |
| PCBP1 | hepG2 | SPLICED | 56.69 | 18.93 | 75.6208 | 0.0000 | 0.0000 | 0.25 | 10.00 | 10.00 | 0.00 | 367.56 | 466.61 |
| PCBP1 | k562 | UTR5 | 17.03 | 64.03 | 81.0605 | 0.0000 | 0.0060 | 1.00 | 1.00 | 1.00 | 7.88 | 1000.00 | 1000.00 |
| KHSRP | k562 | INTRONS | 3,024.69 | -2,964.56 | 60.1264 | 0.0000 | 0.0000 | 3.83 | 4.88 | 5.77 | 2.47 | 2.77 | 3.01 |
| KHSRP | k562 | UTR3 | 1,109.36 | -1,207.95 | -98.5861 | 0.0000 | 0.0000 | 5.02 | 6.45 | 8.42 | 2.49 | 2.80 | 3.17 |
| KHSRP | k562 | SPLICED | 1,047.76 | -1,213.70 | -165.936 | 0.0000 | 0.0000 | 6.90 | 10.00 | 10.00 | 2.82 | 3.21 | 3.33 |
| KHSRP | k562 | CDS | 871.72 | -1,171.78 | -300.061 | 0.0000 | 0.0000 | 3.36 | 4.96 | 6.73 | 0.20 | 2.25 | 2.97 |
| KHSRP | k562 | UTR5 | 708.85 | -631.82 | 77.0383 | 0.0000 | 0.0000 | 2.60 | 3.95 | 5.95 | 1.97 | 3.12 | 3.68 |
| KHSRP | hepG2 | INTRONS | 655.79 | -678.05 | -22.258 | 0.0000 | 0.0000 | 1.33 | 4.50 | 7.94 | 0.00 | 2.28 | 2.60 |
| KHSRP | hepG2 | SPLICED | 434.85 | -423.66 | 11.1972 | 0.0000 | 0.0000 | 10.00 | 10.00 | 10.00 | 3.31 | 3.44 | 3.56 |
| KHSRP | hepG2 | UTR3 | 193.30 | -130.61 | 62.6865 | 0.0000 | 0.0000 | 1.99 | 2.94 | 5.01 | 2.04 | 2.81 | 3.72 |
| KHSRP | hepG2 | UTR5 | 80.43 | -18.38 | 62.0524 | 0.0000 | 0.0008 | 1.00 | 1.00 | 1.00 | 0.00 | 0.00 | 0.96 |
| KHSRP | hepG2 | CDS | 2.22 | -31.68 | -29.4607 | 0.3504 | 1.0000 | 1.00 | 1.00 | 1.00 | 0.00 | 4.27 | 12.20 |
| IGF2BP2 | k56 | UTR3 | 68.76 | 9.76 | 78.5 | 0.00 | 0.0 | 1.00 | 1.00 | 1.0 | 7.9 | 16.0 | 50.2 |

| | | | | | | | | | | | | |
|---|---|---|---|---|---|---|---|---|---|---|---|---|
| | 2 | | | | 194 | 00 | 143 | | | 0 | 4 | 2 | 5 |
| IGF2BP2 | k562 | SPLICED | 41.01 | 63.76 | 104.778 | 0.0000 | 0.3102 | 1.00 | 1.00 | 1.00 | 0.30 | 3.61 | 10.30 |
| IGF2BP2 | k562 | UTR5 | 27.25 | 43.19 | 70.4355 | 0.0000 | 0.0232 | 1.00 | 1.00 | 1.00 | 0.00 | 0.13 | 5.49 |
| IGF2BP2 | k562 | CDS | 11.98 | 51.72 | 63.6986 | 0.0001 | 1.0000 | 1.00 | 1.00 | 1.00 | 0.48 | 6.10 | 20.45 |
| IGF2BP2 | k562 | INTRONS | 0.00 | -190.27 | -190.267 | 1.0000 | 0.9987 | 1.00 | 1.00 | 1.00 | 32.42 | 32.94 | 33.43 |
| IGF2BP1 | hepG2 | CDS | 411.34 | -304.37 | 106.966 | 0.0000 | 0.0000 | 1.74 | 2.49 | 3.43 | 7.60 | 9.98 | 12.62 |
| IGF2BP1 | hepG2 | UTR3 | 68.72 | -22.82 | 45.8971 | 0.0000 | 0.0000 | 1.00 | 1.00 | 1.00 | 22.64 | 53.24 | 406.82 |
| IGF2BP1 | k562 | CDS | 26.52 | -25.51 | 1.00917 | 0.0000 | 1.0000 | 1.00 | 1.00 | 1.00 | 6.40 | 16.48 | 534.14 |
| IGF2BP1 | hepG2 | UTR5 | 8.78 | 84.72 | 93.4996 | 0.0015 | 1.0000 | 1.00 | 1.00 | 1.00 | 12.14 | 45.86 | 153.09 |
| IGF2BP1 | hepG2 | SPLICED | 6.08 | 81.95 | 88.0326 | 0.0162 | 0.0123 | 1.00 | 1.00 | 1.00 | 5.86 | 13.55 | 36.33 |
| IGF2BP1 | k562 | UTR5 | 2.45 | 25.14 | 27.5916 | 0.2969 | 1.0000 | 1.00 | 1.00 | 1.00 | 43.86 | 113.03 | 678.20 |
| IGF2BP1 | k562 | SPLICED | 0.00 | 59.52 | 59.5208 | 1.0000 | 0.9999 | 1.00 | 1.00 | 1.00 | 31.46 | 42.97 | 1000.00 |
| IGF2BP1 | k562 | UTR3 | 0.00 | -292.98 | -292.978 | 1.0000 | 0.9992 | 1.00 | 1.00 | 1.00 | 27.22 | 27.57 | 491.61 |
| IGF2BP1 | hepG2 | INTRONS | 0.00 | -1,112.05 | -1112.05 | 1.0000 | 0.9978 | 1.00 | 1.00 | 1.00 | 45.28 | 45.95 | 46.62 |
| IGF2BP1 | k562 | INTRONS | 0.00 | -2,603.89 | -2603.89 | 1.0000 | 0.9973 | 1.00 | 1.00 | 1.00 | 37.81 | 38.59 | 39.30 |
| HNRNPL | hepG2 | INTRONS | 202,029.48 | -202,273.48 | -244.005 | 0.0000 | 0.0000 | 1.48 | 1.57 | 1.66 | 0.66 | 0.81 | 0.95 |
| HNRNPL | k562 | INTRONS | 64,997.71 | -64,869.53 | 128.175 | 0.0000 | 0.0000 | 1.24 | 1.38 | 1.53 | 0.20 | 0.38 | 0.57 |
| HNRNPL | hepG2 | SPLICED | 7,113.60 | -6,996. | 116.712 | 0.0000 | 0.0000 | 1.86 | 3.55 | 5.52 | 0.00 | 1.31 | 1.66 |

| | | | | | | | | | | | | | |
|---|---|---|---|---|---|---|---|---|---|---|---|---|---|
| | | | | 89 | | | | | | | | | |
| HNRNPL | k562 | UTR3 | 5,955.83 | -5,875.12 | 80.7153 | 0.0000 | 0.0000 | 1.52 | 1.64 | 1.78 | 0.00 | 0.00 | 0.00 |
| HNRNPL | hepG2 | UTR3 | 3,827.83 | -3,733.54 | 94.2855 | 0.0000 | 0.0000 | 2.27 | 2.83 | 3.39 | 0.00 | 0.00 | 0.11 |
| HNRNPL | k562 | SPLICED | 3,676.32 | -3,563.01 | 113.31 | 0.0000 | 0.0000 | 1.40 | 1.94 | 3.28 | 0.00 | 0.83 | 1.63 |
| HNRNPL | k562 | CDS | 826.80 | -743.75 | 83.0512 | 0.0000 | 0.0000 | 1.99 | 2.45 | 10.00 | 0.00 | 2.09 | 7.17 |
| HNRNPL | k562 | UTR5 | 362.53 | -273.78 | 88.7528 | 0.0000 | 0.0000 | 1.99 | 2.62 | 4.17 | 0.36 | 0.42 | 6.79 |
| HNRNPL | hepG2 | UTR5 | 331.22 | -307.71 | 23.5048 | 0.0000 | 0.0000 | 1.03 | 3.66 | 6.32 | 0.00 | 0.37 | 0.43 |
| HNRNPL | hepG2 | CDS | -3,716.09 | 106.55 | -3609.53 | 1.0000 | 1.0000 | 1.00 | 1.00 | 1.00 | 0.00 | 0.50 | 0.50 |
| HNRNPK | k562 | INTRONS | 7,157.96 | -7,436.93 | -278.973 | 0.0000 | 0.0000 | 3.42 | 3.86 | 4.29 | 1.38 | 1.63 | 1.82 |
| HNRNPK | hepG2 | INTRONS | 6,610.25 | -6,767.13 | -156.88 | 0.0000 | 0.0000 | 2.86 | 3.16 | 3.57 | 0.61 | 0.94 | 1.22 |
| HNRNPK | k562 | UTR3 | 4,534.27 | -4,771.79 | -237.528 | 0.0000 | 0.0000 | 3.23 | 3.76 | 4.55 | 1.46 | 1.91 | 2.39 |
| HNRNPK | k562 | SPLICED | 3,143.06 | -3,227.81 | -84.7532 | 0.0000 | 0.0000 | 2.78 | 3.17 | 3.68 | 0.93 | 1.33 | 1.73 |
| HNRNPK | hepG2 | SPLICED | 1,340.98 | -1,346.59 | -5.60891 | 0.0000 | 0.0000 | 3.05 | 3.54 | 4.28 | 0.18 | 1.01 | 1.49 |
| HNRNPK | hepG2 | UTR3 | 1,069.11 | -1,078.09 | -8.98901 | 0.0000 | 0.0000 | 2.97 | 3.44 | 4.21 | 0.01 | 1.00 | 1.66 |
| HNRNPK | k562 | CDS | 618.05 | -596.44 | 21.61 | 0.0000 | 0.0000 | 3.15 | 3.85 | 5.01 | 0.86 | 1.36 | 1.82 |
| HNRNPK | k562 | UTR5 | 438.84 | -381.22 | 57.6228 | 0.0000 | 0.0000 | 1.98 | 2.67 | 3.96 | 0.00 | 1.27 | 2.36 |

| | | | | | | | | | | | | | |
|---|---|---|---|---|---|---|---|---|---|---|---|---|---|
| HNRNPK | hepG2 | CDS | 433.11 | -468.94 | -35.8248 | 0.0000 | 0.0000 | 0.01 | 4.63 | 6.79 | 0.03 | 0.79 | 1.33 |
| HNRNPK | hepG2 | UTR5 | 274.76 | -221.84 | 52.9213 | 0.0000 | 0.0000 | 3.28 | 4.69 | 6.47 | 1.68 | 2.36 | 2.82 |
| HNRNPC | hepG2 | INTRONS | 55,733.40 | -56,379.92 | -646.519 | 0.0000 | 0.0000 | 0.30 | 0.33 | 0.45 | 0.00 | 0.00 | 0.13 |
| HNRNPC | k562 | INTRONS | 18,884.44 | -18,853.04 | 31.3971 | 0.0000 | 0.0000 | 0.57 | 0.61 | 0.70 | 0.00 | 0.01 | 0.09 |
| HNRNPC | k562 | UTR3 | 6,596.30 | -6,519.45 | 76.8487 | 0.0000 | 0.0000 | 0.50 | 0.62 | 0.75 | 0.00 | 0.00 | 0.00 |
| HNRNPC | k562 | SPLICED | 4,090.37 | -4,065.99 | 24.3882 | 0.0000 | 0.0000 | 0.50 | 0.65 | 0.81 | 0.00 | 0.00 | 0.09 |
| HNRNPC | hepG2 | UTR3 | 2,444.52 | -2,306.88 | 137.647 | 0.0000 | 0.0000 | 0.22 | 0.29 | 0.39 | 0.00 | 0.31 | 5.80 |
| HNRNPC | hepG2 | SPLICED | 1,487.13 | -1,382.13 | 105.005 | 0.0000 | 0.0000 | 0.28 | 0.34 | 0.45 | 0.00 | 0.12 | 1.46 |
| HNRNPC | k562 | CDS | 390.52 | -302.31 | 88.2047 | 0.0000 | 0.0001 | 1.00 | 1.00 | 1.00 | 13.89 | 41.17 | 91.80 |
| HNRNPC | k562 | UTR5 | 302.54 | -247.01 | 55.5231 | 0.0000 | 0.0000 | 0.26 | 0.33 | 0.51 | 0.00 | 0.02 | 12.09 |
| HNRNPC | hepG2 | UTR5 | 225.71 | -185.51 | 40.1998 | 0.0000 | 0.0000 | 0.28 | 0.42 | 10.00 | 0.00 | 0.32 | 28.56 |
| HNRNPC | hepG2 | CDS | 204.35 | -135.50 | 68.8444 | 0.0000 | 0.0526 | 1.00 | 1.00 | 1.00 | 11.61 | 71.78 | 342.84 |
| HNRNPA1 | hepG2 | INTRONS | 196.03 | -182.01 | 14.0222 | 0.0000 | 0.0000 | 2.81 | 5.91 | 10.00 | 176.64 | 198.79 | 248.86 |
| HNRNPA1 | k562 | UTR5 | 31.46 | 46.73 | 78.1862 | 0.0000 | 1.0000 | 1.00 | 1.00 | 1.00 | 0.00 | 1.89 | 55.99 |
| HNRNPA1 | hepG2 | UTR5 | 4.83 | 70.51 | 75.3422 | 0.0464 | 0.1489 | 1.00 | 1.00 | 1.00 | 5.25 | 37.65 | 573.59 |
| HNRNPA1 | k562 | SPLICED | 0.00 | 44.89 | 44.8857 | 1.0000 | 0.9996 | 1.00 | 1.00 | 1.00 | 0.00 | 17.50 | 18.30 |
| HNRNPA1 | k562 | UTR3 | 0.00 | 19.55 | 19.5542 | 1.0000 | 0.9994 | 1.00 | 1.00 | 1.00 | 0.00 | 12.07 | 12.34 |

| | | | | | | | | | | | | | |
|---|---|---|---|---|---|---|---|---|---|---|---|---|---|
| HNRNPA1 | hepG2 | SPLICED | 0.00 | 24.83 | 24.8277 | 1.0000 | 0.9994 | 1.00 | 1.00 | 1.00 | 18.47 | 19.07 | 19.67 |
| HNRNPA1 | hepG2 | UTR3 | 0.00 | -17.60 | -17.6022 | 1.0000 | 0.9991 | 1.00 | 1.00 | 1.00 | 14.21 | 14.56 | 14.77 |
| HNRNPA1 | k562 | CDS | 0.00 | -67.90 | -67.8961 | 1.0000 | 0.9989 | 1.00 | 1.00 | 1.00 | 13.19 | 13.39 | 13.58 |
| HNRNPA1 | hepG2 | CDS | 0.00 | -253.58 | -253.58 | 1.0000 | 0.9987 | 1.00 | 1.00 | 1.00 | 13.81 | 14.01 | 14.21 |
| HNRNPA1 | k562 | INTRONS | -109.42 | -41.41 | -150.825 | 1.0000 | 0.0000 | 0.05 | 10.00 | 10.00 | 0.50 | 192.25 | 261.21 |
| FUS | k562 | INTRONS | 4,319.64 | -4,596.80 | -277.166 | 0.0000 | 0.0000 | 4.37 | 5.84 | 6.96 | 0.17 | 0.21 | 0.23 |
| FUS | k562 | UTR3 | 2,026.94 | -2,315.92 | -288.983 | 0.0000 | 0.0000 | 3.94 | 4.52 | 5.48 | 0.20 | 0.23 | 0.25 |
| FUS | hepG2 | INTRONS | 884.31 | -1,531.08 | -646.771 | 0.0000 | 0.0000 | 0.08 | 10.00 | 10.00 | 0.00 | 0.23 | 0.50 |
| FUS | k562 | UTR5 | 331.34 | -250.39 | 80.9558 | 0.0000 | 0.0000 | 1.00 | 1.00 | 1.00 | 0.00 | 0.00 | 0.06 |
| FUS | k562 | SPLICED | 52.99 | -39.86 | 13.1371 | 0.0000 | 0.0000 | 0.01 | 4.47 | 6.63 | 0.22 | 0.28 | 0.62 |
| FUS | hepG2 | UTR5 | 5.58 | 27.03 | 32.6089 | 0.0249 | 0.0001 | 1.00 | 1.00 | 1.00 | 0.00 | 0.09 | 24.41 |
| FUS | hepG2 | SPLICED | 0.00 | 30.24 | 30.237 | 1.0000 | 0.9998 | 1.00 | 1.00 | 1.00 | 27.74 | 33.45 | 41.36 |
| FUS | hepG2 | UTR3 | 0.00 | -66.09 | -66.0899 | 1.0000 | 0.9996 | 1.00 | 1.00 | 1.00 | 37.20 | 39.07 | 40.64 |
| FUS | hepG2 | CDS | 0.00 | -188.65 | -188.654 | 1.0000 | 0.9995 | 1.00 | 1.00 | 1.00 | 33.03 | 34.05 | 35.05 |
| FUS | k562 | CDS | 0.00 | -165.35 | -165.348 | 1.0000 | 0.9995 | 1.00 | 1.00 | 1.00 | 35.23 | 37.51 | 328.18 |
| FUBP3 | hepG2 | UTR3 | 4,722.27 | -4,575. | 146.704 | 0.0000 | 1.0000 | 1.00 | 1.00 | 1.00 | 0.78 | 1.43 | 2.21 |

| | | | | | | | | | | | | | |
|---|---|---|---|---|---|---|---|---|---|---|---|---|---|
| | | | | 57 | | | | | | | | | |
| FUBP3 | hepG2 | SPLICED | 3,572.21 | -3,463.76 | 108.45 | 0.0000 | 0.0000 | 1.12 | 1.62 | 2.37 | 0.59 | 1.79 | 2.91 |
| FUBP3 | hepG2 | CDS | 201.17 | -85.09 | 116.077 | 0.0000 | 1.0000 | 1.00 | 1.00 | 1.00 | 2.04 | 10.65 | 26.76 |
| FUBP3 | hepG2 | UTR5 | 96.99 | -9.32 | 87.6748 | 0.0000 | 1.0000 | 1.00 | 1.00 | 1.00 | 1.99 | 6.62 | 22.19 |
| FUBP3 | hepG2 | INTRONS | 31.89 | 75.43 | 107.319 | 0.0000 | 0.0000 | 0.04 | 5.61 | 10.00 | 0.48 | 480.42 | 827.25 |
| EWSR1 | k562 | INTRONS | 11,858.78 | -11,794.42 | 64.3641 | 0.0000 | 0.0000 | 2.28 | 2.67 | 3.04 | 21.58 | 24.68 | 27.49 |
| EWSR1 | k562 | SPLICED | 926.04 | -876.79 | 49.2523 | 0.0000 | 0.0000 | 3.05 | 4.11 | 5.48 | 18.10 | 23.38 | 27.74 |
| EWSR1 | k562 | CDS | 773.48 | -705.03 | 68.4559 | 0.0000 | 0.0000 | 1.99 | 3.03 | 4.08 | 24.13 | 32.59 | 40.57 |
| EWSR1 | k562 | UTR5 | 704.96 | -630.36 | 74.5995 | 0.0000 | 1.0000 | 1.00 | 1.00 | 1.00 | 0.00 | 0.78 | 4.59 |
| EWSR1 | k562 | UTR3 | 478.48 | -541.30 | -62.8198 | 0.0000 | 0.0000 | 1.85 | 4.12 | 5.46 | 0.39 | 18.12 | 21.80 |
| EIF4G2 | k562 | INTRONS | 877.49 | -986.19 | -108.7 | 0.0000 | 0.0000 | 10.00 | 10.00 | 10.00 | 2.03 | 2.15 | 2.31 |
| EIF4G2 | k562 | UTR5 | 73.59 | 5.40 | 78.9949 | 0.0000 | 0.0000 | 3.46 | 5.45 | 10.00 | 1.73 | 2.27 | 2.76 |
| EIF4G2 | k562 | SPLICED | 70.42 | -22.94 | 47.4825 | 0.0000 | 0.0018 | 1.00 | 1.00 | 1.00 | 0.00 | 0.17 | 0.80 |
| EIF4G2 | k562 | UTR3 | 64.61 | -26.24 | 38.3745 | 0.0000 | 0.0000 | 0.01 | 5.44 | 10.00 | 0.21 | 1.42 | 2.65 |
| EIF4G2 | k562 | CDS | 32.34 | 45.05 | 77.3882 | 0.0000 | 1.0000 | 1.00 | 1.00 | 1.00 | 0.00 | 0.92 | 19.64 |
